# Integrated longitudinal multi-omics study identifies immune programs associated with COVID-19 severity and mortality in 1152 hospitalized participants

**DOI:** 10.1101/2023.11.03.565292

**Authors:** Jeremy P. Gygi, Cole Maguire, Ravi K. Patel, Pramod Shinde, Anna Konstorum, Casey P. Shannon, Leqi Xu, Annmarie Hoch, Naresh Doni Jayavelu, IMPACC Network, Elias K. Haddad, Elaine F. Reed, Monica Kraft, Grace A. McComsey, Jordan Metcalf, Al Ozonoff, Denise Esserman, Charles B. Cairns, Nadine Rouphael, Steven E. Bosinger, Seunghee Kim-Schulze, Florian Krammer, Lindsey B. Rosen, Harm van Bakel, Michael Wilson, Walter Eckalbar, Holden Maecker, Charles R. Langelier, Hanno Steen, Matthew C. Altman, Ruth R. Montgomery, Ofer Levy, Esther Melamed, Bali Pulendran, Joann Diray-Arce, Kinga K. Smolen, Gabriela K. Fragiadakis, Patrice M. Becker, Alison D. Augustine, Rafick P. Sekaly, Lauren I. R. Ehrlich, Slim Fourati, Bjoern Peters, Steven H. Kleinstein, Leying Guan

## Abstract

Hospitalized COVID-19 patients exhibit diverse clinical outcomes, with some individuals diverging over time even though their initial disease severity appears similar. A systematic evaluation of molecular and cellular profiles over the full disease course can link immune programs and their coordination with progression heterogeneity. In this study, we carried out deep immunophenotyping and conducted longitudinal multi-omics modeling integrating ten distinct assays on a total of 1,152 IMPACC participants and identified several immune cascades that were significant drivers of differential clinical outcomes. Increasing disease severity was driven by a temporal pattern that began with the early upregulation of immunosuppressive metabolites and then elevated levels of inflammatory cytokines, signatures of coagulation, NETosis, and T-cell functional dysregulation. A second immune cascade, predictive of 28-day mortality among critically ill patients, was characterized by reduced total plasma immunoglobulins and B cells, as well as dysregulated IFN responsiveness. We demonstrated that the balance disruption between IFN-stimulated genes and IFN inhibitors is a crucial biomarker of COVID-19 mortality, potentially contributing to the failure of viral clearance in patients with fatal illness. Our longitudinal multi-omics profiling study revealed novel temporal coordination across diverse omics that potentially explain disease progression, providing insights that inform the targeted development of therapies for hospitalized COVID-19 patients, especially those critically ill.

## INTRODUCTION

Coronavirus disease 2019 (COVID-19), caused by severe acute respiratory syndrome coronavirus 2 (SARS-CoV-2) infection, reflects a complex balance between viral replication, host immune response and physiological manifestations such as hypoxia, organ dysfunction, and systemic inflammation (1–4). Patients hospitalized with COVID-19 exhibit a broad range of clinical outcomes, from moderate severity and short hospital stay to critical illness with prolonged hospitalization, and even mortality (5, 6). Profiling the immune response in a clinically diverse cohort would enable the linking of molecular and cellular mechanisms with these differential outcomes and could inform the development of targeted therapies.

Several hallmarks of severe COVID-19 have been identified, including over-production of proinflammatory cytokines (7–9), lymphopenia (7, 10, 11), neutrophil extracellular traps (NETs) formation (12), impaired interferon (IFN) signaling (13–16), anti-IFN auto-antibodies (17), and immune senescence (18). While these studies have identified many individual components underlying the pathophysiology of SARS-CoV-2 infection, it is still unclear how their temporal coordination interplay contributes to the observed heterogeneity of responses among hospitalized patients, especially among the critically ill. Why do some COVID-19 patients survive while others experience fatal illnesses despite seemingly similar severity at hospital admission? Additionally, most existing studies focus on measurements from peripheral blood and leverage one, or a small number, of assays thus limiting opportunities to identify biologically relevant connections between tissues and mechanisms that operate across scales. Such a systems-level understanding requires longitudinal multi-omics studies on large-scale and clinically well-defined hospitalized cohorts to characterize innate, adaptive, and mucosal immune cascades associated with disease severity (19–22).

The Immunophenotyping Assessment in a COVID-19 Cohort (IMPACC) study aims to gain a panoramic understanding of SARS-CoV-2 infection via collection and analysis of detailed clinical, laboratory, and radiographic data along with longitudinal biologic samplings of blood and respiratory secretions on participants hospitalized with COVID-19 across the US (23). Previously, IMPACC conducted deep immunophenotyping on longitudinal samples for 539 hospitalized participants mostly enrolled before September 2020 to identify biological states associated with SARS-CoV-2 disease course trajectories (6).

In this current work, we first carried out an integrative multi-omics analysis of the existing IMPACC data to develop models that predict disease severity and mortality. To test these models, we then generated independent immunophenotyping data on an additional 613 hospitalized IMPACC participants enrolled after September 2020. Our integrative analysis revealed strong orchestrated variations across serum soluble proteins (cytokines, chemokines, and secreted receptors), plasma proteins and metabolites, gene expression from PBMC and nasal swabs, circulating immune cell frequencies, as well as viral loads, and SARS-CoV-2 serum antibodies. For example, early dysregulation of metabolism in plasma, depressed B-cell functions, and disrupted balance in interferon signaling could distinguish fatality amongst critically ill patients whose disease severity levels were similar at hospital admission. Although the dysregulation of IFN signaling in severe COVID-19 has been hypothesized previously (24, 25), our integrative analysis revealed evidence for the potentially critical role of inhibitory genes in this dysregulation through investigating the temporal coordination. In summary, we present in this work a set of robust immunological findings that are capable of distinguishing between varying severities of hospitalized COVID-19 and yield insight into the pathophysiology of severe disease. We foresee the general utility of applying our analysis strategy for mining novel coordinated and temporal dynamics to investigate other infectious diseases beyond the current pandemic.

## RESULTS

### Longitudinal multi-omics profiling of SARS-CoV-2 infection response

IMPACC enrolled 1,152 participants from 20 US sites from May 2020 to March 2021, prior to the widespread rollout of SARS-CoV-2 vaccines (PMID: design paper). All participants were assigned to one of five COVID-19 disease trajectory groups (TGs) using latent class modeling based on a modified WHO score, which is referred to as respiratory ordinal status (5) (Fig. 1B). Groups ranged from Trajectory Group 1 (TG1, length of hospital stay (LOS) around 3-5 days and with a largely uncomplicated hospital course), Trajectory Group 2 (TG2, LOS around 7-14 days and discharged with no limitations), Trajectory Group 3 (TG3, LOS around 10-14 days and discharged with limitations), Trajectory Group 4 (TG4, LOS around 28 days or more) to Trajectory Group 5 (TG5, fatal illness by day 28). Both TG4 and TG5 were considered critically ill, with TG5 uniquely containing participants who incurred fatal illness within the first 28 days following hospitalization.

**Fig. 1.**
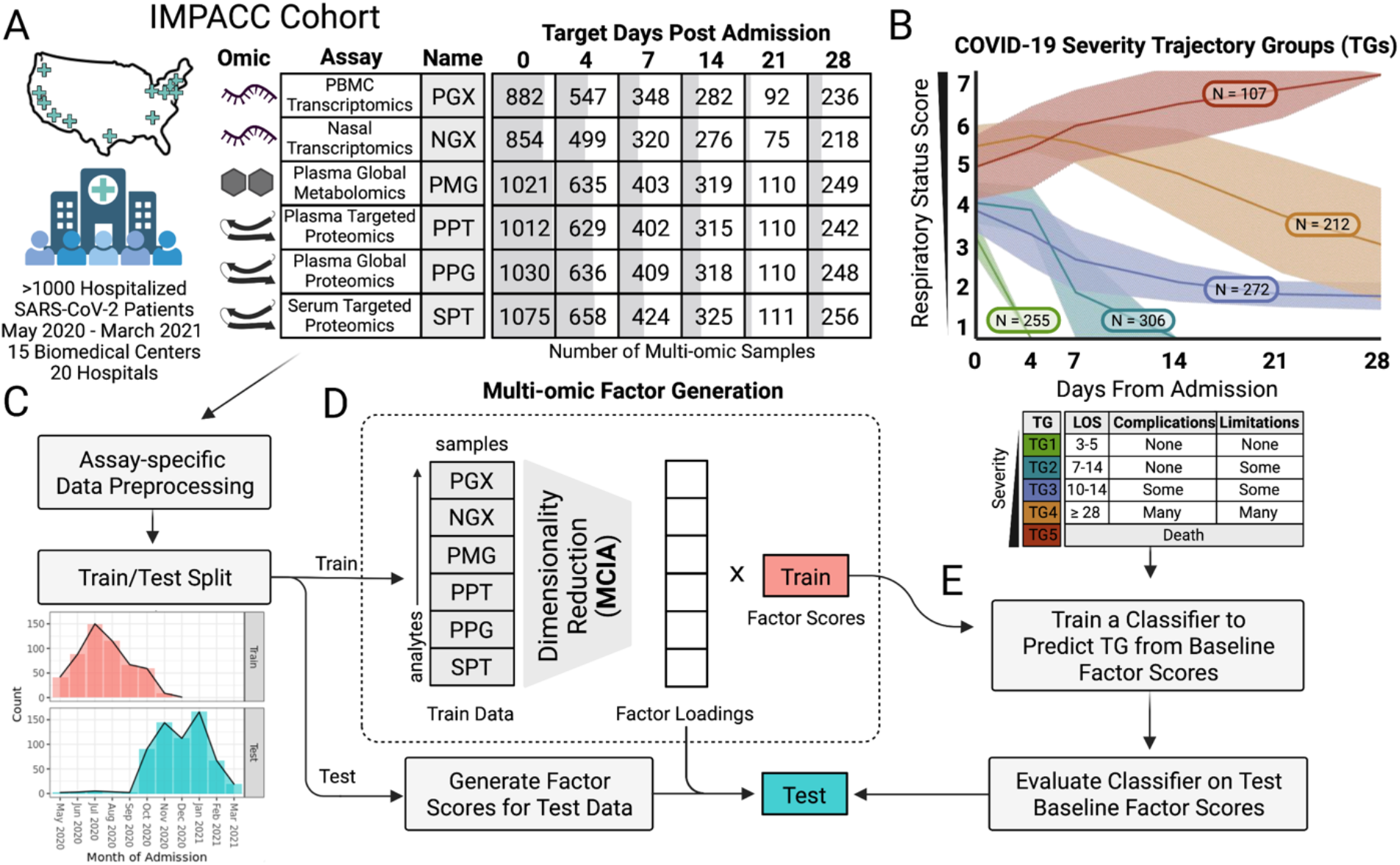
Data overview and multi-omics factor generation. **(A)** Table with the number of samples used in the integrative analysis separated by assay (rows) and scheduled time of collection (columns). Cells are shaded by the relative number of multi-omics samples available. **(B)** Plot of clinical trajectory group (TG) assignments for all IMPACC cohort participants (N=1,164) and clinical descriptions for each TG. X-axis represents days from hospital admission and y-axis represents ordinal respiratory status score (1,2=discharged, 3-6=hospitalized, 7=fatal). **(C)** Preprocessed data for different assays were split into train and test cohorts. **(D)** Dimensionality reduction was performed via MCIA on the training cohort assays to construct multi-omics factors and loadings. **(E)** Baseline factor scores were used to train a classifier for predicting TG with model selected via cross-validation. The classifier was used to predict TG for the testing cohort factor scores.

We carried out and previously reported deep immunophenotyping from 539 IMPACC participants who were enrolled primarily before September 2020 (6). These data included profiling of targeted proteomics of serum (SPT), global plasma metabolomics (PMG), global and targeted proteomic profiles of plasma (PPG and PPT, respectively), as well as transcriptomics from PBMCs and nasal swabs (denoted as PGX and NGX, respectively). All assays were collectively measured longitudinally across six scheduled visits from 0 days (referred to as baseline) to 28 days post hospital admission (Fig. 1A). In this work, we used these published profiles as training data to construct covarying immune programs and models that predict disease severity and mortality. Molecular immune programs were examined using hold-out assays including immunological outcomes via serum antibody titers, virological outcomes via nasal viral loads, and whole blood cell frequencies measured by mass cytometry (CyTOF). To test the prediction models on independent data, we have now carried out new deep immunophenotyping on an additional 613 IMPACC participants who were enrolled after September 2020. In total, these IMPACC data include 20,544 distinct assay measurements comprising 3,077 multi-omics profiles (referred to hereafter as samples).

### Multi-omics factors predict clinical trajectory groups

We focused on two clinically relevant objectives for stratifying disease severity using baseline multi-omics profiles (26–28) (Fig. 2A): predicting disease severity (identifying factors separating TG1 versus TG2/TG3 versus TG4/TG5; referred to as the “severity task”) and predicting fatal illness among critically ill participants (identifying factors separating TG4 versus TG5; referred to as the “mortality task”). To address the high-dimensionality of the data, we first constructed low-dimensional multi-omics factors using Multiple Co-Inertia Analysis (MCIA) (29) on the 539 IMPACC participants in our training cohort. The dimension reduction combined 27,320 analytes (92 for SPT, 210 for PPT, 1,430 for PPG, 722 for PMG, 12,408 for PGX, 12,458 for NGX) into multi-omics factors that contained covarying patterns across assays.

**Fig. 2.**
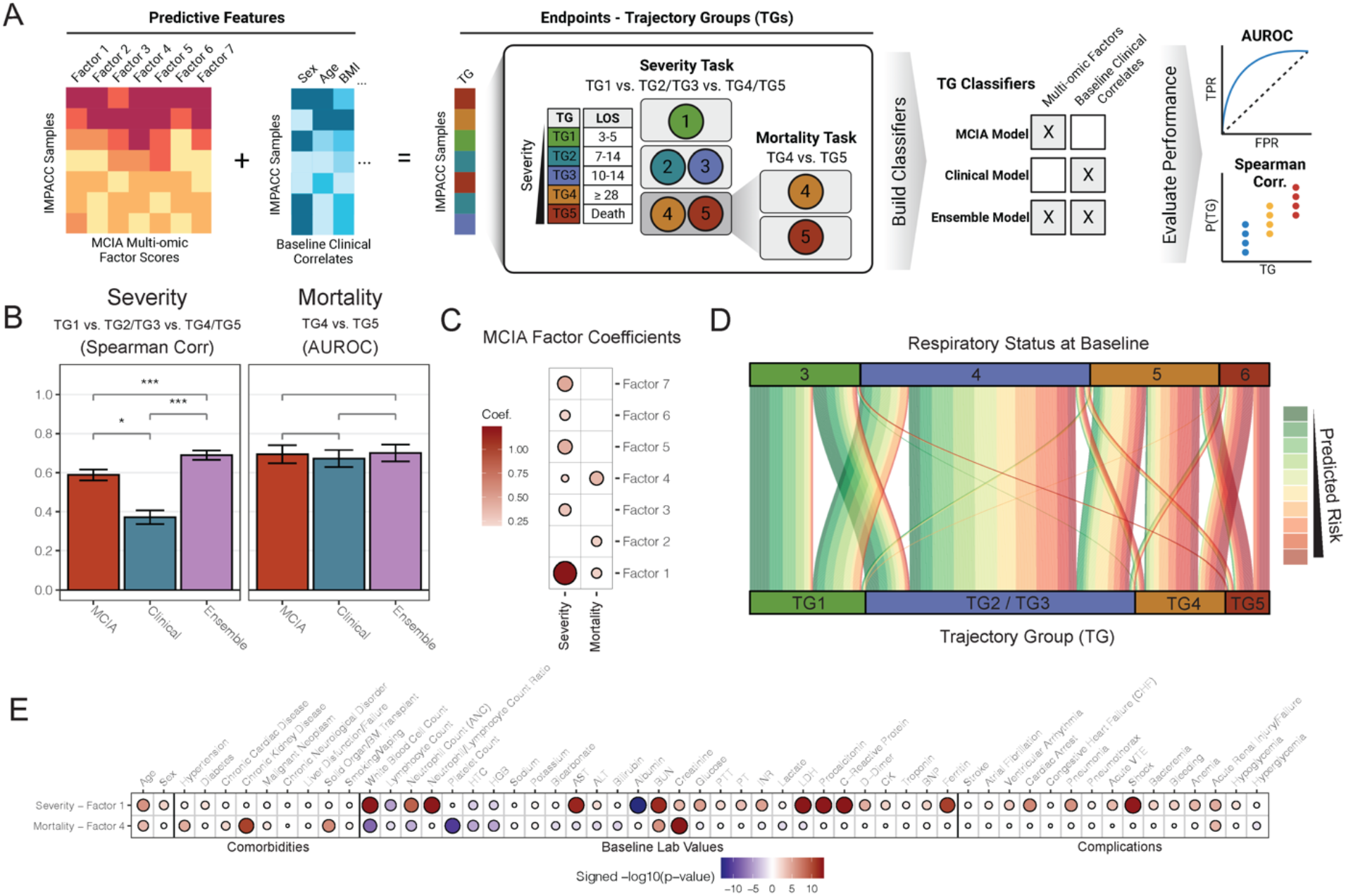
Multi-omics factor prediction results. **A)** Prediction schema (TG – trajectory group, LOS – length of stay). **B)** Spearman correlations and AUROC values for the severity task (TG1, TG2/TG3, TG4/TG5 vs. P(TG4/5)) and mortality task (TG4, TG5 vs. P(TG5|TG4/TG5)) using different baseline models (higher=better performance), evaluated on the test cohort. Significance test was performed via standard normal approximation of the bootstrapped differences (* p<=.05, ** p<=.001, *** p<=.00001). **C)** Dot plot of classification model coefficients for the MCIA model, with size and color representing magnitude and direction respectively. **D)** Alluvial plot showing the distribution of testing cohort individuals in each trajectory group linked to their initial baseline respiratory status. Line color represents the predicted risk from the MCIA model. **E)** Dot plot of Spearman correlation test significance between baseline factor scores and various baseline clinical measurements, complications during hospital stay, and comorbidities, with size and color representing significance and direction respectively. Values are only shown for adj.p<=.05.

We developed prediction models using multi-omics factors from baseline samples (MCIA model) for each task to identify multi-omics factors at hospital admission that correlated with clinical trajectory groups. We also trained models using 28 clinical characteristics (clinical model) and combined multi-omics factors and clinical characteristics (ensemble model), to assess and compare the performance of the different models (see Methods). To validate these trained models, we applied the fitted MCIA factor construction and prediction model to the test cohort of 613 independent IMPACC participants first reported here and evaluated the resulting classification accuracy (Fig. 1C-E).

Both the MCIA model and the ensemble model significantly outperformed the clinical model on the severity task (p.val<1E-6), as measured by Spearman correlation (MCIA: ρ=0.59, ensemble: ρ=0.69, clinical: ρ=0.37) (Fig. 2B). Inspection of the MCIA model coefficients for the severity task revealed a remarkable contribution from multi-omics factor 1 (Factor 1) (Fig. 2C and fig. S1A). The MCIA and ensemble models showed comparable performance on the mortality task to the clinical model, as measured by AUROC (MCIA: AUROC=0.69, ensemble: AUROC=0.70, clinical: AUROC=0.67) (Fig. 2B). While MCIA and the clinical models displayed similar prediction performances for the mortality task, the MCIA model can provide insights into the immune program that distinguishes TG4 and TG5. Key multi-omics factors for the mortality task included multi-omics factor 4 (Factor 4) as the most salient, followed closely by factors 1 and 2 (Fig. 2C and fig. S1A). Predictions from the severity and mortality tasks can be aggregated for TG-specific probabilities per sample (Fig. 2A). The MCIA models also achieved higher prediction accuracy than the clinical models for separating each TG group from the others using the aggregated predictions (fig. S1C).

We further investigated if multi-omics factors can predict COVID-19 disease progression given baseline respiratory status upon hospitalization (3=no supplemental oxygen, 6=invasive mechanical ventilation and/or extracorporeal membrane oxygenation) (5). We assigned a predicted risk for each participant as the geometric mean of being critically ill and experiencing fatal illness from MCIA models, and tested if the predicted risk can predict clinical trajectory group after adjusting for baseline respiratory status (see Supplementary Methods). Remarkably, participants in the test cohort who had the same baseline respiratory status skewed towards more severe TGs with higher predicted risks (Fig. 2D), both among those with moderately impaired baseline respiratory statuses (baseline score in [3,4], p.val=6.9E-08) and those with severely impaired baseline respiratory statuses (baseline score in [5,6], p.val=6.6E-11).

### Multi-omics factors exhibit diverse associations with clinical profiles

Factor 1 and Factor 4 were identified as the strongest contributors to the severity task and the mortality task, respectively (Fig. 2C). Hence, we refer to Factor 1 as the “severity factor” and Factor 4 as the “mortality factor” hereafter. Each of these factors captured correlated metabolite, protein, and gene profiles representing a coordinated immune program. We identified diverse significant associations -- with multiple testing corrected p-values (adj.p) <0.05 -- between these two factors and comorbidities, clinical laboratory testing, and complications during the hospital stay (Fig. 2E). The severity factor showed strong positive associations with baseline total white blood cell (adj.p=6.6E-14) and neutrophil counts (adj.p=1.3E-9) while being negatively associated with lymphocyte count (adj.p=2.0E-5). Significant associations were also seen with baseline values of several acute-phase reactants (adj.p<0.05), including C-Reactive Protein (CRP), LDH, ferritin, procalcitonin, and albumin, and with D-dimer, PT (prothrombin time), PTT (partial thromboplastin time), and INR (International Normalized Ratio), which are used to measure blood clotting functions. Moreover, the severity factor was predictive of high acuity complications such as shock (adj.p=1.3E-13) and cardiac arrest (adj.p=2.4E-7). In contrast, the mortality factor demonstrated a strong negative association with baseline platelet (adj.p=2.4E-11) and total white blood cell (adj.p=2.0E-8), neutrophil (adj.p=9.6E-5), and lymphocyte counts (adj.p=1.7E-2), and strong positive associations with baseline serum creatinine and pre-existing chronic kidney disease, hypertension, and solid organ transplantation (adj.p<0.05). With respect to its association with complications during the hospital course, the mortality factor was most positively associated with acute renal failure (adj.p=8.1E-5).

### The severity factor unravels broad immune dysregulation as hallmarks of severity

While the severity factor and mortality factor were identified by the two prediction tasks using baseline omics measurements, the multi-omics factors were constructed using samples from all visits to capture multi-omics covariations at the baseline visit and over time. At the baseline visit, the severity factor displayed a significant increase with severity and was also elevated in the mortality group compared to other critically ill participants (Fig. 3A, severity adj.p=1.4E-30, mortality adj.p=0.049, table s4), with the difference between groups increasing over time (Fig. 3B). The more moderate groups (TG1-3) showed a sharp reduction in the severity factor levels over time compared to the more severe groups (TG4-5) (adj.p=5.2E-28). Interestingly, while the critical group that survived (TG4) also exhibited a gradual decrease in the severity factor over time, the critical group that died from the disease (TG5) displayed a significant increase in the severity factor level (Fig. 3B, mortality slope adj.p=7.1E-15). These observations suggest that the severity factor captured features that are not only linked with overall COVID-19 severity but are also associated with death among critically ill participants after hospitalization.

**Fig. 3.**
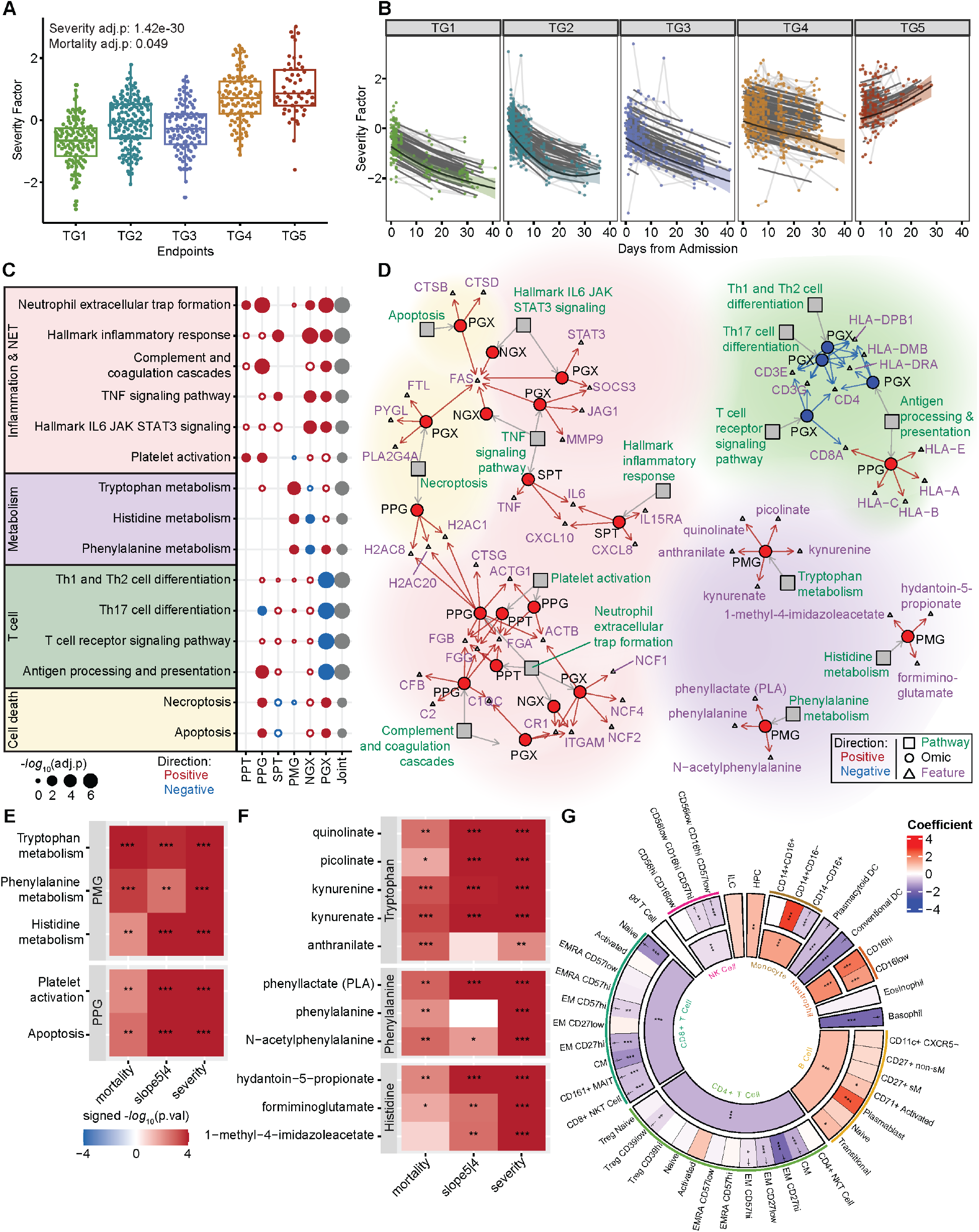
The severity factor increased in severe COVID-19. **(A)** The severity factor scores across clinical trajectory groups at baseline (severity adj. p=1.4E-30, mortality adj.p=0.049). **(B)** Longitudinal trajectory of the severity factor for different clinical trajectory groups (mortality slope adj.p=7.1E-15). Shaded region denotes 95% confidence interval of the fitted trajectory (thick black line), thin black lines show individual participant-fitted models, and light gray lines connect participant’s timepoints. **(C)** Pathway enrichment of the severity factor. **(D)** Network of enriched pathways and selected high-contribution features. The full list of associated features is in table s5. **(E)** Heatmaps of differential expression tests for pathways in (C) that showed early separation between TG4 and TG5. **(F)** Heatmap of differential expression tests of leading-edge metabolites from metabolism pathways in (E). **(G)** Regression coefficients between the severity factor and normalized cell frequencies from whole blood (CyTOF) of both parent and child populations. Daggers indicate a significant association between the reduction of a child cell population frequency which is significantly associated with the severity factor and severity factor apoptosis signaling in PGX. (mortality/severity=baseline mortality/severity task, slope5|4 = TG5 vs.TG4 longitudinally; * p<=.05, ** p<=.01, *** p<=.001; *PPT/PPG=Plasma Proteomics Targeted/Global, SPT =Serum Proteomics Targeted, PMG = Plasma Metabolomics Global, NGX/PGX=Nasal/PBMC gene expression, joint = aggregated p-value across omics*)

We defined high-contribution features for a factor as those highly correlated with this factor (see Methods) and used the high-contribution features to characterize the associated immune program. The severity factor showed strong covarying patterns across the different assays with 58 secreted cytokines/ chemokines (out of 92), 124 targeted plasma proteins (out of 210), 685 global plasma proteins (out of 1430), 366 plasma metabolites (out of 722), 3637 PBMC genes (out of 12408), 768 nasal genes (out of 12458) identified as high-contribution features (table s3). To characterize the biological dynamics underlying the severity factor, a minimum hypergeometric test (mHG) was used to assess the enrichment of known biological pathways among the high-contribution features of the factor, with enrichment being either positive or negative, corresponding to pathways directly and inversely associated with the factor. The pathway databases are comprised of many broad and redundant biological pathways with highly overlapping gene sets (30). To handle this redundancy, biological pathways significantly enriched (adj.p<0.1) in the severity factor were clustered based on the shared leading-edge features and separated into four major functional categories: inflammation, T-cell activity, cell death, and dysregulated metabolism of essential amino acids (fig. S2A, table s5). The joint multi-omics enrichment was calculated by aggregating mHG p-values from different omics within each functional category for prioritization (fig S2B; see Supplementary Methods). The representative pathways from each of the four functional categories that reflected specialized biological functions with the most significant aggregated p-values were selected for further evaluation (Fig 3C). The highlighted functions have been associated with COVID-19 disease severity previously in other study cohorts, but understanding of their coordination remains a challenge.

### Dysregulation of essential amino metabolism as an early hallmark of mortality among critically ill patients

The severity factor was characterized by increased plasma metabolites from tryptophan catabolism (kynurenine and its derivatives, anthranilate and kynurenate, adj.p=1.10e-4), phenylalanine metabolism (phenylalanine and its derivatives, phenyllactate and N-acetylphenylalanine, adj.p=0.056) and histidine metabolism (precursors of glutamate hydantoin-5-propionate, formiminoglutamate, and 1-methyl-4-imidazoleacetate, adj.p=0.0247) (Fig. 3, C and D). Interestingly, the observed metabolic dysregulations were not only associated with overall disease severity at baseline (Fig. 3E severity p.val < 0.05, table s6), but also demonstrated the most power separating TG4 and TG5 at baseline among the pathways in Fig. 3C (Fig. 3E mortality p.val<0.05, fig. S3A, and table s6). This early elevation was particularly interesting because TG4 and TG5 have similar disease severities at the time of hospital admission (Fig. 1B). Furthermore, the metabolic dysregulations were persistently elevated in TG5 throughout hospitalization (Fig. 3E slope5|4 p.val<0.05, and table s6) and were associated with the diverging severity factor kinetics between TG4 and TG5 (fig. S2C). These suggest that dysregulations of essential amino acid metabolisms, including the overaccumulation of downstream products of essential amino acids such as kynurenine and phenylalanine substrates (Fig. 3F), may play a key role in COVID-19 severity and contribute to the sustainment of the broad immune dysregulation captured by the severity factor in fatal illnesses at later stages.

To further evaluate the potential contribution of metabolic dysregulation in plasma to systemic immune dysregulation, we conducted an inter-omics association analysis to directly associate the plasma metabolites with dysregulated cytokines and biological pathways (enriched pathway activities) from proteomics and transcriptomics assays across blood and nasal compartments (see Methods). Indeed, tryptophan, phenylalanine, and histidine pathways showed strong associations with high-contribution soluble proteins, after controlling for demographic covariates, TG groups, and visits (adj.p<0.01), including CD274, CD40, CX3CL1 (Fractalkine), and IL15RA, which exhibited the strongest average associations (fig. S4A). IDO1-driven tryptophan breakdown correlates with the release of HGF (31), and heightened IL-10 (32), IL-6 (33), and TNF levels (34), all of which were also significantly associated with them (fig. S4A). Additionally, the metabolic pathways exhibited strong negative associations with T-cell and antigen presentation pathways in PBMC transcriptomics (fig. S4A). Tryptophan deprivation has been shown to sensitize T cells to apoptosis and inhibit T-cell proliferation (35–37) while the tryptophan catabolite kynurenine induces regulatory T-cell (Treg) development and suppresses cytotoxic T-cell responses (38, 39). Similarly, phenylalanine metabolism has been implicated in regulating the suppression of T-cell immune responses (40).

### T-cell Lymphopenia is associated with increasing COVID-19 severity

The severity factor was also characterized by decreased signatures of T-cell activities in PBMC transcriptomics (e.g., the T-cell receptor signaling pathway (adj.p=7.6E-7) and the antigen processing and presentation pathway(adj.p=9.2E-7), which included genes coding for the T cell receptor complex, such as *CD4*, *CD3E*, *CD3G*, *CD8A,* genes involved in TCR signal transduction such as *NFATC1-3* and *LCK,* and genes coding for the MHC class II molecules, *HLA-DPB1*, *HLA–DRA* and *HLA-DMB)* (Fig. 3, C and D). Reduced T-cell-associated gene expression likely reflected persistent T-cell lymphopenia observed in TG5 participants (fig. S2C). To validate this finding, we correlated whole blood CyTOF cell frequencies against the severity factor and confirmed that the factor was associated with lower circulating T cells (Fig. 3G). These results were also corroborated by the clinical association of lower absolute lymphocyte counts with the factor (Fig. 2G) and were consistent with a targeted analysis on transcriptomics signatures of T cells from the literature (41–43) (fig. S3B). Additionally, PBMC transcriptomic signatures of Th1, Th2, and Th17 cell differentiation pathways exhibited a downward trend over time in TG5 after adjusting for T-cell frequencies (fig. S3C), potentially reflecting dysregulated T-cell functions besides lymphopenia.

Interestingly, the severity factor was also positively enriched in cell death pathways such as Necroptosis (including the genes *PLA2G4A*, *PYGL* and *FTL* coding for inducers of ROS in PBMC transcriptomic; adj.p=0.052), and apoptosis (including the cathepsins genes *CTSD*, *CTSB*, and the gene coding for the cell death receptor *FAS* in PBMC transcriptomics; adj.p.=0.087). Apoptotic pathway activity was positively associated with increased levels of several high-contribution soluble proteins involved in regulating cell death (fig. S3D), such as CASP8 (44, 45), CD274 (PDL1) (46) and TNF (47), and was negatively associated with the CD4+/CD8+ T cell frequencies in whole blood measured by CyTOF (Fig. 3G, daggers). Our results suggest that increased induction of apoptosis, which could be modulated by inflammatory cytokines (Fig. 3D), might contribute to the loss of circulating T-cells observed in the most severe patients (5).

These findings are also consistent with the negative associations between T cell pathways and elevated tryptophan metabolism discussed earlier, which could reflect T cell apoptosis and dysregulated T cell signaling. Interestingly, increased protein levels of CD274 (PD-L1), a ligand for the inhibitory receptor PD-1 on T cells (46), was the top associated feature with the observed metabolic pathways enriched in the severity factor (fig. S4A). The inter-omics association among essential metabolite dysregulation, CD274, and T cell pathway activity revealed an orchestrated suppression of T cell responses across the three assays (metabolomics, serum proteomics and transcriptomics).

### An inflammation and NET formation network associated with worse outcomes

The severity factor was also positively enriched in diverse inflammatory pathways across proteomics and transcriptomics of both the nasal and blood compartments. Within these, inflammatory markers previously associated with COVID-19 severity were among the high-contribution features of the severity factor (Fig. 3D and table s3), including IL6, CXCL10, CXCL8 (IL8) proteins in serum. Proinflammatory soluble proteins are known to modulate metabolism (48, 49), transcription (50, 51) and other cellular activities (52–54). Similarly, our analyses revealed strong associations of proinflammatory soluble proteins with transcriptomic, proteomic and metabolomic pathways and cellular compositions in blood (fig. S4, A and B).

Markers of NET formation were strongly enriched in the severity factor across multiple omics (including the protease CTSG/Cathepsin G, the histone H2AC20, H2AC1, H2AC8 proteins in plasma, and genes encoding Neutrophil NADPH Oxidase Factors *NCF1*, *NCF2* and *NCF4* in PBMC transcriptomics; adj.p=7.6E-7). In support of an increase in NET formation with increasing disease severity, the severity factor also showed positive enrichment of upstream inducers of NET formation, such as IL-6 signaling (Cytokine IL6 and the genes *SOCS3* and *STAT3* in the PBMC transcriptomics; adj.p=1.4E-4), platelet activation (fibrinogens FGA, FGB, and FGG in plasma proteins (55); adj.p=0.02), complement and coagulation (including the plasma proteins C2, CFb and C1QC and genes coding for the *CR1* (complement receptor 1) and *ITGAM* in PBMC transcriptomics; adj.p=2.5E-6). Notably, the platelet activation pathway showed significant elevation in TG5 compared to TG4 at baseline (Fig. 3E and fig. S3A), supporting the possibility that platelet activation might trigger distinct kinetics of NETosis in TG4 versus TG5. Intracellular pathways triggered during NET formation were also enriched, such as actin degradation (actins ACTB and ATCG1 in the plasma proteins (56)) and TNF/NFkB signaling (Cytokine TNF, NFkB target genes *MMP9*, *FAS*, *JAG1*; adj.p=1.6E-4), as were receptors expressed by and cytokines produced by neutrophils, including receptor *IL15RA* (57) and cytokines CXCL8 (IL8) and IL17A (58, 59) (Fig. 3, C and D). Notably, CXCL8 is a neutrophil chemoattractant, indicating a potential increase in neutrophil production or greater recruitment in more severe patients. These enriched inflammatory pathways were elevated over time uniquely in the mortality group, which suggests prolonged and unresolved inflammation associated with neutrophils preceding death (fig. S2C).

The enrichment of NET formation signatures also included ERK and p38 signaling pathways in transcriptomics, which form core inflammatory signaling pathways triggered by many high-contribution cytokines grouped into “Cytokines produced by Macrophages” and “Cytokines produced by Neutrophils” (6) (Fig. 4A). Notably, among these cytokines, IL10, IL6, CXCL10, and CXCL7 were the most strongly associated with the severity factor and are known to elicit ERK and p38 signaling (Fig. 4B). Along with enrichment of ERK and p38 signaling, there was also significant enrichment of AP-1-regulated genes in the PBMC transcriptomics, and the transcriptional factor AP-1 is a downstream target of ERK and cytokine signaling (fig. S3E). AP-1 has been highlighted in previous studies as one of the top COVID-19-associated severity features along with p38 and MAPK signaling (7). Receptors for the top four high-contribution cytokines in the severity factor were detected in PBMC transcriptomics (Fig. 4B), with IL6 receptor components positively enriched in the factor. However, other receptors such as CXCR3, the receptor for CXCL10, were weakly positively or negatively associated with the severity factor, which may reflect a reduction of certain cell types in the circulating PBMCs. Notably, reduced CXCR3 expression is consistent with lymphopenia suggested by reduced clinical absolute lymphocyte counts (Fig. 2G). Such mixed correlations of receptors were also observed for other high-contribution cytokines to the severity factor (fig. S4C).

**Fig. 4.**
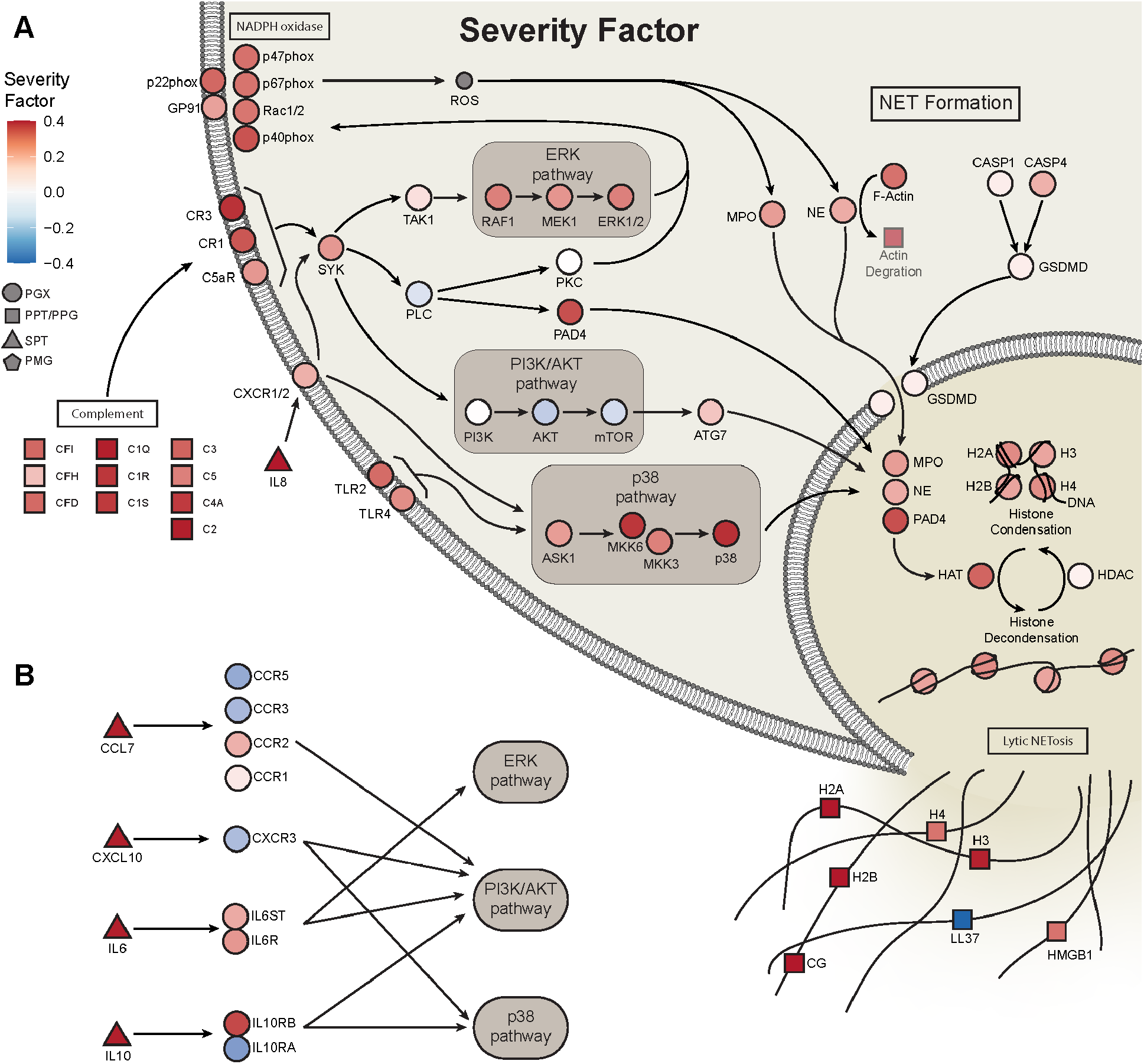
Integrative multi-omics network identifies upstream regulators and mediators of NET formation. **(A)** Broad elevation of transcriptomics and proteomics features in NET formation and Complement in the severity factor. Pathway connections are from the KEGG NET formation pathway. **(B)** Top cytokines in the severity factor, when bound to their receptors, trigger downstream signaling pathways, including ERK and p38 signaling pathways, which play an important role in NET formation. (*PPT/PPG=Plasma Proteomics Targeted/Global, SPT =Serum Proteomics Targeted, PMG = Plasma Metabolomics Global, NGX/PGX=Nasal/PBMC gene expression*)

To complement our pathway analysis, we investigated the correlation of the severity factor with immune cell frequencies in whole blood measured by CyTOF. We also evaluated the overlap of severity factor high-contribution genes with transcriptomic markers of immune cells from blood and nasal tissue (41–43). Consistent with the enrichment of NET formation, the severity factor positively correlated with neutrophil frequencies in whole blood and was enriched for transcriptomic markers of neutrophils in the nasal transcriptomics (Fig. 3G and fig. S3A). The severity factor was also positively correlated with monocytes and was significantly enriched in transcriptomic markers of monocytes (fig. S3B), suggesting that monocytes may play a critical role in the inflammatory response identified and possibly promote NETosis.

### The mortality factor reveals B cell and plasma immunoglobulin reduction as early hallmarks of mortality among critical illness

The mortality factor was significantly higher at baseline in those who died within the first 28 days after hospitalization (TG5) compared to critically ill participants who survived (TG4) (Fig. 5A, severity adj.p=0.14, mortality adj.p=0.049, table s4). Over time, the relative levels of the mortality factor dropped and were sustained at low levels in all groups (Fig. 5B). Hence, this factor represents a mortality-related immune state at hospitalization and is of particular interest as it stratifies mortality from other critically ill patients.

**Fig. 5.**
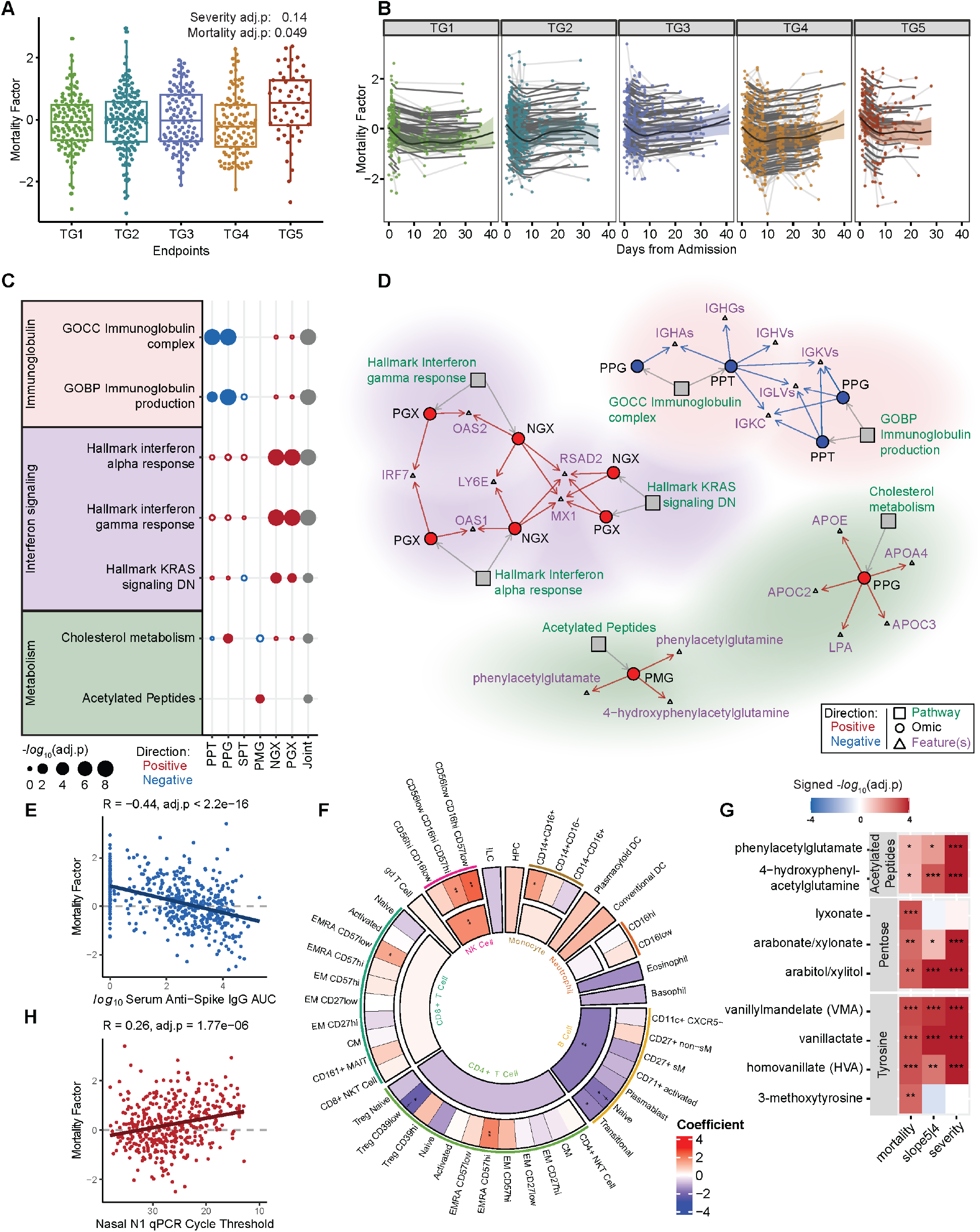
Multi-omics mortality factor enriched for antibodies, interferon signaling and cellular metabolic changes. **(A)** The mortality factor scores across clinical trajectory groups at baseline (severity adj. p=0.14, mortality adj.p=0.049). **(B)** Longitudinal trajectory of the mortality factor for different clinical trajectory groups. Shaded region denotes 95% confidence interval of the fitted trajectory (thick black line), thin black lines show individual participant-fitted models, and light gray lines connect participant’s timepoints. **(C)** Functional pathway enrichment of the mortality factor revealed downregulation of immunoglobulins, upregulation of IFN response, cholesterol metabolism, and acetylated peptides. **(D)** Network of enriched pathways in (C) and top selected high-contribution features. The full list of associated features is in table s7. **(E)** Spearman correlation between the mortality factor and Serum anti-Spike IgG antibody using baseline samples. **(F)** Regression Coefficient of the mortality factor with normalized cell frequencies from whole blood (CyTOF) of both parent and child populations. Daggers indicate a significant association between the reduction of a child cell population frequency which is significantly associated with the mortality factor and severity factor apoptosis signaling in PGX. **(G)** Differential expression tests of leading-edge metabolites in highlighted metabolomic pathways. **(H)** Spearman correlation between the mortality factor and Nasal SARS-CoV-2 qPCR cycle threshold using baseline samples. (mortality/severity=baseline mortality/severity task, slope5|4 = TG5 vs.TG4 longitudinally; * p<=.05, ** p<=.01, *** p<=.001; *PPT/PPG=Plasma Proteomics Targeted/Global, SPT =Serum Proteomics Targeted, PMG = Plasma Metabolomics Global, NGX/PGX=Nasal/PBMC gene expression, joint = aggregated p-value across omics*)

The features with the highest contribution to the mortality factor were primarily plasma proteins and metabolites (table s3), including 89 targeted plasma proteins, 289 global plasma proteins, and 172 plasma metabolites. Only 14 secreted cytokines/chemokines (out of 92), 42 PBMC genes (out of 12408), and 31 nasal genes (out of 12458) are highly contributing features to this factor (table s3). At baseline, there were seven enriched pathways (adj.p<0.1, table s7) that separated TG5 and TG4 from at least one assay (p.val<0.05, Fig. 5C, fig. S5A and table s8).

The most prominent characteristic of the mortality factor was a reduction in immunoglobulins (Ig) in the proteomic assays, including heavy and light chain variable regions and constant regions from multiple isotypes (e.g., IGHGs, IGHAs, IGHVs, IGKVs, IGLVs) (Fig. 5D, adj.p=2e-08 for immunoglobulin complex in PPT; 1.48e-9 for immunoglobulin complex in PPG; 0.00637 for immunoglobulin production in PPT; 3.8e-9 for immunoglobulin production in PPG). The mortality factor was also negatively associated with serum anti-Spike IgG (Fig. 5E, adj.p < 2.2E-16). Furthermore, this factor was negatively correlated with the frequency of total circulating B cells, particularly naïve and transitional B cell subsets measured by CyTOF (Fig. 5F, adj.p=4.59E-19), and total circulating B cells were also positively correlated with plasma Ig (fig. S5B, adj.p = 2.34E0-8). The 42 high-contribution PBMC genes were also negatively enriched for transcriptomic markers of B cells (fig. S5C, adj.p=3.00E-10). These findings suggest that the mortality factor captures a lower level of B cell activity, plasma Igs and anti-Spike IgG at the baseline in patients who died within the first 28 days following hospitalization (fig. S5, D and E). The decline in B cells could partially reflect increased apoptosis, especially of naïve B cells, as suggested by the positive association with the apoptosis pathway constructed from the severity factor (Fig. 5F, adj.p <= 0.05).

### Dysregulated interferon responsiveness and cellular metabolic changes indicate mortality

Alongside the reduced Ig and B cell activity, the mortality factor exhibited a strong positive enrichment of IFN-stimulated genes (ISGs) in PBMC transcriptomics (*OAS1*, *OAS2* encoding viral RNA sensors, and *IRF7*, which encodes a key transcription factor mediating regulation of ISGs, see Fig. 5D) and in nasal transcriptomics (*MX1*, *RSAD2*, *LY6E* coding for antiviral immune genes). Further examination revealed that the leading-edge ISGs are enriched in antiviral rather than proviral functions (60, 61). IRF-regulated and STAT-transcribed genes were also positively enriched in the mortality factor across both nasal and PBMC transcriptomics (fig. S6A), confirming the propagation of IFN signal through transcriptional factor activity. Along with elevated JAK/STAT IFN signaling, known inhibitors of IFN signaling (IFN inhibitors) (62), including *USP18*, *SOCS1*, and *PIAS4,* were positively enriched in the mortality factor (p.val=0.030), suggesting the possibility of dysregulated IFN responsiveness in critically ill patients.

The mortality factor was also enriched in acetylated peptides (4-hydroxyphenylacetylglutamine, phenylacetylglutamate, and phenylacetylglutamine) in the plasma metabolites (adj.p=0.08) and cholesterol synthesis related plasma proteins (adj.p=1.75e-4), including APOC2, APOC3, APOA4, APOE, and LPA (Fig. 5, C and D). To comprehensively explore soluble markers contributing to the early separation between TG4 and TG5 in the mortality factor, we performed enrichment analysis on high-contribution metabolites and cytokines that also separated TG4 and TG5 at baseline (p.val < 0.05, table s8). We identified positive enrichment of pentose metabolism (including lyxonate, arabitol/xylitol, arabonate/xylonate, adj.p =0.07) and tyrosine metabolism (including VMA, HVA, vanillactate, 3-methoxytyrosine, adj.p=0.07), in addition to acetylated peptides, whose leading-edge metabolites showed separation between TG4 and TG5 at hospital admission (Fig. 5G and table s8). An inter-omics association analysis further revealed significant associations between the three highlighted metabolomics pathways and proteomics functions, including Ig complex reduction and cholesterol metabolism elevation, as well as soluble proteins CST5, CX3CL1, CCL25, CSF1 and KITLG (adj.p<0.01, fig. S6B).

We note that the mortality factor was also negatively correlated with platelet count on hospital admission (Fig. 2F, adj.p=2.36E-11), which was consistent with the observed positive enrichment in complement and coagulation pathways in plasma proteomics (including FGA, FGB, C3, C4A, C4B, and C9, adj.p=0.007 (PPT), adj.p=0.03 (PPG), table s7). Additionally, the mortality factor was positively enriched in participants with pre-existing chronic kidney disease and renal complications during the hospital stay. Moreover, clinical laboratory testing demonstrated a positive correlation between the mortality factor and baseline clinical creatinine, a biomarker of kidney function, in addition to creatine being a high-contribution feature to the mortality factor in plasma metabolomics (table s3). Creatinine was noted to have a strong positive correlation with acetylated peptides, cholesterol metabolism, tyrosine metabolism, and pentose metabolism (ρ=0.46, 0.24, 0.53, 0.53; adj.p=2.10E-65, 6.69E-18, 2.98E-92, 9.82E-91 respectively, fig. S6C).

### The mortality factor reveals a dysregulated viral response immune cascade

Reduction of total plasma immunoglobulins and elevation of IFN-stimulated genes were the most prominent features of the mortality factor, together with strong negative associations with serum SARS-CoV-2 antibody titers and a robust positive association with nasal viral loads (Fig. 5H). This suggests a dysregulated host immune cascade may contribute to failure of viral clearance in TG5. Notably, the total immunoglobulin level was strongly associated with the serum SARS-CoV-2 antibody titer levels and has an inverse relationship with viral load (Fig 6A and fig. S6C, adj.p<=0.05).

**Fig. 6.**
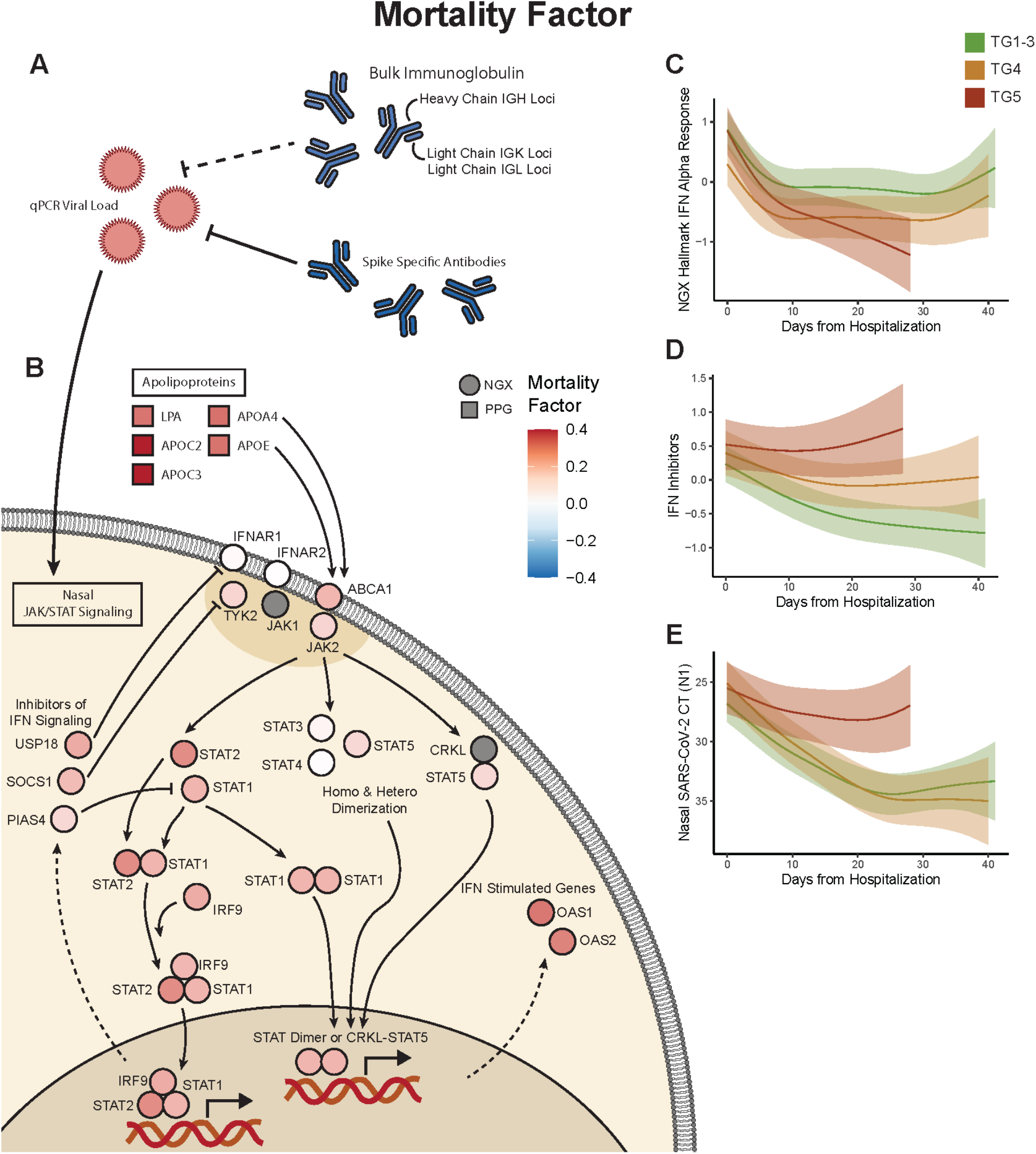
Virus-centered integrative multi-omics network of the mortality factor. **(A)** Positive association of nasal SARS-CoV-2 viral load and inverse associations of total and SARS-CoV-2 specific antibodies were top features of the mortality factor. **(B)** JAK/STAT IFN signaling was positively associated with the mortality factor and viral load, and IFN signaling inhibitors are also positively associated with the mortality factor, potentially contributing to the dysregulation of IFN responsiveness and uncontrolled viral load TG5 despite early ISG elevation. The elevation of Apolipoproteins from the plasma proteomics may also contribute to the heightened STAT activity. Longitudinal trajectories from generalized additive mixed modeling of **(C)** Hallmark IFN alpha response genes in the NGX, **(D)** IFN inhibitors in the NGX, and **(E)** nasal SARS-CoV-2 viral load determined from RT-qPCR Cycle Threshold (CT). (*PPT/PPG=Plasma Proteomics Targeted/Global, SPT =Serum Proteomics Targeted, PMG = Plasma Metabolomics Global, NGX/PGX=Nasal/PBMC gene expression*)

In contrast, both IFN pathway activity and IFN inhibitor levels showed a strong inverse relationship with serum SARS-CoV-2 antibody titers and total immunoglobulins, and a positive correlation with viral loads, in both nasal (Fig. 6B and fig. S6C) and PBMC (fig. S7 and fig. S6C). The observed elevation of nasal ISGs in TG5 at baseline potentially reflected virus associated IFN production. After adjusting for viral load, the nasal ISG levels became more comparable between TG4 and TG5 (fig. S6D, p.val=6E-3 before adjustment, p.val=0.11 after adjustment, see Supplementary Methods). This adjustment accentuated the overall lower ISG expression levels in the more severe versus moderate participant groups (TG4/5 versus TG1-3, fig. S6D), consistent with reduced IFN responsiveness among severe patients observed in previous studies (13). Interestingly, type I IFN gene expression in nasal transcriptomics declined more rapidly in TG5 than TG4, both before and after adjustment (Fig. 6C and S6E, p.val=9E-3 before adjustment and p.val=7E-3 after adjustment). This significantly faster ISG decline in TG5 was accompanied by elevated expression of known inhibitors of IFN signaling (Fig. 6D, adj.p=0.0015), which was uniquely observed in TG5 and matched the trend of viral load in TG5 (Fig. 6E). These observations supported the possibility of dysregulated IFN responsiveness among critically ill participants which was unresolved in those who succumbed to infection, suggesting that higher viral loads could trigger elevated IFN signaling that may counterproductively turn on a negative feedback loop to suppress IFN signaling before the virus is cleared (62, 63), and thereby contributing to mortality (Fig. 6, B to E). Additionally, plasma proteomics demonstrated an elevation of apolipoproteins that can bind receptors capable of activating JAK signaling, possibly contributing to the heightened STAT activity (64) (Fig. 6B). Overall, our results suggest mortality among critically ill patients can significantly associate with dysregulation of the systemic and tissue host transcriptome, plasma metabolome and proteome, and loss of B cells, which was concomitant with the persistent viremia observed in the critically ill participants that died from the disease in the first 28 days.

## DISCUSSION

Our study represents one of the most extensive evaluations of patients hospitalized with COVID-19, encompassing a wide range of omics. As participants were studied before the widespread availability of SARS-CoV-2 vaccines, our study provides a unique perspective on the naïve response to the virus. The use of a previously published 539 participant training cohort and validation with a newly introduced 613 participant test cohort which varied in demographics, clinical laboratory testing (table s2) and enrollment periods during the pandemic, enabled robust identification of multi-omics factors associated with COVID-19 severity and mortality. These factors outperformed clinical features previously studied clinical features (5), single-omic analytes, and several single assay COVID-19 molecular signatures reported in the literature (65–67) in predictive modeling (fig. S1I).

A key severity factor (Factor 1) captured the severity trend among all hospitalized participants (TG1 vs. TG2/TG3 vs. TG4/TG5) and distinguished the critically ill group who survived during the first month (TG4) from the mortality group (TG5), with its level elevated over time uniquely in participants from TG5. This severity factor illuminates a spectrum of immune changes across six omics platforms extending beyond mere cellular alterations, which contains an increase in metabolites related to tryptophan, phenylalanine, and histidine pathways, enhanced signatures of inflammation coagulation, and NET formation, heightened signatures of cell apoptosis, and a decrease in T cell signatures and circulating T-cell numbers, which also showed strong associations with metabolic dysregulations and proinflammatory soluble proteins (Fig. 7A). Elevated serum concentrations of multiple circulating cytokines and chemokines were associated with the severity factor, including elevation of two clusters of soluble proteins characterized by “Cytokines produced by Neutrophils” and “Cytokines produced by Macrophages” as well as a negative association with the cluster “Activators of cytotoxic NKs” (fig. S4). The elevated longitudinal trend in TG5 thus could reflect an unresolved dysregulation and elevation of inflammatory cytokines in fatal illness when the cytotoxic antiviral activities of T cells and NK cells were insufficient to reduce viral burden. The negative association with activators of cytotoxic NKs cluster was consistent with previous analysis results from the targeted serum proteomics assay alone, which identified this cytotoxic NKs cluster as a marker of recovery (6).

**Fig. 7.**
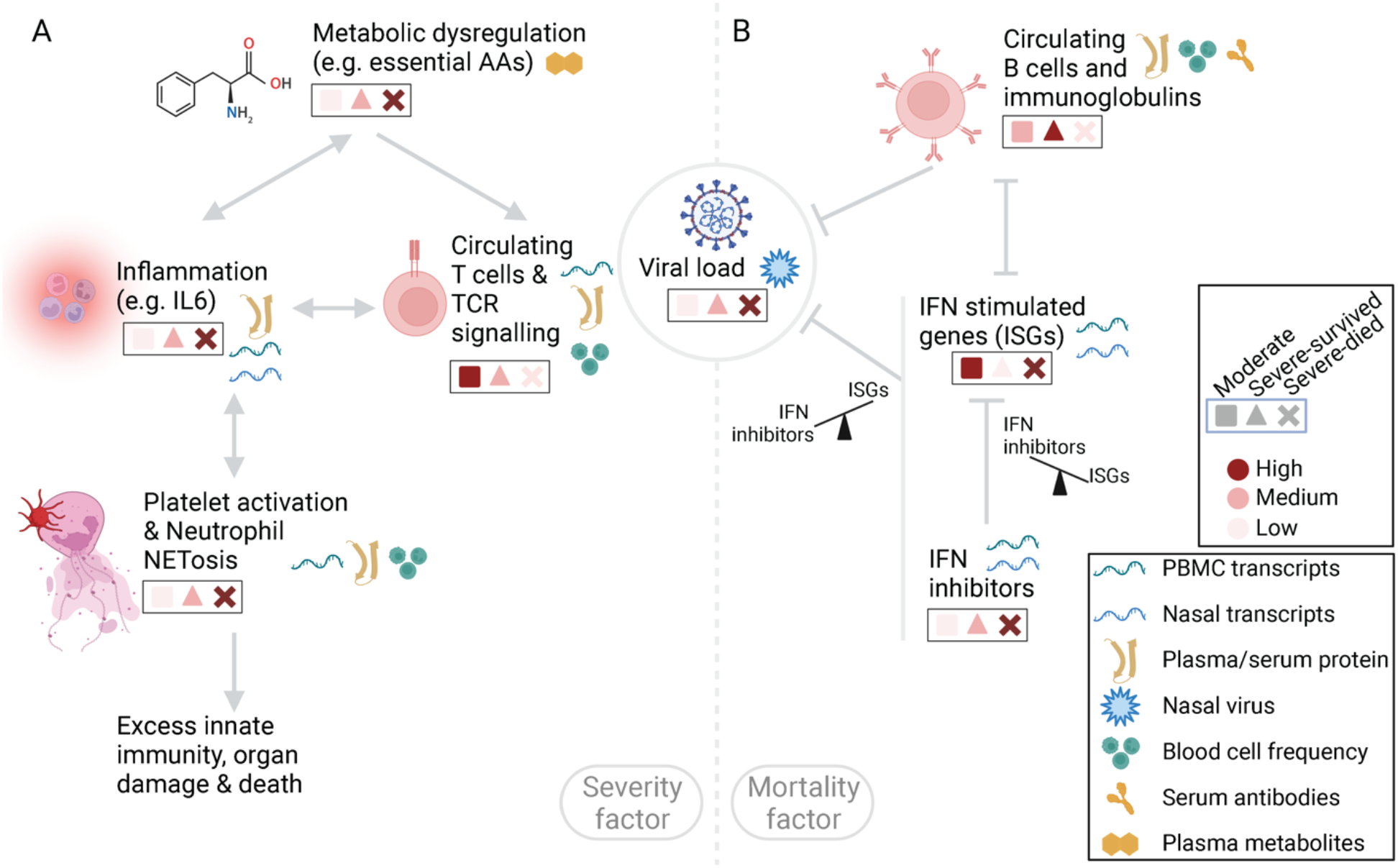
Summary of highlighted host immune programs. **(A**): The severity factor identified an integrated multi-omics cascade associated with disease severity, characterized by dysregulated metabolisms, e.g., essential amino acid (AA) metabolism, and elevated inflammatory soluble proteins and transcripts, signature of coagulation and NETosis, and reduced T cell circulation and signaling in more severe patients. The dysregulated metabolisms potentially served as early modulators of this broadly dysregulated immune state. The links in this panel did not yield conclusive evidence but rather reflected hypotheses formulated based on our findings. **(B**) The mortality factor revealed a viral-centered multi-omics immune state as early hallmarks of mortality among most critical patients (ICU, ventilation, or mortality), including reduced immunoglobulins and B cell circulation, dysregulated interferon responsiveness as suggested by elevated IFN inhibitor levels in both nasal and PBMC transcriptomics, along with persistently elevated viral loads in patients with fatal illness. (*PPT/PPG=Plasma Proteomics Targeted/Global, SPT =Serum Proteomics Targeted, PMG = Plasma Metabolomics Global, NGX/PGX=Nasal/PBMC gene expression*)

A joint investigation of molecular signatures’ baseline levels and kinetics in the severity factor revealed that essential amino acid metabolism dysregulation potentially contributed to its distinct kinetics between TG4 and TG5. These amino acids, and their metabolites, act as important protein building blocks, key energy sources in metabolic pathways (e.g., Kreb’s cycle), and modulators of immunity (68). Our analysis also suggested the immune modulatory role of metabolites associated with the severity factor, shown by the strong associations of essential amino acid dysregulation and high-contribution cytokines produced by neutrophils and macrophages, as well as protein and transcriptomic pathways in the severity factor. For example, significant associations observed between CD274 (PD-L1), T-cell pathway activity, and dysregulated tryptophan metabolism suggested a coordinated suppression of T cell responses across metabolomics, plasma/serum proteomics, and transcriptomics. Furthermore, essential amino acid metabolism dysregulation was significantly associated with TNF, IL-6, IL-10, HGF, and CD40, all linked to IDO1-driven tryptophan breakdowns (31–34). Interestingly, metabolites resulting from IDO1-driven tryptophan breakdowns, such as kynurenine, kynureninate, and 3-hydroxykynurenine, were highly elevated in the severity factor, suggesting functional activity of IDO1. Additionally, IDO1 participates in the phenylalanine catabolic pathway, resulting in the formation of phenylpyruvate. The functional activity of IDO1 was also suggested by the positive correlation of phenylalanine pathway metabolites with HPD, FAH, and SLC16A1 in PBMC transcriptomics, which are associated with the active conversion of phenylpyruvate from phenylalanine (69–71) (fig. S3F). These observations suggest an opportunity to evaluate these metabolites clinically in early COVID-19 to assess disease severity, and intervention via modulation of nutrients may prove beneficial, with several clinical trials currently underway in COVID-19 patients to evaluate amino acid supplementation (72–74).

Another highlighted factor, the mortality factor (Factor 4) was significantly higher at baseline in TG5 compared to TG4, and captured an immune program of dysregulated IFN signaling, elevated nasal viral load, and a reduction in circulating B cells, bulk Ig and SARS-CoV-2 specific serum IgG (Fig. 7B). Both SARS-CoV-2 specific IgGs and total plasma Ig were reduced in the mortality factor and were negatively associated with viral load. These findings suggest that B cell immune responses are important for controlling the virus among critically ill patients during early infection. In addition, the overall reduction in circulating Ig in participants who failed to recover could contribute to general inflammation, as intravenous immunoglobulin (IVIG) therapy suppresses inflammatory pathology (75–77). Thus, the reduction of B-cell activity and circulating Ig in patients who succumb to COVID-19 during the first month after hospitalization might not only allow higher viral replication in critical illness, which could fuel additional inflammation, but could also potentially promote inflammation indirectly, resulting in tissue damage.

Antiviral ISG activity increased with the mortality factor and was positively associated with viral load, consistent with a virus-induced increase in type I interferons (78). Remarkably, while ISG levels were relatively higher in mortality group TG5 compared to TG4 at baseline, the expression of IFN inhibitory genes in the upper respiratory tract was also elevated in TG4 and TG5 at baseline which could suggest suppression of effective IFN signaling in critically ill patients. In addition to the elevated IFN inhibitory genes in TG4 and TG5 at baseline, TG4 and TG5 had a higher proportion of participants who displayed anti-IFN antibodies, which are associated with COVID-19 severity and can inhibit IFN and STAT signaling (17) (fig. S6, F and G). When evaluated longitudinally, IFN-inhibitory genes were uniquely elevated in TG5, which could also contribute to the faster decline in ISGs in TG5 compared to other groups, leading to inadequate IFN signaling in the mortality group TG5. Notably, viral load remained high uniquely in TG5 over time, suggesting failed control of viral clearance. The combined results from analyzing viral loads, Ig levels and IFN signaling/inhibitor genes raise the possibility that while IFN-induced viral clearance may be successful in moderate patients (TG1-TG3), IFN responsiveness might be dysregulated, potentially due to elevated expression of IFN inhibitors driven by sustained high viral loads in the critically ill patients (TG4 and TG5).The observed dynamics of viral load, IFN, and antibodies support the early administration of antiviral and antibody therapies as essential interventions to reduce mortality. The mortality factor stratified participants in TG4 versus TG5 primarily during early stages of hospitalization, indicating the importance of timely interventions, as supported by clinical trial data on paxlovid (79, 80). Several trials showed early monoclonal antibody (mAb) therapies in outpatients to reduce hospitalizations and severe disease when the mAb matched the virus variants in circulation (81–83). However, IVIG and convalescent plasma therapies failed to meet their primary endpoints in several acute COVID-19 clinical trials, with their roles remaining inconclusive (84–88). One potential explanation for the failure may be related to the timing of administration of these therapies and patient inclusion criteria in the trials, both of which could be critical factors for intervention, as suggested by our analyses.

The mortality factor was strongly associated with specific disease comorbidities including chronic kidney disease, immunosuppression, and hypertension, which can potentially explain some high-contribution molecular markers in the mortality factor such as creatinine. Additionally, acute renal disease, a frequent complication in critically ill patients, was linked to the mortality factor. This association may account for the observed increase in tyrosine and pentose metabolites, typically cleared by the kidneys, as well as the mortality factor’s correlation with elevated creatinine and BUN levels (89–91). The tyrosine metabolites (HVA, VMA, and VLA) are also byproducts of catecholamine biosynthesis/degradation and are major terminal urinary metabolites converted from L-Dopa, dopamine, and norepinephrine which may reflect the use of exogenous pressors in ventilated patients (fig. S3F). Furthermore, previous work has suggested that dopamine inhibits SARS-CoV-2 viral replication and stimulates type-I IFNs, with SARS-CoV-2 possibly disrupting dopamine pathways which could also result in increased catecholamine downstream byproducts (92, 93).

Another interesting observation from comparing the severity and mortality factors is that “Complement and coagulation” was a significantly enriched functional pathway in both factors despite their low correlation. Overlap between the factors could reflect different biological processes that influence the complement and the coagulation pathway activities. Although the two factors shared many leading-edge proteins, the severity factor was more strongly associated with classical complement-associated proteins (C1Q, C1R, C1S, C2), whereas the mortality factor had a more significant association with the alternative pathway (C3, FB, FD). The severity factor was enriched in inflammation, platelet activation, and NET formation, which are closely related to the complement and coagulation cascades (94). Using baseline clinical laboratory measures, we observed that the severity factor was positively associated with D-dimer, partial thromboplastin time (PTT) and prothrombin time (PT), suggesting a relationship to coagulopathy in COVID-19 patients (95). In contrast, the mortality factor was negatively associated with platelet count and demonstrated no significant association with D-dimer, PT or PTT. These associations collectively suggest that impaired coagulation is associated with severe disease, while prolonged thrombocytopenia may be predictive of mortality. Our findings contribute mechanistic evidence to the evolving concept of “immunothrombosis”, which refers to the complex interplay between immune cells, complement, coagulation factors, and NETs, that play an important role in COVID-19 associated coagulopathy (12, 96, 97).

Our integrative longitudinal analysis yields novel insights and formulates mechanistic hypotheses concerning temporal coordination, elucidating the heterogeneity in disease progression among hospitalized patients. Our expansive analysis across training and validation cohorts corroborates these hypotheses. While not facilitating further validation through functional assays, this study establishes a substantive foundation for future, in-depth investigations of detailed mechanisms behind disease heterogeneity. Although all participants in the IMPACC cohort are hospitalized due to COVID-19 as part of the study design (23) and may exhibit molecular patterns non-representative of non-hospitalized patients yet providing complement data to the extensive body of existing work comparing healthy controls, mild non-hospitalized patients, and severe COVID-19 cases. Of note, the IMPACC deep multi-omics immunophenotyping data presents a rich resource for further investigations not covered by this study. For example, while the severity factor showed many significant pathways in the enrichment analysis, we focused on the specialized biological functions corroborated by multiple assays except for metabolism functions due to the limitation of existing knowledge bases. Broader or single-dataset-restricted pathways, unexplored in this manuscript, can illuminate additional complex aspects of COVID-19. In addition, the present manuscript did not explore sub-analyses on other questions of interest, including differences in immune responses by age, sex, COVID-19-associated comorbidities, medication, and post-acute-sequelae of COVID-19, which were planned for future follow-up work.

## METHODS

### Study design

IMPACC is a prospective longitudinal study designed to enroll 1,000+ hospitalized patients with COVID-19 (23). The study cohort was enrolled from 20 hospitals affiliated with 15 geographically distributed academic institutions across the U.S. Eligible participants were patients hospitalized with symptoms or signs consistent with COVID-19, and those with SARS-CoV-2 infection confirmed by RT-PCR to remain in the study. Detailed clinical assessments and nasal swabs, blood, and endotracheal aspirates (intubated patients only) were collected within 72h of hospitalization (Visit 1) and on days 4, 7, 14, 21, 28 after hospital admission for 1,152 participants, amounting to 3,077 sampling events. The levels of plasma proteomics (global and targeted; PPG and PPT, respectively), serum proteins (SPT), plasma metabolites (PMG) and nasal and PBMC mRNAs (NGX & PGX, respectively) were measured. The cohort was divided into training (1,493) and test (1,584) sets. The MCIA model was constructed from the preprocessed training multi-omics dataset, which was used on the test cohorts to achieve predictions on the testing cohort. The MCIA factors that robustly captured COVID-19 severity and mortality were identified and were analyzed to comprehensively characterize the cross-tissue, multi-omics immunological signatures that were associated with the severity and mortality. The measurements of nasal viral load, serum SARS-CoV-2 antibody titers, and whole blood CyTOF on the same cohort was used to validate the immunological signatures and to develop further mechanistic insights on immune programs of COVID-19 severity and mortality. More details regarding the study cohort, sample processing and batch correction, and multi-omics construction can be found in Supplementary Methods.

### Statistics

#### Baseline classifiers of TG groups

We constructed a hierarchical classifier using multi-omics factors and clinical features from baseline on the training cohort, where we constructed a regularized ordinal regression model capturing the overall severity increase and a sub-model separating TG5 from TG4 via regularized logistic regression. Model tuning was done through cross-validation on the training cohort. The ensemble model combines the MCIA model and the clinical model through a weighted average, with the weights determined via cross-validation on the training cohort. Detailed information on classifier construction, parameter tuning, and comparisons of different prediction models--including models utilizing individual omics features, concatenated features from all six omics, and proposed signatures from the literature--are presented in the Supplementary Methods.

#### Definition of high-contribution features

The MCIA dimension reduction uses all features in its multi-omics factor construction. To identify the top contributing features associated with each MCIA multi-omics factor, we defined high-contribution features for each assay as the features whose absolute factor weights >=0.2 and Benjamini-Hochberg adjusted p-values for correlation with a factor satisfying p.adj<=0.01.

#### Factor annotation and enrichment analysis

Functional enrichment analysis of multi-omics factors was conducted using the minimum hypergeometric test (mHG) on high-contribution features associated with each factor. We utilized publicly available databases, including KEGG, Hallmark, Subpathway provided by Metabolon, Go Immunoglobulin gene sets, MSigDB C3, Ligand-receptor signaling gene sets from Omnipath, and IFN inhibitors from prior literature, for functional annotations. We will only consider functions with joint enrichment significance that meet the Benjamini-Hochberg corrected FDR/adj.p<0.1 criterion. For all enrichment tests, we used analytes measured in the corresponding assay as the background. Additional details regarding the databases, enrichment tests, and enriched pathway selections for severity and mortality factors can be found in the Supplementary Methods

#### Pathway activity construction

The pathway activities were calculated as a weighted sum of selected features of a pathway. Specifically, for pathways included in the enrichment analysis, we identified the features that contributed to the enrichment of the pathway, also known as the leading-edge features. We multiplied the levels of each leading-edge feature with its absolute factor weight in the factor and calculated a sum of the weighted levels across the features as pathway activity. For an unbiased evaluation of the NGX IFN inhibitory genes and NGX Hallmark IFN Alpha Response in Fig. 6, the pathway activities were instead calculated and modeled as the average of all detected features in the gene sets.

#### Differential expression analysis at baseline and longitudinally with mixed effect modeling

Mixed effect regressions were used for differential expression test of multi-omics factors, pathway activities, and omics analytes across TG groups, after controlling for age, sex, and enrollment site (random effect). They were also used to assess the significance of predicted risks from the baseline MCIA classifier for inferring disease progression after further controlling for baseline respiratory status in Figure 2D. Mixed effect additive regressions were used for identifying kinetic differences in multi-omics factors, pathway activities and omics analytes across TG groups, where, in addition to age, sex, and enrollment sites, participant id was included as a random effect nested with the enrollment site to account for correlated samples from the same participant. We examined if the average (referred to as intercept in the gamm4 documentation) or shape (referred to as the smoothing term in the gamm4 documentation) differ across the clinical trajectory groups. Kinetics directions from linear mixed effect regression were extracted to denote the directionality of the shape (slope/trend) in the additive model. Please refer to the Supplementary Methods for additional details. Multiple test correction was performed using the Benjamini-Hochberg procedure.

#### Inter-omics association analysis

The inter-omics association analysis was conducted to test for strong associations between two different omics. We test for strong associations using the Pearson correlation test after adjusting for age, sex, and enrollment site. In addition, we adjusted visit numbers and clinical trajectory groups to account for the global co-varying patterns and the feature selection effects when evaluating the associations among high-contribution features. We used Bonferroni correction to control the family-wise error rate (FWER) with adj.p<0.01 to identify strong associations between assays. This inter-omics analysis evaluated the direct associations between high-contribution functions/signatures, providing more details than the MCIA construction, which evaluates the associations between analytes and the factor instead. It also complements the multi-omics functional enrichment analysis by linking analytes from the targeted assay SPT, where significant pathways are scarce, to significant pathways from other assays. More details can be found in the Supplementary Methods.

#### Study approval&rdquo

NIAID staff conferred with the Department of Health and Human Services Office for Human Research Protections (OHRP) regarding potential applicability of the public health surveillance exception [45CFR46.102] to the IMPACC study protocol. OHRP concurred that the study satisfied criteria for the public health surveillance exception, and the IMPACC study team sent the study protocol, and participant information sheet for review, and assessment to institutional review boards (IRBs) at participating institutions. Twelve institutions elected to conduct the study as public health surveillance, while 3 sites with prior IRB-approved biobanking protocols elected to integrate and conduct IMPACC under their institutional protocols (University of Texas at Austin, IRB 2020-04-0117; University of California San Francisco, IRB 20-30497; Case Western Reserve University, IRB STUDY20200573) with informed consent requirements. Participants enrolled under the public health surveillance exclusion were provided information sheets describing the study, samples to be collected, and plans for data de-identification, and use. Those that requested not to participate after reviewing the information sheet were not enrolled. In addition, participants did not receive compensation for study participation while inpatient, and subsequently were offered compensation during outpatient follow-ups.

## Data availability

Data files are available at ImmPort under accession number SDY1760 and dbGAP accession number phs002686.v1.p1. All analysis codes have been deposited at https://bitbucket.org/kleinstein/impacc-public-code and are publicly available. DOIs are listed in the key resources table.

## Supplementary Materials

Supplementary Materials and Methods. Supplementary Methods

Fig. S1. Additional computational results for prediction models, severity and mortality task, and MOFA imputation, related to Figure 2

Fig. S2. Enrichment term clustering and selected pathway trajectories for the severity factor, related to Figure 3

Fig. S3. Additional characterization of immune pathway associated with COVID-19 severity, related to Figures 3 and 4

Fig. S4. Inter-omics analysis on top-contribution cytokines and significant pathways for the severity factor, related to Figures 3 and 4

Fig. S5. Additional characterization of immune pathway associated with COVID-19 mortality, related to Figure 5.

Fig. S6. Additional characterization of interferon signaling, anti-IFN autoantibodies, and inter-omics analysis of top-contribution cytokines and significant pathways for the severity factor, related to Figures 5 and 6.

Fig. S7. Virus-centered integrative multi-omics network of the mortality(-associated) factor (Factor 4) in PGX, related to Figure 6

Table s1. Baseline Characteristic by Clinical Trajectory Groups. p-value from chi-square test for categorical variables and Kruskal-Wallis test for continuous variables (age, SOFA score). Trajectory 1= brief length of stay; trajectory 2= intermediate length of stay; trajectory 3= intermediate length of stay with discharge limitations; trajectory 4= prolonged hospitalization; trajectory 5= fatal. ^1^not including asthma, ^2^current or former.

Table s2. Baseline Characteristic by Cohort. p-value from chi-square test for categorical variables and Kruskal-Wallis test for continuous variables (age, SOFA score). Trajectory 1= brief length of stay; trajectory 2= intermediate length of stay; trajectory 3= intermediate length of stay with discharge limitations; trajectory 4= prolonged hospitalization; trajectory 5= fatal. ^1^not including asthma, ^2^current or former.

Table s3. Analyte contributions for MCIA factors

Table s4. MCIA factors baseline and longitudinal testing Table s5. Functional enrichment analysis of Factor 1

Table s6. Functional enrichment analysis of Factor 1, filtered for baseline TG4 and TG5 separation

Table s7. Functional enrichment analysis of Factor 4

Table s8. Functional enrichment analysis of Factor 4, filtered for baseline TG4 and TG5 separation

Table s9. Batch-effect evaluation using PVCA

## Acknowledgments

The Clinical and Data Coordinating Center: Sanya Thomas, Mitchell Cooney, Shun Rao, Sofia Vignolo, Elena Morrocchi. David Geffen School of Medicine at the University of California-Los Angeles: Arash Naeim, Marianne Bernardo, Sarahmay Sanchez, Shannon Intluxay, Clara Magyar, Jenny Brook, Estefania Ramires-Sanchez, Megan Llamas, Claudia Perdomo, Clara E. Magyar, Jennifer A. Fulcher, and the UCLA Center for Pathology Research Services and the Pathology Research Portal. Yale School of Medicine: M. Catherine Muenker, Dimitri Duvilaire, Maxine Kuang, William Ruff, Khadir Raddassi, Denise Shepherd, Haowei Wang, Omkar Chaudhary, Syim Salahuddin, John Fournier, Michael Rainone, Maxine Kuang. Figures 1, 2A, and 7 were created with Biorender.com.

## Author contributions

Conceptualization: JG, CM, RKP, IMPACC Network, JDA, EM, KS, GF, PMB, ADA, LIRE, BjP, BaP, SHK, RPS, SF, LG

Formal analysis: JG, CM, RKP, PS, AK, LX, AH, LG Software: AK, RKP, CM, JG, LG

Methodology: LG

Resources: SB, SKS, FK, LR, HvB, MW, HS, WE, CL, OL, MCA, HM, RRM

Funding Acquisition: IMPACC Network Supervision: GP, LIRR, BJP, SF, SHK, LG

All authors wrote, edited, and reviewed the manuscript.

