## Supplementary Materials for "Integrated longitudinal multi-omics study identifies immune programs associated with COVID-19 severity and mortality in 1152 hospitalized participants"

**Supplementary Methods**


- **RESOURCE AVAILABILITY**
  - Lead contact
  - Materials availability
  - Data and code availability
- **EXPERIMENTAL MODEL AND STUDY PARTICIPANT DETAILS**
  - IMPACC Cohort characteristics
- **METHOD DETAILS**
  - Sample processing and batch randomization
  - Assay preparation and processing
- **QUANTIFICATION AND STATISTICAL ANALYSIS**
  - Cohort definition
  - Data preprocessing
  - Data imputation via MOFA
  - Multi-omic factor construction via MCIA
  - Hierarchical classification model construction and evaluation
    - MCIA model construction and rank selection
    - Clinical model construction
    - Ensemble (MCIA + clinical) model construction
    - Model exploration and comparisons on training cohort via cross-validation
  - MCIA separates TG groups with aggregated predictions
  - MCIA predicted risk and hypothesis testing conditional on baseline
  - Baseline and longitudinal differential analysis of factors
  - Factor annotation and enrichment analysis
  - Detailed evaluations of pathway activities
  - Inter-omics association analysis
  - Viral-adjusted IFN signaling analysis


### RESOURCE AVAILABILITY

All requests for information regarding reagents and resources should be directed to the lead contact and will be fulfilled by the lead contact or corresponding authors.

#### Lead contact


Further information and requests for resources and reagents should be directed to and will be fulfilled by Dr. Leying Guan.


#### Materials availability

This study did not generate new unique reagents.

#### Data and code availability

Data files are available at ImmPort under accession number SDY1760 and dbGAP accession number phs002686.v1.p1. All analysis codes have been deposited at https://bitbucket.org/kleinstein/impacc-public-code and are publicly available. DOIs are listed in the key resources table.

### EXPERIMENTAL MODEL AND STUDY PARTICIPANT DETAILS

#### IMPACC Cohort characteristics

The IMPACC cohort enrolled participants from 20 hospitals affiliated with 15 geographically distributed academic institutions across the U.S. Eligible participants were patients hospitalized with symptoms or signs consistent with COVID-19, which had SARS-CoV-2 infection confirmed by RT-PCR to remain in the study. The detailed study design and schedule for clinical data and biological sample collection were previously described (1, 2). Briefly, detailed clinical assessments and nasal swabs, blood, and endotracheal aspirates (intubated patients only) were collected within 72h of hospitalization (Visit 1) and on days 4, 7, 14, 21, 28 after hospital admission. If a participant required escalation of care or was readmitted to the hospital prior to Day 28, additional samples were collected within 24 and 96 hours of care escalation or readmission. If participants were discharged prior to day 14 or 28, attempts were made to collect limited clinical information and biologic samples on days 14 and/or 28 in outpatients. Disease severity was assessed using a 7-point ordinal scale based on degree of respiratory illness (*23*), modified from Beigel et al. (3). The sex of participants was determined via physician-reported sex at birth.

#### Cohort definition

In this study, we used 1,152 IMPACC participants with measurements for at least one of the following assays: plasma targeted proteomics, plasma global proteomics, serum proteomics, plasma global metabolomics, PBMC transcriptomics and nasal transcriptomics. The sample processing and data generation was performed in phases (subsets of samples). The participants were divided into training and test cohorts in the following manner. In a previous IMPACC manuscript, we analyzed a subset of the IMPACC cohort (enrolled through September 2020; phase 1 and 2 of the study) to perform deep immunophenotyping of COVID-19 disease using unimodal analyses (4). This subset was used as a training cohort in the present multi-modal integrative analysis to investigate the immune and molecular signatures of SARS-CoV-2 infection across tissue compartments at systems level. One participant in this subset lacked all six assays mentioned above, it was, therefore, excluded, resulting in 1,493 sample collection events from 539 participants in the training set. Unpublished data from another subset of IMPACC cohort (enrolled after September 2020; phase 3 of the study) was used as a test cohort to validate our conclusions of systems-level COVID-19 signatures from the training cohort. Few samples of the test cohort participants were included in phases 1 and 2, which were excluded from the test set, resulting in 1,584 sample collection events from 613 participants in the test set. The data for both the training and test sets were generated in the same manner to avoid any major technical biases.

### METHOD DETAILS

#### Sample processing and batch randomization

Biological sample collection and processing followed a standard protocol utilized by every participating academic institution. The complete IMPACC sample processing protocol was published previously (2). Briefly, blood samples (10 ml per time point) and nasal swabs (mid-turbinate) were collected at each specified time point, and blood was processed within 6 hours of collection. Whole blood was used to identify distinct immune cell populations and quantify changes in cell populations [cytometry by time-of-flight (CyTOF)], and peripheral blood mononuclear cells (PBMCs) were collected to measure gene expression (bulk transcriptomics; PBMC gene expression: PGX) over the course of COVID-19. Serum was used to characterize SARS-CoV-2–specific antibodies, including virus neutralization. Plasma was used for proteomics (Plasma proteomics targeted: PPT & global: PPG) and metabolomics (Plasma metabolomics global: PMG), and serum was used to measure soluble inflammatory mediators (e.g., cytokines and chemokines) using oligonucleotide-linked antibody detection (Olink) (Serum proteomics targeted: SPT). RNA from the nasal swab was used to assess SARS-CoV-2 viral load and to evaluate changes in immune-related upper airway epithelial gene expression (i.e., bulk transcriptomics; Nasal gene expression: NGX). To mitigate potential batching effects, a randomization procedure was developed to help ensure that longitudinal samples from the same individuals were run on the same plates and were randomly distributed across the plates. We stratified this randomization by disease severity (moderate versus severe) and age (younger versus older) with the representation of these strata across plates. In addition, we verified that race, ethnicity, gender, and site were well-represented across the plates.

#### Assay preparation and processing

Raw sample collection and processing was performed as defined previously (*95*) in the following sections (names in the parentheses are section names from previous work (*95*)): NGX and viral loads (Nasal viral PCR and host transcriptomics), antibody titers (Antibody correlates: titers), SPT (Serum Olink), PPG (Plasma global proteomics), PPT (Plasma targeted proteomics), PMG (Plasma global metabolomics), whole blood CyTOF (Blood CyTOF), and PGX (Peripheral Blood Mononuclear Cell transcriptomics). Additional details on PPG sample processing were recently published (*96*). The experimental methods for these assays follow the same protocols as described in our previous work [Core Assay] and are included below for the sake of completeness.

##### *Nasal host gene expression (NGX)*

Inferior nasal turbinate swabs were placed in 1mL of Zymo-DNA/RNA shield reagent (Zymo Research). RNA was extracted from 250 µL of sample and eluted into a volume of 50ul using the KingFisher Flex sample purification system (ThermoFisher) and the quick DNA-RNA MagBead kit (Zymo Research) following the manufacturer’s instructions. Each sample was extracted twice in parallel. The 2 eluted RNA samples were pooled and aliquoted into 20 µL aliquots using a Rainin Liquidator 96 pipettor for downstream RT-qPCR and RNA-sequencing.

From each nasal RNA sample, 10ul was aliquoted to a library construction plate using the Perkin Elmer Janus Workstation (Perkin Elmer, Janus II). Ribosomal depletion, cDNA synthesis, and library construction steps were performed using the Total Stranded RNA Prep with Ribo-Zero Plus kit, following the manufacturer's instructions (Illumina). All steps were automated on the Perkin Elmer Sciclone NGSx Workstation to reduce batch-to-batch variability and increase sample throughput. Final cDNA libraries were quantified using the Quant-it dsDNA High Sensitivity assay, and library insert size distribution was checked using a fragment analyzer (Advanced Analytical; kit ID DNF474). Samples, where adapter dimers constituted more than 4% of the electropherogram area, were failed before sequencing. Technical controls (K562, Thermo Fisher Scientific, cat# AM7832) were compared to expected results to ensure that batch to batch variability was minimized. Successful libraries were normalized to 10nM for sequencing.

Barcoded libraries were pooled using liquid handling robotics prior to loading. Massively parallel sequencing-by-synthesis with fluorescently labeled reversibly terminating nucleotides was carried out on the NovaSeq 6000 sequencer using S4 flowcells with a target depth of 50 million 100 base-pair paired-end reads per sample (25 million read pairs).

*Nasal Viral RT-qPCR (viral load)*

The RNA samples extracted from inferior nasal turbinate swabs (as described above) were used for this assay. Master mixes containing nuclease-Free water, combined primer/probe mixes, and One-Step RT-qPCR ToughMix (Quantabio) were prepared on ice, and 15 µL was dispensed in each well of a 384-reaction plate (Thermofisher) CoV-2 was quantitated using the CDC qRT-PCR assay (primers and probes from IDT). Briefly, this comprises two reactions targeting the CoV-2 nucleocapsid gene (N1 and N2) and one reaction targeting RPP30 (RP). Each batch included positive controls of plasmids containing N1/N2 and RP target sequence (2019-nCoV_N_Positive Control and Hs_RPP30 Positive Control, IDT) to allow quantitation of each transcript. Primer/probe sequences were: 2019-nCOV_N1-F GAC CCC AAA ATC AGC GAA AT, 2019-nCOV_N1-R TCT GGT TAC TGC CAG TTG AAT CTG, 2019-nCOV_N1-P ACC CCG CAT TAC GTT TGG TGG ACC, 2019-nCOV_N2-F TTA CAA ACA TTG GCC GCA AA, 2019-nCOV_N2-R GCG CGA CAT TCC GAA GAA, 2019-nCOV_N2-P ACA ATT TGC CCC CAG CGC TTC AG, RP-F AGA TTT GGA CCT GCG AGC G, RP-R GAG CGG CTG TCT CCA CAA GT and RP-P TTC TGA CCT GAA GGC TCT GCG CG. After RNA extracts were gently vortexed and added 5 µL per sample. Plates were centrifuged for 30 s at 500 × g, 4C. Quantitative polymerase chain reaction was performed using a Quantstudio5 (Thermo Fisher) with cycling conditions:1 cycle 10 min at 50°C, followed by 3 min at 95°C, 45 cycles 3 s at 95°C, followed by 30 s at 55.0°C.

*Antibody titers*

Antibody levels against the recombinant receptor-binding domain (RBD) and full-length spike were measured using a research-grade ELISA (5, 6). Briefly, samples were heat-inactivated at 56°C for 1 h. 96-well plates (Thermo Fisher Lot # 4199147) were coated with 50 µL/well of RBD or spike proteins at 2 µg/mL concentration in phosphate-buffered saline (PBS; Gibco lot #

2388102) and incubated overnight at 4°C. Plates were washed 3× in an automatic plate washer (BioTek) with PBS 0.01% Tween 20 (Fisher Scientific, Cat#BP337-100, TPBS) and blocked for 1 h with 200 µL/well of 3% non-fat dry milk (Cat#AB10109-01000) prepared in TPBS. Serum samples were serially diluted (3-fold starting at 1:80 dilution) in 1% non-fat dry milk in TPBS. The blocking solution was removed, and 100 µL/well of serially diluted samples were added to the plates and incubated for 2h at 20°C. Plates were washed 3× with TPBS, and 50 µL/well of the corresponding secondary antibody, prepared in 1% non-fat dry milk in TPBS, were added for 1h at RT: Anti-human IgG (Fc specific)-Peroxidase antibody produced in goat (Sigma-Aldrich Cat#A0170); Goat anti-human IgM-HRP (SouthernBiotech Cat#2020–05); Anti-human IgA (α-chain specific)-Peroxidase antibody produced in goat (Sigma-Aldrich Cat#A0295). Plates were washed 3× with TPBS, and 100 µL/well of peroxidase substrate (SigmaFAST o-phenylenediamine dihydrochloride, Sigma-Aldrich Cat#P9187) were added for 10 min 50 µL/well of 3M hydrochloric acid (HCl, Thermo Fisher Scientific, Cat#S25856) was added to stop the reaction. Optical density (OD) was measured in a Synergy 4 (BioTek) plate reader at 490 nm. The area under the curve was calculated, considering 0.15 OD as the cutoff. Data were analyzed using Graphpad Prism 9.

*Serum proteomics targeted (Olink; SPT)*

Study samples were assayed in plate batch layouts following a centralized randomized scheme that we described previously (4). Three samples (IMPACC_Serum, IMPACC_Plasma, and IMPACC_Plasma_Stim) were used as IMPACC inter-plate references (Reference samples) in every plate. All samples (participant sera and reference) were subjected to PEA (Olink) multiplex assay Inflammatory panel (Olink Bioscience, Uppsala, Sweden), according to the manufacturer's instructions. This inflammatory panel included 92 proteins associated with human inflammatory conditions. An incubation master mix containing pairs of oligonucleotide-labeled antibodies to each protein was added to the samples and incubated for 16 h at 4°C. Each protein was targeted with two different epitope-specific antibodies, increasing the assay's specificity. The presence of the target protein in the sample brought the partner probes in close proximity, allowing the formation of a double-strand oligonucleotide polymerase chain reaction (PCR) target. On the following day, the extension master mix in the sample initiated the specific target sequences to be detected and generated amplicons using PCR in 96 well plates. For the detection of the specific protein, Dynamic array integrated fluidic Circuit (IFC) 96 × 96 chip was primed, loaded with 92 protein-specific primers, and mixed with sample amplicons, including three inter-plate controls (IPS) and three negative controls (NC). Real-time microfluidic qPCR was performed in Biomark (Fluidigm, San Francisco, CA) for the target protein quantification.

*Plasma proteomics global (PPG)*

Fifty microliters of neat plasma samples were diluted with 450 µL of water, and 25 µL of perchloric acid was added (7). After vigorous agitation, the suspension was kept at −20°C for 15 min, then centrifuged for 60 min (4°C, 3200 ×g). 390 µL of the supernatant was mixed with 40 µL of 1% trifluoroacetic acid and loaded onto a µSPE HLB plate, previously conditioned once with 300 µL methanol and twice with 500 µL of 0.1% trifluoroacetic acid. Proteins were eluted from the µSPE HLB plate with 100 µL of 90% acetonitrile and 0.1% trifluoroacetic acid. After elution, the samples were dried with a Speedvac, resuspended with 35 µL of 50 mM ammonium bicarbonate, and digested with 10 µL trypsin (500 ng) overnight at 37°C. Digestion was stopped by the addition of 5 µL 10% formic acid. The samples were stored at −80°C before LC/MS analysis. Two microliters of tryptic peptides were loaded onto Evotips and analyzed using an Evosep ONE liquid chromatography system (EVOSEP, Odense, Denmark) connected to a timsTOF Pro mass spectrometer (Bruker Daltonics, Billerica, MA, USA). The Evosep ONE was set to 60 samples per day, and the mass spectrometer was operated in DDA-PASEF mode. DDA-PASEF parameters were set as follows: m/z range 100–1700, the mobility (1/K0) range was set to 0.70–1.45 Vs./cm2, and the accumulation time was set to 100 ms.

*Plasma proteomics targeted (PPT)*

All chemicals and reagents were purchased at the highest purities available. Solvents used in this study were LC/MS grade and were purchased from Fisher Chemicals (Thermo Fisher Scientific). Briefly, a volume of 10 µL of 10-fold diluted plasma was mixed with 60 µL of urea buffer (8M urea in 50 mM ammonium bicarbonate, Sigma Aldrich) and 15 µL of dithiothreitol buffer (DTT, 50 mM in urea buffer, Sigma Aldrich) before incubated 30 min on a thermomixer (800 rpm, room temperature). The samples were alkylated using iodoacetamide buffer (IAA, 375 mM in urea buffer, Sigma Aldrich) and incubated for 30 min (800 rpm, room temperature, and dark). A volume of 10 µL of DTT buffer was added to quench the alkylation. The samples were transferred to the SP3 beads mixture (Sera-Mag SpeedBeads, 1:1 v/v, GE Healthcare) previously washed with HPLC water (scale 1:10 protein to beads). Then a volume of 150 µL of absolute ethanol (Supelco) was added, and the mix was incubated for 15 min on a thermomixer (1,000 rpm at room temperature). The samples were placed on the magnetic rack, and the clear supernatant was removed. The beads were washed in three cycles in 200 µL of 80% ethanol. After the final washing step, the samples were trypsinized using 100 µL of trypsin buffer (Promega, 20 µg/mL in 50 mM ammonium bicarbonate) and placed on a thermomixer (1,000 rpm, 2 h, 37°). After digestion, the samples were centrifuged to pulldown the liquid and placed on a magnetic rack to collect the supernatant and were then acidified with 2% v/v formic acid in HPLC water (Sigma Aldrich). The C18 cleanup was performed using a 96-well MACROSPIN C18 plate (TARGA, The NestGroup Inc.), and the tryptic peptides were eluted off the C18 particles using 40% ACN/0.1% FA. The samples were dried and stored at −20°C until LC/MS analysis (8). The samples were analyzed using an LC system (Nexera Mikros, Shimadzu) equipped with a Capillary C18 column (0.2 × 100mm, 2.7um particle diameter, Shimadzu) coupled online to an 8060 triple quadrupole mass spectrometer instrument (Shimadzu). From each sample, 1 µg peptide quantity was separated using a non-linear gradient over a 15-min run time operated at 10 µL/min (5% solvent B for 0.2 min; 5 to 40%B for 10.3 min; 85%B for 1.5 min and 5% for 3 min). The final scheduling method was performed using the following parameters: 1.2 s of maximum loop time with minimum dwell time of 2 msec and pause time of 1 msec, Q1 and Q3 resolution set at the ‘unit’ level.

*Plasma metabolomics global (PMG)*

Plasma metabolite profiling was conducted by Metabolon using in-house standards (9, 10). The samples were divided into randomized sample batches, extracted, and prepared for analysis using Metabolon's solvent extraction method (Evans, 2008). Recovery standards were added to the first step in the extraction process to ensure proper quality control. Protein was removed by methanol precipitation under vigorous shaking for 2 min (Glen Mills GenoGrinder 2000) and then by centrifugation. The supernatants were divided into five fractions: two for analysis by two separate reverse phases (RP)/UPLC-MS/MS methods with positive ion mode electrospray ionization (ESI); one for analysis by RP/UPLC-MS/MS with negative ion mode ESI; one for analysis by HILIC/UPLC-MS/MS with negative ion mode ESI; and one sample was reserved for backup analysis using Waters ACQUITY ultra-performance liquid chromatography (UPLC) and a Thermo Scientific Q-Exactive high resolution/accurate mass spectrometer interfaced with a heated electrospray ionization (HESI-II) source and Orbitrap mass analyzer operated at 35,000 mass resolution. Metabolites were identified by comparison to Metabolon library entries of standard metabolites (9) based on three criteria: retention index (RI) within a narrow RI window of the proposed identification; accurate mass match to the library ±10 ppm; and the MS/MS forward and reverse scores between the experimental data and authentic standards. Compounds were categorized according to reporting standards set by the Chemical Analysis Working Group of the Metabolomics Standards Initiative (11–13), and appropriate orthogonal analytical techniques were applied to the metabolite of interest and a chemical reference standard. Metabolites were reported that had their corresponding accurate mass confirmed via MS with retention index, chemical, and composition ID.

*Blood CyTOF*

Samples from a given batch were acquired on the Fluidigm Helios mass cytometer in multiple acquisitions. The PROT-1 fixed whole blood samples were processed in batches of 20 samples. Due to sample quality issues, some samples remained pink or red after the barcoding step; those samples were discarded, and the remaining samples were pooled for the remaining staining steps. After staining was completed, the pooled sample was counted and split into 2–3 subsamples to be frozen as FBS/DMSO samples stored at −80C until the day of acquisition. On the day of acquisition, the Helios instrument was tuned according to the manufacturer's software standards; if the signal of Tb159 or Tm169 from the Fluidigm Tuning Solution was more than 10% lower than previous days, the process was repeated until the margin was achieved. The final Tuning results were exported as a CSV from the software for the record.

One FBS/DMSO subsample was thawed, washed once with Fluidigm Cell Staining Buffer, and then counted on a Bio-Rad TC20 cell counter. If necessary, the sample was split into subsamples of 2 × 106 cells, centrifuged, and the resulting pellet was left with a minimal overlay of CSB. One CSB subsample was washed twice in MilliQ water, or Fluidigm Cell Acquisition Solution then resuspended to 7–8 x 105/mL in CAS or MilliQ containing a 10-fold dilution of Fluidigm EQ 4-Element normalization beads and acquired on the tuned Helios instrument using either the PSI or SuperSampler for sample introduction. This dilution was chosen to give approximately 250–

350 events/sec acquisition rate. The next CSB subsample or FBS/DMSO subsample was processed when the previous sample had less than 1mL of sample remaining. The instrument was cleaned with Fluidigm Wash Solution whenever clogging occurred, or approximately every 2 × 106 cell events were acquired. These cleaning steps resulted in multiple FCS files per pooled sample acquisition. Pooled samples were acquired until a total of 6 × 106 cell events had been collected, or all FBS/DMSO samples were collected, whichever occurred first. This corresponds to an average target event number of 3 × 105 events per original donor subsample.

*Peripheral blood mononuclear cell (PBMC) gene expression (PGX)*

RNA was extracted from cells (2.5 × 105 PBMCs) homogenized in 200 µL of Buffer RLT (Qiagen) and then extracted using the Quick-RNA MagBead Kit (Zymo) with DNase digestion. RNA quality was quantitated using Qubit HS RNA assays and assessed using a Fragment Analyzer (Agilent). Library preps were performed using the SMART-Seq v4 Ultra Low Input RNA Kit (Takara Bio) to synthesize full-length cDNA from an input of 10ng of RNA. After a bead-based clean-up to purify the cDNA, the Nextera XT kit was used to create libraries through a process of tagmentation and fragment amplification and appended with dual-indexed bar codes using the NexteraXT DNA Library Preparation kit (Illumina). Libraries were validated by capillary electrophoresis on a Fragment Analyzer (Agilent), pooled at equimolar concentrations, and sequenced on an Illumina NovaSeq6000 (Emory) at 100 bp, paired-end read length targeting ~25 million reads per sample. Repeated measures from a group of PBMC samples collected from healthy controls and repeated measures of a subset of IMPACC samples were used across library prep and sequencing batches to assess inter-site batch effects throughout the study. Universal Human References controls were included to assess intra-site batch variation.

### QUANTIFICATION AND STATISTICAL ANALYSIS

**Quantification:** OMIC-specific processing from raw to computable matrices

*Nasal gene expression (NGX)*

Base calls were generated in real-time on the NovaSeq6000 instrument (RTA 3.1.5). Demultiplexed, unaligned BAM files were produced by Picard (14) ExtractIlluminaBarcodes, and IlluminaBasecallsToSam were converted to FASTQ format using SamTools bam2fq (15) (v1.4). The sequence read, and base quality were checked using the Trimmomatic-toolkit (16) (v0.36.5). Reads were processed using workflows managed on the Galaxy platform. Reads were trimmed by 1 base at the 3′ end, then trimmed from both ends until base calls had a minimum quality score of at least 30. Any remaining adapter sequence was removed as well. The STAR aligner (17) (v2.4.2a) with the GRCh38 (18) reference genome and gene annotations from Ensembl release 91 (19) was used to align the trimmed reads. Gene counts were generated using HTSeq-count (20) (v0.4.1). Quality metrics were compiled from Picard (v1.134), FASTQC (21) (v0.11.3), Samtools (15) (v1.2), and HTSeq-count (v0.4.1). Failed samples were identified as median cv gene coverage >0.8 and Aligned Counts <1 million. These samples were removed from further downstream analyses.

*Serum proteomics targeted (SPT)*

Data were analyzed using Real-time PCR analysis software via the ΔΔCt method and Normalized Protein Expression (NPX) manager. NPX is calculated in three steps from the Cq-values: (i) ΔCqsample = Cqsample − Cqextensioncontrol, (ii) ΔΔCq = ΔCqsample − ΔCqinterplatecontrol, (iii) NPX = Correction factor − ΔΔCqsample. Data were normalized using internal controls in every sample, inter-plate control (IPC) and negative controls, and correction factor and expressed as Log2 scale proportional to the protein concentration. One NPX difference equals to the doubling of the protein concentration.

Batch normalization was performed to account for potential batch effects caused by re-assayed samples which were not able to adhere to the study randomization scheme or assay condition changes including those due to assay kit lot# changes or differences in study collection phases. Olink Data Analysis Normalization employed identical reference samples in all plates. NPX value for each analyte was adjusted based on the adjust factor that makes the median of all reference samples the same for all plates. Sequential steps included: 1) the reference sample the-inter-plate-median was calculated; 2) for each assay, the pairwise difference from the inter-plate median was calculated in first step 1 for each of the reference sample on all plates; 3) plate- and assay-specific differences in step 2 were used as normalization factors; and 4) plate- and assay-specific normalization factors were added from step 3 to each value for each assay and plate.

*Plasma global proteomics data processing and quality control*

All raw timsTOF data were searched on a high-performance computing environment where Fragpipe (including MSFragger, Philosopher, and IonQuant (22–25)) was run to identify and quantify peptides and protein throughout the data (26). MSFragger 3.4 was run using the standard settings without the fixed modification of carboxylmethylation and with the variable modification's oxidation and N-term acetylation. Data were scored against a human FASTA file without isoforms where SARS-COV-2 proteins were manually added. Philosopher 4.1.1 was used where PeptideProphet was used for statistical validation of identified peptides. IonQuant 1.7.17 was used for quantification, where a minimum of 1 ion was used for peptide quantification.

Genes were first filtered based on "Homo Sapiens" and "Homo sapiens OX = 9606". For each sample, the "Total intensity" column was selected. Then Genes without any values across the samples were removed. Finally, sample outliers were removed. A sample is considered an outlier if its total number of quantified proteins is more than 3 standard deviations below the mean of quantified proteins of all samples. In brief, the number of proteins quantified for each sample was calculated, and log2-transformed. Then the mean and standard deviation of quantified proteins across all samples was calculated, and any samples outside 3 standard deviations were considered an outlier and removed. Finally, a protein had to be identified and quantified in at least half of all samples to be analyzed in any of the downstream analyses. We identified 508 proteins that were present in at least 699 (50%) of the samples (out of 2109 proteins in total).

*Plasma targeted proteomics data processing*

The raw data were exported into Skyline software (27) (v20.2.1.315) for peak area and retention time refinement. The peptide intensity (average of transition pairs) and the protein abundance (average of peptide intensities) in all samples were exported from Skyline. These effects were corrected using Combat (28). The means of the peptide intensities were used for the different protein abundances, which were exported for further analysis using RStudio Pro Server.

*Plasma global metabolomics data processing and quality control*

Raw data were measured based on LC-MS peak areas proportional to feature concentration. For quality control, missing values were imputed with half the minimum detected level for a given metabolite. Metabolites with an interquartile range of 0 were excluded from the analysis, as previously described (29). All features were log-transformed, normalized then Pareto-scaled to reduce variation in fold-change differences between features (Figures S5A and S5B). After pre-processing, 5 metabolites were filtered out with zero interquartile range, yielding 1012 remaining metabolites (Figure S5C). Statistical analyses for univariate, chemometrics, and clustering analysis used in-house algorithms, R statistical packages, and MetaboAnalyst 5.0 (30, 31).

*Blood CyTOF data processing and demultiplexing*

Samples from a given batch were acquired on the Fluidigm Helios mass cytometer in multiple acquisitions. The resulting FCS files were normalized and concatenated using Fluidigm's CyTOF software. The FCS file was further cleaned using the Human Immune Monitoring Center at Mt. Sinai's internal pipeline. The pipeline removed any aberrant acquisition time windows of 3 s where the cell sampling event rate was too high or too low (2 standard deviations from the mean). EQ normalization beads that were spiked into every acquisition and used for normalization were removed, along with events that had low DNA signal intensity.

The pipeline was also used to demultiplex the cleaned and pooled FCS files into single sample files. The cosine similarity of every cell's Pd barcoding channels to every possible barcode used in a batch was calculated and then was assigned to its highest similarity barcode. Once the cell had been assigned to a sample barcode, the difference between its highest and second highest similarity scores was calculated and used as a signal-to-noise metric. Any cells with low signal-to-noise were flagged as multiplets and removed from that sample. Finally, acquisition multiplets were removed based on the Gaussian parameters Residual and Offset acquired by the Helios mass cytometer.

Cells from a single biological sample were clustered into 1000 K-means clusters. A subset of samples was then selected and manually annotated into cell types using Clustergrammer2's widget interface (https://github.com/ismms-himc/clustergrammer2) to create a training dataset (n x n matrix of cell types by median marker intensities) for each manually annotated sample.

To annotate a given sample's 1000 K-means clusters, the cosine similarity of every cluster to all possible cell types within the training datasets was calculated, and that cluster was assigned to either its highest similarity score cell type or the greatest consensus cell type across the training datasets. Finally, the cluster cell-type annotation was assigned back to the single cells within that cluster, and the number of cells was calculated for a cell type within a given single sample.

*PBMC transcriptomics data processing and quality control*

Processing and quality control was performed using an internal Snakemake workflow for RNA-Seq analysis (Github: https://github.com/yerkes-gencore/IMPACC-RNA_Seq). Reads were trimmed for adapter sequence and quality score with cutadapt v1.14112. Reads were aligned with STAR v2.4.2a (17) to a composite reference of human (GRCh38) (18) reference sequence with gene annotations from Ensembl release 91 (19) and SARS-CoV-2 (NCBI strain MN908947.3). Transcript abundance estimates were calculated internal to the STAR aligner using the algorithm of htseq-count94. Sequencing quality metrics were determined using FastQC77 (v0.11.5), alignment quality metrics with Picard tools (v2.22)93 and STAR logs and gene counts, including average quality per read > Q30, percent and absolute counts of reads uniquely mapped to annotated transcripts.

**Data preprocessing and additional quality control**

Samples included for analysis have undergone prior core internal and assay-specific quality control steps. In addition, proper procedures for quality assurance outlined previously were performed to ensure the data standards for each assay were met. The table below provides information on additional steps to prepare the data for statistical analysis.

| **Assay name** | **Sample filtering** | **Feature filtering** | **Additional batch correction** | **Missing value imputation** | **Data transformation** |
|---|---|---|---|---|---|
| PBMC and Nasal gene expression (PGX & NGX) | Passed & questionable QC | Protein coding, genes with CPM >= 1 in >5% of samples in a trajectory group, top 75% highly variable genes | removeBatcheffect from limma | N/A | Scaling |
| Blood CyTOF | Passed & questionable QC | | N/A | N/A | Normalized by total counts per sample with Log1p normalization. |
| Serum proteomics targeted (SPT) | Passed & questionable QC | N/A | N/A | Impute.knn | Scaling |
| Plasma proteomics global and targeted (PPG & PPT) | Passed & questionable QC | Removed features with > 30% missing values in training and test sets, separately | N/A | Half-min | Median normalization in linear space,<br>Log2 transformation with pseudocount of 1 and scaling |
| Plasma metabolomics global (PMG) | Passed & questionable QC | Removed non-xenobiotic features with IQR = 0 | N/A | Half-min | Pareto-Scaling |

**Table S10: Information on data preparation.** For each assay, we first filtered out samples using the sample filtering criterion and followed by a filtration on features based on the feature filtering criteria, we performed data imputation and data transformation as indicated in the table. N/A: no additional step taken. Half-min: replacing missing value using half of the minimum of observed values for the corresponding feature. Impute.knn: using impute.knn function from R package impute. Pareto-Scaling: in-house function of dividing each centered variable by the square root of the standard deviation. We evaluated the influence of potential batch effects on different assays using Principal variance component analysis (PVCA) (table S9).

The preprocessing of omic datasets was performed as defined previously (4). These processing steps are briefly described below. Samples with failed or missing QC information were removed for all assays.

*Gene expression (PBMC and Nasal) (PGX and NGX)*

We filtered for the protein-coding genes and removed lowly expressed genes (genes that do not qualify for counts per million >= 1 in more than 5% of samples in at least one outcome group (trajectory group)). We also removed the 25% least variable gene as evaluated using median absolute deviation (MAD) of log-transformed counts per million values. Finally, we transformed the count data using voom (32), performed batch-correction to remove the effects of technical variables, including phase sample processing batch (phase) and library preparation plates using removeBatchEffect of limma (33), and scaled the data.

*Serum proteomics targeted (SPT)*

The missing measurements (missing at random) were imputed using nearest neighbor averaging using impute.knn function in R package impute and scaled the data.

*Plasma Proteomics (Global and Targeted)*

Features with more than 30% missing values separately in training and test sets were removed, and the data was median normalized in linear space, imputed using the half-min approach, log-transformed using pseudocount of 1 and scaled.

*Plasma metabolomics global (PMG)*

The missing measurements (missing at random) were imputed using the half-min approach, the non-xenobiotic features with IQR = 0 were removed, and pareto-scaled the data.

*Whole Blood CyTOF*

Cellular population counts were converted to a normalized frequency by dividing by the total counts per sample then log1p transformed. Granulocytes were excluded from the total counts for non-granulocyte populations.

**Data imputation via MOFA**

Complete assay imputation for the six multi-omic assays (Fig. 1A) was performed using Multi-Omic Factor Analysis (MOFA) via the MOFA2 R package (34, 35). MOFA performs dimensionality reduction through a variational Bayesian framework, modeling high-dimensional multi-omic assays as a product of low-dimensional factor loadings and scores with error, allowing the imputation of entire missing biological assays from the loadings and scores. Imputation was performed on the preprocessed training and test datasets separately to prevent any imputation-wise association between them.

The effects of MOFA imputation were explored by constructing separate composite regression models for samples missing 0, 1, or 2 assays (fig. S1J). All three models were then tested using the severity task (TG1 vs. TG2/TG3 vs. TG4/TG5) as described in the manuscript and achieved comparable results (fig. S1K).

### Multi-omics factor construction via MCIA

Factors were constructed from the preprocessed training multi-omic dataset using Multiple Co-Inertia Analysis (MCIA), which projects data into low-dimensional factors that maximize covariance between each omic and the global data matrix (36). MCIA has shown strong performance in comparison to other unsupervised joint dimensionality reduction methods in multi-omics benchmark comparisons (37, 38). Prior to implementing MCIA, the multi-omics data were standardized using a centered row profile (36), followed by block-level variance normalization (39). Implementation was performed using the mogsa R Bioconductor package (40), with custom modification of the deflation step to constrain global (factor) scores to be orthogonal. MCIA factor scores for preprocessed testing multi-omic samples were calculated by generating block and global scores using coefficients derived from the pretrained MCIA model.

### Hierarchical classification model construction and evaluation

Many multi-omic signatures have been found to be associated with disease severity. However, it has also been noticed that mortality sometimes does not lead to immunological variation along the direction of increased severity (41). To capture the signatures indicative of a more severe disease course while accounting for abrupt changes happening for the mortality group, we built a composite prediction model combining a global ordinal regression with a sub-model separating the two most severe groups TG4 and TG5. More specifically, we first fit an ordinal regression model grouping TG4 and TG5 together using the ordinalNet R package (42):

$$P(y \in \cup_{l \geq k} G_l) = \frac{1}{1 + exp\,(-x^{\top}\beta - \alpha_k)},\ k = 1,2,3$$

where G1= {1}, G2= {2,3}, G3= {4,5}. Then, we further fit a logistic regression model to separate TG4 and TG5 using the glmnet R package (43):

$$P(y = 4) = P(y \in G_3)P(y = 4|y \in G_3) = P(y \in G_3)\frac{1}{1+exp\,(x^{\top}\theta+\nu)},$$

$$P(y = 5) = P(y \in G_3)P(y = 5|y \in G_3|) = P(y \in G_3)\frac{1}{1+exp\,(-x^{\top}\theta-\nu)}.$$

Both included Lasso penalties to reduce the influence from nuisance features. The Lasso penalty is tuned using a 10-fold cross-validation, which is recommended to account for both the training efficiency and model variability (44). When compared it to the unstructured multinomial regression model, this structured composite model is more interpretable since the model is characterized by two coefficients: 1) the coefficient '$\beta$' measuring the global trend of being severe, and 2) the coefficient '$\theta$' identifying the high-risk sub-population among severe patients. Despite being simpler, this composite model provides similar performance to the multinomial logistic regression in our analyses (fig. S1E). Finally, we opted to combine TG2 and TG3 together for both classifiers due to poor performance when separating TG2 from TG3 in the training cohort (AUROC=0.558, fig. S1G).

#### MCIA model construction and rank selection

Composite regression models were constructed using different ranks of MCIA multi-omic factors from the grid (2, 3, …., 10, 15, 20, 25, 30, 40) using a 50-folds (nested) cross-validation on the training cohort as shown steps 1-2 of the figure below. Minimum deviance loss was achieved for the predictive model with 7 multi-omic MCIA factors (fig. S1F) which we established as the MCIA model. The MCIA model was then used on both the training and testing cohorts to achieve predictions on the testing cohort as shown in step 3 below.

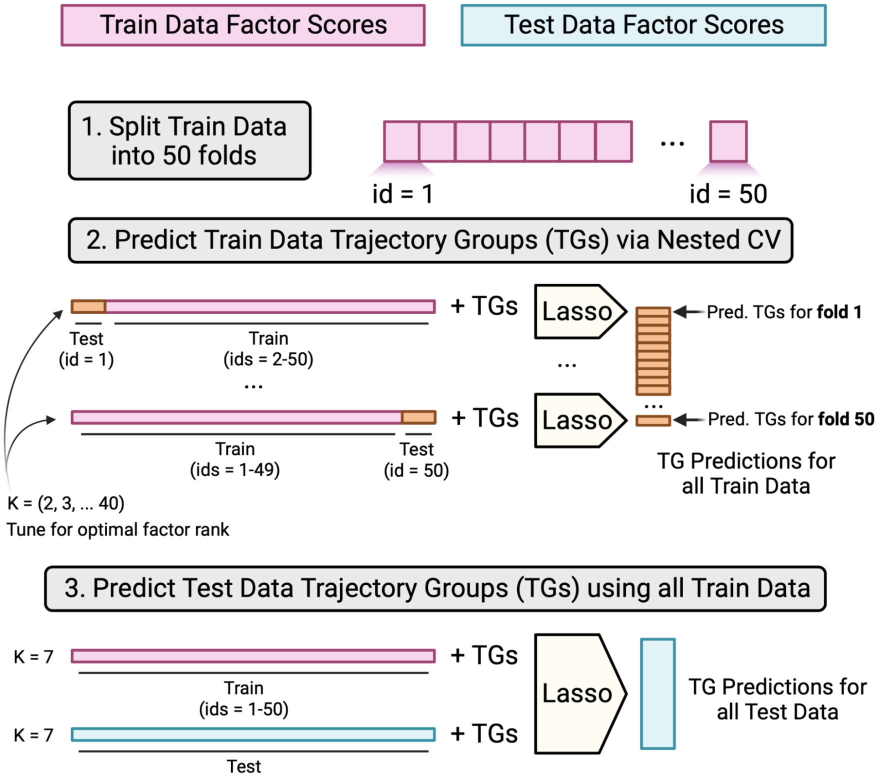


#### Clinical model construction

A composite regression model utilizing baseline clinical measurements, denoted as the clinical model, was constructed for comparison with the MCIA model. The clinical model features consisted of sex, BMI, age, ethnicity, and race as well as various baseline laboratory values and comorbidities (fig. S1A). We opted to not include the baseline respiratory status into the model as they were utilized in the construction trajectory groups (*5*).

#### Ensemble (MCIA + clinical) model construction

We considered a simple ensemble of the MCIA model and the clinical model by weighting their predictions where the weights are chosen based on the cross-validation prediction. More specifically, let ${p_i}^{mcia}$ and ${p_i}^{clin}$ denote the prediction of severity (TG4|TG5), we consider the following ensemble prediction for the severity task

$$log \frac{p_i^{ensemble}}{1-p_i^{ensemble}} \leftarrow \alpha_{mcia}\, log \frac{p_i^{mcia}}{1-p_i^{mcia}} + \alpha_{clin}\, log \frac{p_i^{clin}}{1-p_i^{clin}},$$

where $\alpha_{mcia}$ and $\alpha_{clin}$ are chosen by considering an ordinal regression model with response being the TGs and features being log odds of the nested cross-validation prediction version of $p_i^{mcia}$ and $p_i^{clin}$ on the training cohort.

Similarly, we consider the weighted prediction for the mortality task where the weights are chosen by considering an ordinal regression model with response being the TG5 or TG4 on the among critical illness, and features being log odds of the nested cross-validation prediction for TG5 in the mortality task.

**Model exploration and comparisons via training cohort cross-validation**

Both MCIA and ensemble models improved over the clinical model on the training data based on cross-validation (fig. S1B) and conveyed a similar message as the results from the test cohort (Fig. 1C). Additional predictive models were constructed and assessed to the MCIA model (fig. S1, E, H, and I). Groups of constructed models are described below with results sharing the corresponding title. Model performance was measured using both the severity (TG1 vs. TG2/TG3 vs. TG4/TG5) and mortality (TG4 vs. TG5) tasks as described in the manuscript (Fig. 1A).

*Multi-omics vs. Single-omics using Concatenation*

The MCIA model (MCIA) was compared against the following predictive models: PPT, PPG, SPT, PMG, NGX, and PGX – six separate composite regression models using concatenated analytes from each of the 6 multi-omic assays (without performing dimensionality reduction), All Assays (Concat.) - A composite regression model using all concatenated 27,320 analytes as predictors. MCIA (Multinom.) - A multinomial regression model using the 7 MCIA multi-omics factors rather than the composite classifier framework (fig. S1E).

*Multi-omics vs. Single-omics using MCIA*

The following models were constructed by using MCIA per-block construction on each assay individually (rather than together): PPT, PPG, SPT, PMG, NGX, and PGX. The MCIA model was shown to emphasize the most predictive assays for each prediction task (fig. S1H).

*MCIA Factors vs. Literature*

A systematic literature review was performed, searching for biomarkers in the context of COVID-19 severity. Publications were manually assessed for suitability, considering time of biospecimen collection, hospitalization, comparator groups and molecular data types to be as similar to the IMPACC study as possible. Three articles were found as viable candidates, each one containing a molecular signature associated with COVID-19 severity (45–47). Each molecular biomarker was matched to the corresponding IMPACC collected analyte and subsequently utilized to train a separate composite regression model as stated above (fig. S1I).

**MCIA separates TG groups with aggregated predictions**

To further explore the MCIA model predictions, we combined the results from the severity and mortality tasks to achieve four unique classes: TG1, TG2/TG3, TG4 and TG5. We chose to keep

TG2 and TG3 binned together after observing a low AUROC when trying to separate them on the training cohort (fig. S1F). The four classes were utilized in a multi-class AUROC prediction (one versus all) framework. The clinical model was outperformed by the MCIA and ensemble models for every class (fig. S1C).

Finally, we combined the classifier-assigned probabilities from the severity and mortality tasks together to embed participants (fig. S1D). Grouping participants by their TGs revealed a gradual shift in both the severity (x-axis) and mortality (y-axis) directions, with severity and mortality scores increasing from TG1 < TG2/TG3 < TG4 < TG5.

**MCIA predicted risk and hypothesis testing conditional on baseline**

To investigate if MCIA model provided additional about clinical trajectory groups conditional on participant's ordinal scale for baseline respiratory, we constructed a predicted severity from the MCIA model and test if the predicted risk is significant in following ordinal regression model:

$$\log P(TG \geq k) = \frac{1}{1+\exp\left(-age*\beta_{age}-sex*\beta_{sex}-baseline*\beta_{base}-risk*\beta_{risk}-\alpha\right)},$$

where "baseline" refers to the score for baseline respiratory status, and "risk" refers to the predicted risk defined as below:

$$predicted\ risk = \sqrt{P(TG=5) * \ P(TG \in \{4,5\})},$$

which assigns higher risk to patients more likely being in TG4 and TG5 from our MCIA model and shows higher aversion for TG5 compared to TG4. We considered the predicted risk to have significant contribution to the clinical trajectory prediction conditional on the baseline respiratory status if the p-value is smaller than 0.05. We consider this test separately for test-cohort patients with moderately impaired baseline respiratory status (baseline score in [3,4]) and for test-cohort patients with severely impaired baseline respiratory status (baseline score in [5,6]) to examine if the predicted risk if conditionally informative about future clinical trajectory given different incoming status.

### Baseline and longitudinal differential analysis of factors

We performed a cumulative link mixed effect analysis using MCIA factor scores from baseline samples and investigated (1) if there is an ordinal trend from TG1 to TG5 and (2) if any pair of groups exhibit significant differences. Enrollment sites were assigned as a random effect and age group (split across five quantiles ([18,35], [36,51], [52,66], [67,81], [82,96])) and sex were included as fixed effects. We tested for the ordinal trend with the clmm function from the R package ordinal (48) (v 2019.12-10) and pairwise difference with the lmer function from the R package lme4 (49) (v 1.1-27.1). We identified significant MCIA factors whose adjusted p values (Benjamini-Hochberg Procedure (50)) are below 0.05 for the ordinal trend or for the pairwise comparisons between TG4 and TG5. Significant factors can potentially be used for separating clinical groups at hospital admission.

We next moved to longitudinal analysis for scheduled visits (Visits 1-6) to identify MCIA factors whose scores differ for different clinical groups from the training set ranging from day 0 (admission) to day 41. We performed a linear mixed modeling analysis for the factor levels using the lme function from the R package nlme (51) (v 3.1-148) and a mixed generalized additive modeling analysis and modeled the factor levels as a smooth function (cubic regression) of admission time using the R package gamm4 (52) (v 0.2-6) function gamm4. For each pair of groups, we tested if the two groups have different longitudinal trends for the MCIA factors after including the participant ID and enrollment site as random effects while sex and age group as fixed effects. We claimed significance when the adjusted p-value is below 0.05, and significant factors could indicate interesting molecular dynamics across clinical groups:

factor ~ s(admission date, bs = cr) + s(admission date, bs = cr, by =TG) + TG + sex + age+ (1|enrollment_site/participant id).

Here, s(., bs=cr) indicates we are using a smooth spline to model the overall kinetics over admission dates, for pooled participants and each TG group (by=TG), and the term (1|enrollment site/participant id) indicates a nested random effect by participant id and enrollment site.

The testing results for baseline (visit) and longitudinal analysis for 7 selected factors are shown in table s4.

**Factor annotation and enrichment analysis**

*Databases*: Publicly available knowledge databases were used for functional enrichment analysis, with details listed below.

- KEGG: A list of KEGG pathways and their corresponding mappings with genes and metabolites were extracted using the KEGG REST API (Release 102.0). KEGG pathways were used for gene/protein/metabolite annotations.
- Hallmark: Hallmark pathways are extracted from MsigDB database (Homo sapien) using msigdbr (version 7.5.1). Hallmark pathways were used for gene and protein annotations.
- Subpathway: Many metabolites in the plasma global metabolomics lack KEGG id correspondence. Hence, when we encountered difficulty using KEGG pathways, we used the subpathway database provided by Metabolon for metabolite annotation (9, 53, 54).
- Immunoglobulin gene sets: The GO terms with “IMMUNOGLOBULIN” in the name were extracted from MSigDB (c5 v7.5.1)
- C3 was extracted from MSigDB (C3 v7.5.1): for transcription factor and miRNA gene sets functional enrichment.
- Ligand-receptor signaling-based gene sets were constructed from the ligand/receptor and intracellular signaling reported in Omnipath, with ligand’s gene sets being composed of all genes within one connection of their receptors.
- A list of IFN inhibitors was extracted from *Porritt and Herzog* (55) for enrichment and generation of composite scores of these genes for baseline and longitudinal analysis.

*Enrichment analysis using high-contribution features with mHG:*

Enrichment of the factors was conducted using the minimum hypergeometric test (mHG) from the R package mHG (56) (v 1.1) using the gene, proteins and metabolite sets from the Hallmark, KEGG (downloaded on August 04, 2022), and C3 on high-contribution features of a factor. The joint p-value combines enrichment p-values from assays which contained non-zero features in each pathway using the Cauchy combination test, which is known to be robust to the dependence structure underlying p-values to be combined (57). The selection of an appropriate background set of features is crucial for the accurate enrichment analysis. We used only those features as background that were measured in our assays. Since the serum Olink (SPT) included only 92 proteins in total, which were preselected for their function in inflammation, the enrichment over the background of 92 features did not yield many significant pathways as expected. Therefore, to capture meaningful signal from the SPT, we also performed a direct comparison of the SPT features with features and pathways of other datasets using an 'inter-omics association analysis' (see below).

*Selection of enriched pathways of severity factor for further consideration*

The pathway databases are comprised of many broad and redundant biological pathways with highly overlapping genesets (58). We obtained significant enrichment for a large number of pathways, many of which represented redundant biological function. To handle this redundancy, we first collected biological pathways that were significantly enriched (adj.p<0.1) in each assay in the severity factor, and we clustered them based on the number of shared leading-edge features and visualized using enrichment map plot (fig. S2A). We observed main four major functional categories: inflammation, T-cell activity, cell death, and dysregulated metabolism of essential amino acids (fig. S2, A and B, table s5). We prioritized the pathways based on the strongest joint p-values (as described above) and selected the representative pathways from each of the four functional categories that reflected specialized biological functions and displayed most significant aggregated p-values. We favored the pathways with significant enrichment in at least two omics as representatives from each functional category, except for "Th1 and Th2 cell differentiation" and "T-cell receptor signaling pathway" in T-cell activity and "Tryptophan metabolism" in dysregulated metabolism categories. (Fig 3C), because the features of some of these pathways are not expected to be highly represented in other datasets, for example, metabolic features are present only in PMG and the genes encoding the enzymes involved in those metabolic pathways are not very well defined in existing knowledge bases. Similarly, TCR-signaling genes are expected to be enriched in the gene expression datasets compared to the plasma/serum proteomics datasets.

For the pathways enriched in the mortality factor, we further selected only those pathways that separated TG4 and TG5 at baseline because the dominant signal that the mortality factor captures is related to the separation of these two groups at baseline (table s7 for all enriched pathways and table s8 for pathways that show separation between TG4 and TG5, p.val < 0.05 ). Transcriptomics pathways including Influenza A, Epstein-Barr virus infection, Hepatitis C, Measles and Herpes simplex virus 1 infection are subsets of the listed interferon signaling pathways shown in Fig. 5C, hence omitted from the main Figure.

*TG4|5 filtered enrichment:*

The mHG test allows for flexible filtering of the ordered feature list before passing to the MHG test function. Apart from the plain enrichment analysis, we further considered a TG4|5 filtered enrichment analysis for each factor where we kept only high-contribution metabolites and soluble proteins who can separate TG4 and TG5 at baseline (training samples only) with p.val <=0.05 and performed MHG test using the filtered list (table s6 and table s8). This helped us avoid overlooking important functions related to the early separation of TG4 and TG5 due to the overwhelming signals from temporal kinetics and related to the overall disease severity (from TG1 to TG5).

*Cell-type enrichment analysis via GSEA:*

Due to the limited size of overlaps between cell markers and high-contribution feature, Gene Set Enrichment Analysis (GSEA) for cell type enrichment of the PBMC and nasal transcriptomics was conducted using gene sets from Nakaya H. et al. (59) and a combined gene set of Ziegler et al. (60) with the neutrophil and eosinophil marker genes from Ordovas-Montanes et al. (61) respectively.

**Detailed evaluations of pathway activities**

For pathways of interest, we performed mixed generalized additive modeling analysis and modeled the pathway activity as a smooth function of admission time using the R package gamm4 (52) (v 0.2-6) function gamm4. We included participant ID and enrollment site as random effects while sex and age quintile as fixed effects, as we have done in the factor longitudinal test:

pathway_activity ~ s(admission date, bs = cr) + s(dmission date, bs = cr, by =TG) + TG + sex + age+(1|enrollment site/participant id)

For the T cell pathway analysis in Fig. S3C, we calculated a sum of PBMC transcriptomic levels of all features in the indicated KEGG pathway and used our longitudinal modal analysis (as described above) while using T cell frequencies measured in whole blood CyTOF (combined across all T cell subsets) as a covariate to adjust for the cell abundance changes:

pathway_activity ~ s(admission date, bs = cr) + s(dmission date, bs = cr, by =TG) + TG + sex + age+ CD4+T cell frequency + CD8+T cell frequency+(1|enrollment site/participant id)

**Inter-omics association analysis**

Although each factor captures co-varying patterns across omics, the factor represents global systematic changes, and does not directly guarantee local connections between a pair of high-contribution features. Additionally, the factors do not provide direct connection of individual features to biological pathways (combined expression signatures over multiple genes/proteins). We performed inter-omics association analysis to investigate the local connections between immune components from different assays. The inter-omics analysis not only assesses the direct association between two analytes or pathway functions, but it also helps to capture meaningful functional descriptions additional to the enrichment test, by linking SPT analytes directly to enriched pathways from other assays. We consider the associations between the following types (fig. S4, fig. S6C):

1. Association between metabolite pathway activities and serum targeted proteomics. Association between metabolite pathway activities and plasma protein/gene pathway activities (both nasal and PBMC).
2. Association between serum targeted proteomics and plasma protein/gene pathway activities (both nasal and PBMC).
3. Association between metabolite pathway activities/ serum targeted proteomics and whole blood cell frequencies (parent population) measured by CyTOF.

We performed steps 1-4 for high-contribution cytokines and highlighted pathways activities for each factor. We summarized assay variability captured by a factor for assays other than serum targeted proteomics (Olink) because they were of high dimensions and individual features revealed less-interpretable information than the pathway activity. In addition, we have adjusted for age, sex, enrollment site in calculating the Pearson correlations. For steps1-3, since the pathway activities and high-contribution soluble proteins were co-selected, we further controlled for visits (1-6) and clinical trajectory groups to account for global co-varying patterns due to common trends in time and variability driven by TGs. We calculated the adjusted Pearson correlation instead of using the lme model here to reduce the computational costs from calculating tens-of-thousands correlations, which offered comparable discoveries with exact p-values varying slightly.

**Viral-adjusted IFN signaling analysis**

To understand the influence of viral loads in the IFN signaling differences observed comparing TG4 and TG5, we conducted a viral-adjusted IFN signaling analysis. Since the interaction between viral loads and IFN signaling are highly non-linear, we adjust for viral loads using random forest to avoid subjective transformation.

1. Baseline comparison: We fitted a random forest model predicting baseline IFN signaling pathway levels from baseline viral loads (default parameter, keeping samples where both viral loads nasal transcriptomics and viral loads measurements), and the fitted residuals were used Wilcoxon test for comparing TG4 and TG5 and moderate illness (TG1/TG2/TG3) vs. critical illness (TG4/TG5).
2. Kinetics comparison: We fitted a random forest model predicting IFN signaling pathway levels from viral loads using all samples (default parameter, keeping samples where both viral loads nasal transcriptomics and viral loads measurements were available), and the fitted residuals were used for comparing the kinetics from TG4 and TG5 (gamm4, mixed effect modeling where participant id is modeled as the random effect).

The viral adjusted results were compared to the unadjusted results. For a fair comparison, in this analysis alone, we used the same set of samples for the unadjusted/adjusted tests where both nasal transcriptomics and viral loads were available.

**Supplemental Acknowledgements**

**#The IMPACC Network**

**National Institute of Allergy and Infectious Diseases, National Institute of Health, Bethesda, MD 20814, USA:** Patrice M. Becker, Alison D. Augustine, Steven M. Holland, Lindsey B. Rosen, Serena Lee, Tatyana Vaysman

**Clinical and Data Coordinating Center (CDCC) Precision Vaccines Program, Boston Children's Hospital, Boston, MA 02115, USA:** Al Ozonoff, Joann Diray-Arce, Jing Chen, Alvin Kho, Carly E. Milliren, Annmarie Hoch, Ana C. Chang, Kerry McEnaney, Brenda Barton, Claudia Lentucci, Maimouna D. Murphy, Mehmet Saluvan, Tanzia Shaheen, Shanshan Liu, Caitlin Syphurs, Marisa Albert, Arash Nemati Hayati, Robert Bryant, James Abraham, Sanya Thomas, Mitchell Cooney

**Benaroya Research Institute, University of Washington, Seattle, WA 98101, USA:** Matthew C. Altman, Naresh Doni Jayavelu, Scott Presnell, Bernard Kohr, Tomasz Jancsyk, Azlann Arnett

**La Jolla Institute for Immunology, La Jolla, CA 92037, USA:** Bjoern Peters, James A. Overton, Randi Vita, Kerstin Westendorf

**Knocean Inc. Toronto, ON M6P 2T3, Canada:** James A. Overton

**Precision Vaccines Program, Boston Children's Hospital, Harvard Medical School, Boston, MA 02115, USA:** Ofer Levy, Hanno Steen, Patrick van Zalm, Benoit Fatou, Kinga Smolen, Arthur Viode, Simon van Haren, Meenakshi Jha

**Brigham and Women's Hospital, Harvard Medical School, Boston, MA 02115, USA:** Lindsey R. Baden, Kevin Mendez, Jessica Lasky-Su, Alexandra Tong, Rebecca Rooks, Michael Desjardins, Amy C. Sherman, Stephen R. Walsh, Xhoi Mitre, Jessica Cauley, Xiofang Li, Bethany Evans, Christina Montesano, Jose Humberto Licona, Jonathan Krauss, Nicholas C. Issa, Jun Bai Park Chang, Natalie Izaguirre

**Metabolon Inc, Morrisville, NC 27560, USA:** Scott R. Hutton, Greg Michelotti, Kari Wong

**Prevention of Organ Failure (PROOF) Centre of Excellence, University of British Columbia, Vancouver, BC V6T 1Z3, Canada:** Scott J. Tebbutt, Casey P. Shannon

**Case Western Reserve University and University Hospitals of Cleveland, Cleveland, OH 44106, USA:** Rafick-Pierre Sekaly, Slim Fourati, Grace A. McComsey, Paul Harris, Scott Sieg, Susan Pereira Ribeiro

**Drexel University, Tower Health Hospital, Philadelphia, PA 19104, USA:** Charles B. Cairns, Elias K. Haddad, Michele A. Kutzler, Mariana Bernui, Gina Cusimano, Jennifer Connors, Kyra

Woloszczuk, David Joyner, Carolyn Edwards, Edward Lee, Edward Lin, Nataliya Melnyk, Debra L. Powell, James N. Kim, I. Michael Goonewardene, Brent Simmons, Cecilia M. Smith, Mark Martens, Brett Croen, Nicholas C. Semenza, Mathew R. Bell, Sara Furukawa, Renee McLin, George P. Tegos, Brandon Rogowski, Nathan Mege, Kristen Ulring, Pam Schearer, Judie Sheidy, Crystal Nagle

**MyOwnMed Inc., Bethesda, MD 20817, USA:** Vicki Seyfert-Margolis

**Emory School of Medicine, Atlanta, GA 30322, USA:** Nadine Rouphael, Steven E. Bosinger, Arun K. Boddapati, Greg K. Tharp, Kathryn L. Pellegrini, Brandi Johnson, Bernadine Panganiban, Christopher Huerta, Evan J. Anderson, Hady Samaha, Jonathan E. Sevransky, Laurel Bristow, Elizabeth Beagle, David Cowan, Sydney Hamilton, Thomas Hodder, Amer Bechnak, Andrew Cheng, Aneesh Mehta, Caroline R. Ciric, Christine Spainhour, Erin Carter, Erin M. Scherer, Jacob Usher, Kieffer Hellmeister, Laila Hussaini, Lauren Hewitt, Nina Mcnair, Susan Pereira Ribeiro

**Icahn School of Medicine at Mount Sinai, New York, NY 10029, USA**: Ana Fernandez-Sesma, Viviana Simon, Florian Krammer, Harm Van Bakel, Seunghee Kim-Schulze, Ana Silvia Gonzalez-Reiche, Jingjing Qi, Brian Lee, Juan Manuel Carreño, Gagandeep Singh, Ariel Raskin, Johnstone Tcheou, Zain Khalil, Adriana van de Guchte, Keith Farrugia, Zenab Khan, Geoffrey Kelly, Komal Srivastava, Lily Q. Eaker, Maria C. Bermúdez-González, Lubbertus C.F. Mulder, Katherine F. Beach, Miti Saksena, Deena Altman, Erna Kojic, Levy A. Sominsky, Arman Azad, Dominika Bielak, Hisaaki Kawabata, Temima Yellin, Miriam Fried, Leeba Sullivan, Sara Morris, Gliulio Kleiner, Daniel Stadlbauer, Jayeeta Dutta, Hui Xie, Manishkumar Patel, Kai Nie

**Immunai Inc. New York, NY 10016, USA:** Adeeb Rahman

**Oregon Health Sciences University, Portland, OR 97239, USA:** William B. Messer, Catherine L. Hough, Sarah A.R. Siegel, Peter E. Sullivan, Zhengchun Lu, Amanda E. Brunton, Matthew Strnad, Zoe L. Lyski, Felicity J. Coulter, Courtney Micheleti

**Stanford University School of Medicine, Palo Alto, CA 94305, USA:** Holden Maecker, Bali Pulendran, R. Kari C. Nadeau, Yael Rosenberg-Hasson, Michael Leipold, Natalia Sigal, Angela Rogers, Andrea Fernandes, Monali Manohar, Evan Do, Iris Chang, Alexandra S. Lee, Catherine Blish, Henna Naz Din, Jonasel Roque, Linda Geng, Maja Artandi, Mark M. Davis, Neera Ahuja, Samuel S. Yang, Sharon Chinthrajah, Thomas Hagan

**David Geffen School of Medicine at the University of California Los Angeles, Los Angeles CA 90095, USA:** Elaine F. Reed, Joanna Schaenman, Ramin Salehi-Rad, Adreanne M. Rivera,

Harry C. Pickering, Subha Sen, David Elashoff, Dawn C. Ward, Jenny Brook, Estefania Ramires-Sanchez, Megan Llamas, Claudia Perdomo, Clara E. Magyar, Jennifer Fulcher

**University of California San Francisco, San Francisco, CA 94115, USA:** David J. Erle, Carolyn S. Calfee, Carolyn M. Hendrickson, Kirsten N. Kangelaris, Viet Nguyen, Deanna Lee, Suzanna Chak, Rajani Ghale, Ana Gonzalez, Alejandra Jauregui, Carolyn Leroux, Luz Torres Altamirano, Ahmad Sadeed Rashid, Andrew Willmore, Prescott G. Woodruff, Matthew F. Krummel, Sidney Carrillo, Alyssa Ward, Charles R. Langelier, Ravi Patel, Michael Wilson, Ravi Dandekar, Bonny Alvarenga, Jayant Rajan, Walter Eckalbar, Andrew W. Schroeder, Gabriela K. Fragiadakis, Alexandra Tsitsiklis, Eran Mick, Yanedth Sanchez Guerrero, Christina Love, Lenka Maliskova, Michael Adkisson, Aleksandra Leligdowicz, Alexander Beagle, Arjun Rao, Austin Sigman, Bushra Samad, Cindy Curiel, Cole Shaw, Gayelan Tietje-Ulrich, Jeff Milush, Jonathan Singer, Joshua J. Vasquez, Kevin Tang, Legna Betancourt, Lekshmi Santhosh, Logan Pierce, Maria Tecero Paz, Michael Matthay, Neeta Thakur, Nicklaus Rodriguez, Nicole Sutter, Norman Jones, Pratik Sinha, Priya Prasad, Raphael Lota, Sadeed Rashid, Saurabh Asthana, Sharvari Bhide, Tasha Lea, Yumiko Abe-Jones

**Yale School of Medicine, New Haven, CT 06510, USA:** David A. Hafler, Ruth R. Montgomery, Albert C. Shaw, Steven H. Kleinstein, Jeremy P. Gygi, Shrikant Pawar, Anna Konstorum, Ernie Chen, Chris Cotsapas, Xiaomei Wang, Leqi Xu, Charles Dela Cruz, Akiko Iwasaki, Subhasis Mohanty, Allison Nelson, Yujiao Zhao, Shelli Farhadian, Hiromitsu Asashima, Omkar Chaudhary, Andreas Coppi, John Fournier, M. Catherine Muenker, Allison Nelson, Khadir Raddassi, Michael Rainone, William Ruff, Syim Salahuddin, Wade L. Shulz, Pavithra Vijayakumar, Haowei Wang, Esio Wunder Jr., H. Patrick Young, Albert I. Ko, Xiomei Wang

**Yale School of Public Health, New Haven, CT 06510, USA:** Denise Esserman, Leying Guan, Anderson Brito, Jessica Rothman, Nathan D. Grubaugh

**Baylor College of Medicine and the Center for Translational Research on Inflammatory Diseases, Houston, TX 77030, USA:** David B. Corry, Farrah Kheradmand, Li-Zhen Song, Ebony Nelson

**Oklahoma University Health Sciences Center, Oklahoma City, OK 73104, USA:** Jordan P. Metcalf, Nelson I. Agudelo Higuita, Lauren A. Sinko, J. Leland Booth, Douglas A. Drevets, Brent R. Brown

**University of Arizona, Tucson AZ 85721, USA:** Monica Kraft, Chris Bime, Jarrod Mosier, Heidi Erickson, Ron Schunk, Hiroki Kimura, Michelle Conway, Dave Francisco, Allyson Molzahn, Connie Cathleen Wilson, Ron Schunk, Trina Hughes, Bianca Sierra

**University of Florida, Gainesville, FL 32611, USA:** Mark A. Atkinson, Scott C. Brakenridge, Ricardo F. Ungaro, Brittany Roth Manning, Lyle Moldawer

**University of Florida, Jacksonville, FL 32218, USA:** Jordan Oberhaus, Faheem W. Guirgis

**University of South Florida, Tampa FL 33620, USA:** Brittney Borresen, Matthew L. Anderson

**University of Texas, Austin, TX 78712, USA:** Lauren I. R. Ehrlich, Esther Melamed, Cole Maguire, Dennis Wylie, Justin F. Rousseau, Kerin C. Hurley, Janelle N. Geltman, Nadia Siles, Jacob E. Rogers.

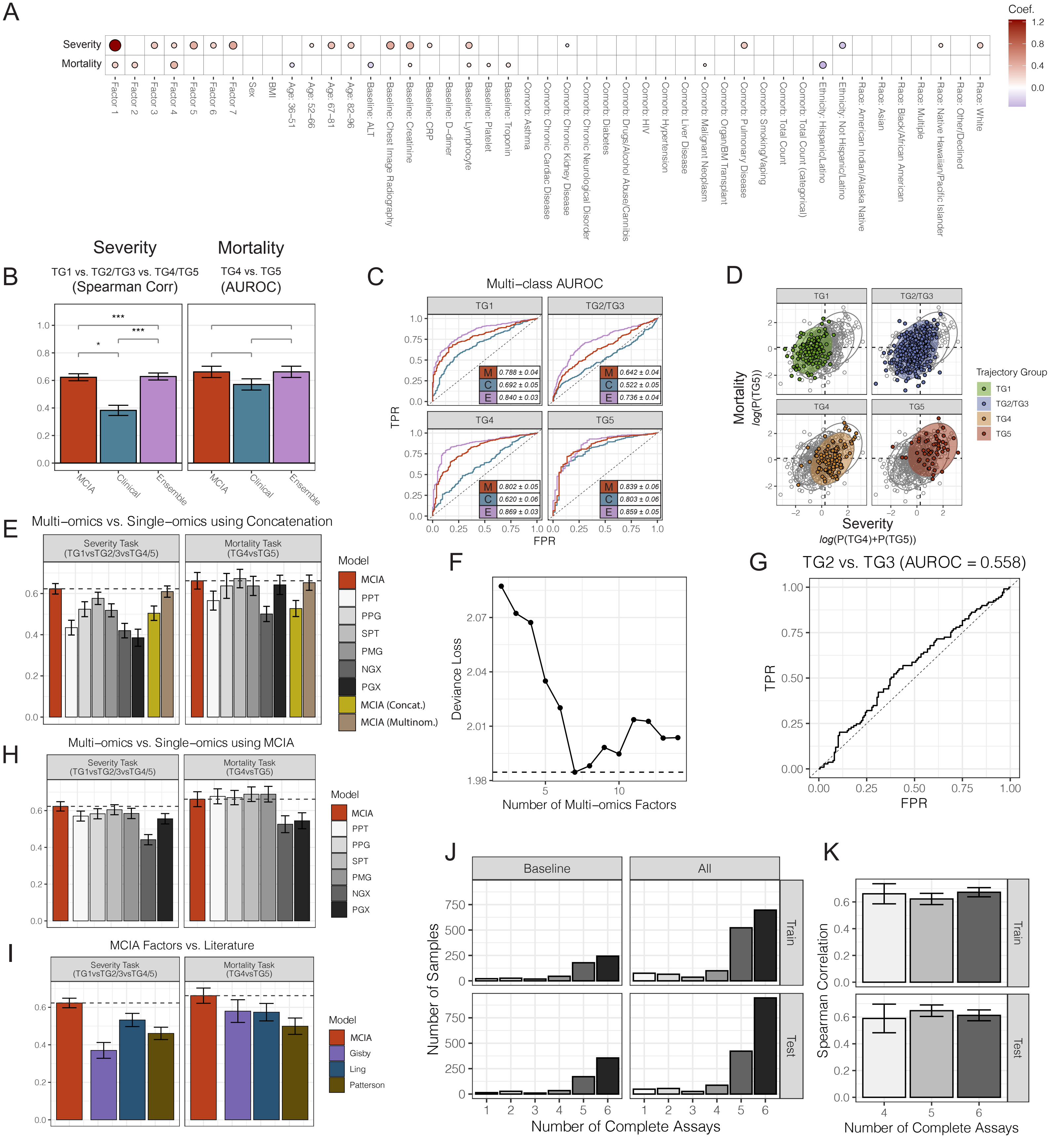

A
Severity
Mortality
Coef.
Factor 1
Factor 2
Factor 3
Factor 4
Factor 5
Factor 6
Factor 7
Sex
BMI
Age: 36–51
Age: 52–66
Age: 67–81
Age: 82–96
Baseline: ALT
Baseline: Chest Image Radiography
Baseline: Creatinine
Baseline: CRP
Baseline: D–dimer
Baseline: Lymphocyte
Baseline: Platelet
Baseline: Troponin
Comorb: Asthma
Comorb: Chronic Cardiac Disease
Comorb: Chronic Kidney Disease
Comorb: Chronic Neurological Disorder
Comorb: Diabetes
Comorb: Drugs/Alcohol Abuse/Cannibis
Comorb: HIV
Comorb: Hypertension
Comorb: Liver Disease
Comorb: Malignant Neoplasm
Comorb: Organ/BM Transplant
Comorb: Pulmonary Disease
Comorb: Smoking/Vaping
Comorb: Total Count
Comorb: Total Count (categorical)
Ethnicity: Hispanic/Latino
Ethnicity: Not Hispanic/Latino
Race: American Indian/Alaska Native
Race: Asian
Race: Black/African American
Race: Multiple
Race: Native Hawaiian/Pacific Islander
Race: Other/Declined
Race: White
B
Severity
TG1 vs. TG2/TG3 vs. TG4/TG5
(Spearman Corr)
Mortality
TG4 vs. TG5
(AUROC)
MCIA
Clinical
Ensemble
C
Multi–class AUROC
TG1
M 0.788 ± 0.04
C 0.692 ± 0.05
E 0.840 ± 0.03
TG2/TG3
M 0.642 ± 0.04
C 0.522 ± 0.05
E 0.736 ± 0.04
TG4
M 0.802 ± 0.05
C 0.620 ± 0.06
E 0.869 ± 0.03
TG5
M 0.839 ± 0.06
C 0.803 ± 0.06
E 0.859 ± 0.05
TPR
FPR
D
Mortality
log(P(TG5))
Severity
log(P(TG4)+P(TG5))
Trajectory Group
TG1
TG2/TG3
TG4
TG5
E
Multi–omics vs. Single–omics using Concatenation
Severity Task (TG1vsTG2/3vsTG4/5)
Mortality Task (TG4vsTG5)
Model
MCIA
PPT
PPG
SPT
PMG
NGX
PGX
MCIA (Concat.)
MCIA (Multinom.)
F
Deviance Loss
Number of Multi–omics Factors
G
TG2 vs. TG3 (AUROC = 0.558)
TPR
FPR
H
Multi–omics vs. Single–omics using MCIA
Severity Task (TG1vsTG2/3vsTG4/5)
Mortality Task (TG4vsTG5)
Model
MCIA
PPT
PPG
SPT
PMG
NGX
PGX
I
MCIA Factors vs. Literature
Severity Task (TG1vsTG2/3vsTG4/5)
Mortality Task (TG4vsTG5)
Model
MCIA
Gisby
Ling
Patterson
J
Baseline
All
Train
Test
Number of Samples
Number of Complete Assays
K
Spearman Correlation
Train
Test
Number of Complete Assays

**FIGURE S1: Additional computational results for prediction models, severity and mortality task, and MOFA imputation, related to Figure 2**. A) Dotplot of coefficients returned from the composite classifier, separated by MCIA factors and clinical features. B) Barplots of the severity and mortality tasks reporting Spearman correlation and AUROC values, respectively, on the training cohort. Results are shown for the MCIA, clinical, and ensemble models. C) Multi-class AUROC analysis predicting TG1, TG2/TG3, TG4, and TG5 separately for each model on the test cohort. D) Embedding of MCIA model predicted probabilities for both the mortality and severity tasks on the test cohort, faceted by each participant's TG. E) Severity and mortality task results on the training cohort for concatenated models. F) Connected scatter plot showing deviance loss of prediction models constructed using increasing numbers of multi-omics factors from MCIA. G) AUROC plot of MCIA model separating TG2 from TG3 on the training cohort. H, I) Severity and mortality task results on the training cohort for single-omic MCIA models and literature models. J) Histogram of participants grouped by the number of complete assays from PPT, PPG, SPT, PMG, NGX, and PGX, split by training/test (y-axis) and baseline/all visits (x-axis). K) Severity task results using only participants with 4, 5, and 6 complete assays on the training cohort.

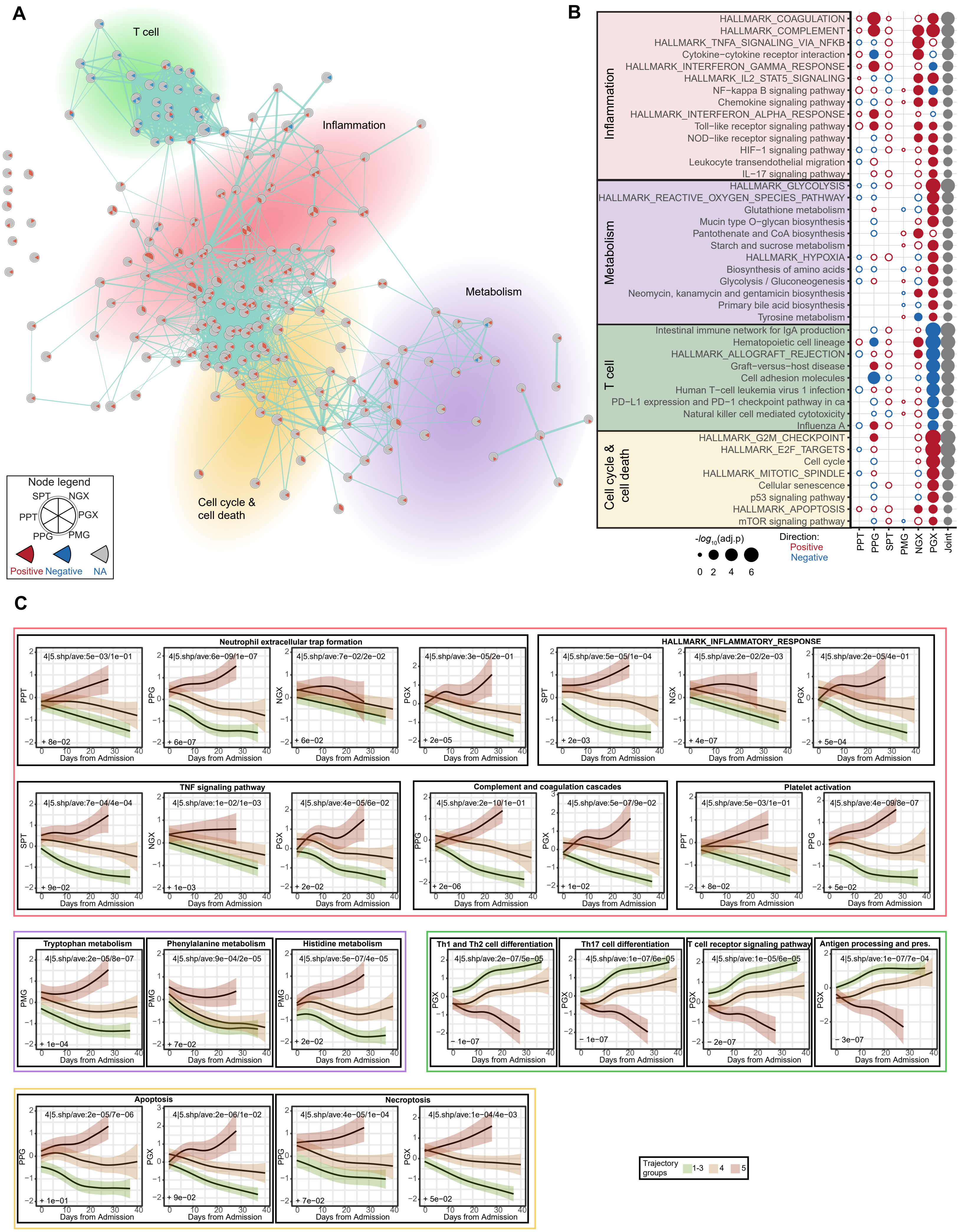
A
T cell
Inflammation
Metabolism
Cell cycle & cell death
Node legend
SPT NGX PPT PGX PPG PMG
Positive Negative NA
B
Inflammation
HALLMARK_COAGULATION
HALLMARK_COMPLEMENT
HALLMARK_TNFA_SIGNALING_VIA_NFKB
Cytokine−cytokine receptor interaction
HALLMARK_INTERFERON_GAMMA_RESPONSE
HALLMARK_IL2_STAT5_SIGNALING
NF−kappa B signaling pathway
Chemokine signaling pathway
HALLMARK_INTERFERON_ALPHA_RESPONSE
Toll−like receptor signaling pathway
NOD−like receptor signaling pathway
HIF−1 signaling pathway
Leukocyte transendothelial migration
IL−17 signaling pathway
Metabolism
HALLMARK_GLYCOLYSIS
HALLMARK_REACTIVE_OXYGEN_SPECIES_PATHWAY
Glutathione metabolism
Mucin type O−glycan biosynthesis
Pantothenate and CoA biosynthesis
Starch and sucrose metabolism
HALLMARK_HYPOXIA
Biosynthesis of amino acids
Glycolysis / Gluconeogenesis
Neomycin, kanamycin and gentamicin biosynthesis
Primary bile acid biosynthesis
Tyrosine metabolism
T cell
Intestinal immune network for IgA production
Hematopoietic cell lineage
HALLMARK_ALLOGRAFT_REJECTION
Graft−versus−host disease
Cell adhesion molecules
Human T−cell leukemia virus 1 infection
PD−L1 expression and PD−1 checkpoint pathway in ca
Natural killer cell mediated cytotoxicity
Influenza A
Cell cycle & cell death
HALLMARK_G2M_CHECKPOINT
HALLMARK_E2F_TARGETS
Cell cycle
HALLMARK_MITOTIC_SPINDLE
Cellular senescence
p53 signaling pathway
HALLMARK_APOPTOSIS
mTOR signaling pathway
PPT PPG SPT PMG NGX PGX Joint
-log10(adj.p)
0 2 4 6
Direction:
Positive
Negative
C
Neutrophil extracellular trap formation
HALLMARK_INFLAMMATORY_RESPONSE
TNF signaling pathway
Complement and coagulation cascades
Platelet activation
Tryptophan metabolism
Phenylalanine metabolism
Histidine metabolism
Th1 and Th2 cell differentiation
Th17 cell differentiation
T cell receptor signaling pathway
Antigen processing and pres.
Apoptosis
Necroptosis
Days from Admission
Trajectory groups
1-3
4
5

**FIGURE S2: Enrichment term clustering and selected pathway trajectories for the severity factor, related to Figure 3.** A) Visualization of functional enrichment using EnrichmentMap. The enriched terms are grouped based on biological functions as indicated using four colors and labels. The nodes correspond to enriched terms and each node is divided into six slices; each corresponds to one assay. The slices are colored based on the direction of enrichment: red if positive, blue if negative, and gray if the term is not tested or significantly enriched. The edge thickness corresponds to the number of shared leading-edge features between the terms. B) Pathway enrichment of severity factor for additional top enriched terms in each group in A apart from the ones in Fig 3C. The filled circles represent pathways with significant enrichment and the open circles without. Joint = aggregated p-value across omics. C) Trajectories for significant pathways in Fig 3C for different clinical trajectory groups. The p-values indicating whether the shape (shp) or average (avg) are significantly different between TG4 and TG5 are mentioned. TG1-TG3 are grouped together for better visualization.

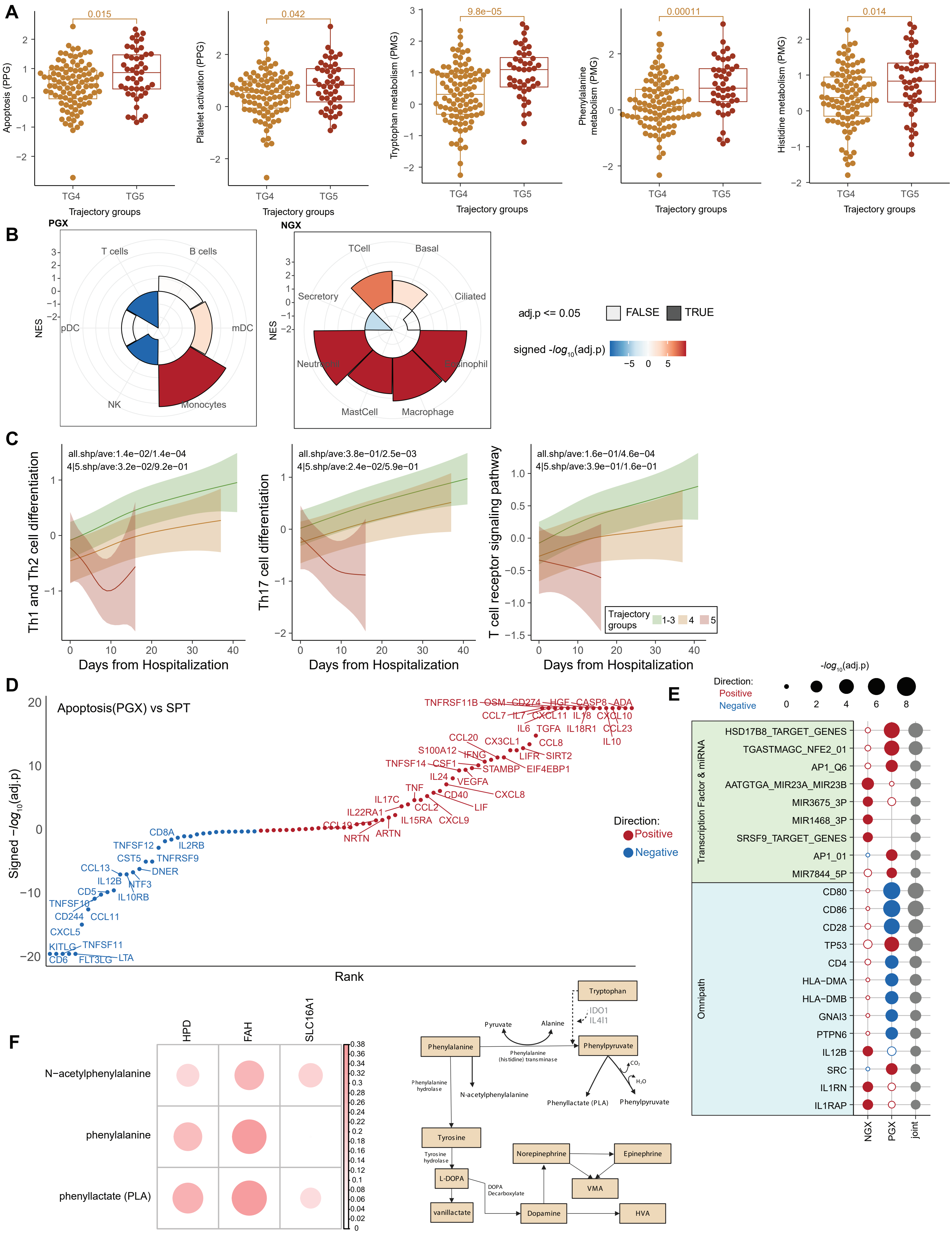
A
0.015
Apoptosis (PPG)
0.042
Platelet activation (PPG)
9.8e−05
Tryptophan metabolism (PMG)
0.00011
Phenylalanine metabolism (PMG)
0.014
Histidine metabolism (PMG)
TG4
TG5
Trajectory groups
B
PGX
T cells
B cells
pDC
mDC
NK
Monocytes
NES
NGX
TCell
Basal
Secretory
Ciliated
Neutrophil
Eosinophil
MastCell
Macrophage
adj.p <= 0.05
FALSE
TRUE
signed -log10(adj.p)
C
all.shp/ave:1.4e−02/1.4e−04
4|5.shp/ave:3.2e−02/9.2e−01
Th1 and Th2 cell differentiation
all.shp/ave:3.8e−01/2.5e−03
4|5.shp/ave:2.4e−02/5.9e−01
Th17 cell differentiation
all.shp/ave:1.6e−01/4.6e−04
4|5.shp/ave:3.9e−01/1.6e−01
T cell receptor signaling pathway
Trajectory groups
1-3
4
5
Days from Hospitalization
D
Apoptosis(PGX) vs SPT
Signed −log10(adj.p)
Rank
Direction:
Positive
Negative
TNFRSF11B OSM CD274 HGF CASP8 ADA
CCL7 IL7 CXCL11 IL18 CXCL10
IL6 TGFA IL18R1 CCL23
CCL20 CX3CL1 CCL8 IL10
S100A12 IFNG LIFR SIRT2
TNFSF14 CSF1 STAMBP EIF4EBP1
IL24 VEGFA
TNF CD40 CXCL8
IL17C CCL2 LIF
IL22RA1 IL15RA CXCL9
CCL19 NRTN ARTN
CD8A
TNFSF12 IL2RB
CST5 TNFRSF9
CCL13 DNER
IL12B NTF3
CD5 IL10RB
TNFSF10
CD244 CCL11
CXCL5
KITLG TNFSF11
CD6 FLT3LG LTA
E
Direction:
Positive
Negative
-log10(adj.p)
Transcription Factor & miRNA
HSD17B8_TARGET_GENES
TGASTMAGC_NFE2_01
AP1_Q6
AATGTGA_MIR23A_MIR23B
MIR3675_3P
MIR1468_3P
SRSF9_TARGET_GENES
AP1_01
MIR7844_5P
Omnipath
CD80
CD86
CD28
TP53
CD4
HLA-DMA
HLA-DMB
GNAI3
PTPN6
IL12B
SRC
IL1RN
IL1RAP
NGX
PGX
joint
F
HPD
FAH
SLC16A1
N−acetylphenylalanine
phenylalanine
phenyllactate (PLA)
Tryptophan
IDO1
IL4I1
Pyruvate
Alanine
Phenylalanine
Phenylpyruvate
Phenylalanine (histidine) transminase
Phenylalanine hydrolase
N-acetylphenylalanine
Phenyllactate (PLA)
Phenylpyruvate
CO2
H2O
Tyrosine
Tyrosine hydrolase
L-DOPA
DOPA Decarboxylate
vanillactate
Dopamine
Norepinephrine
Epinephrine
VMA
HVA

**FIGURE S3: Additional characterization of immune pathway associated with COVID-19 severity, related to Figures 3 and 4.** For the **severity factor**, A) Boxplots comparing levels of identified pathways (from Fig 3F) with moderate separating power between TG4 and TG5 at baseline (p.val < 0.05). B) Cell type enrichment of the PBMC (PGX) and nasal (NGX) transcriptomics based on their contribution to the severity factor. C) Trajectory plots depicting differences in PBMC transcriptomic signature of T cell pathways (sum of levels of all pathway features corrected for T cell frequencies) between trajectory groups. The Th1 and Th2 cell differentiation and Th17 cell differentiation pathways show significant decrease in TG5 compared to TG4 (4|5). T cell receptor signaling pathway is not significant, but the trend is preserved. D) Correlation between high-contribution cytokines and the apoptosis pathway activity constructed using PBMC transcriptomics (PGX). The cytokines are ranked based on the adj.p on X-axis and the cytokines with significant associations (adj.p < 0.05) are labeled. E) mHG Enrichment of the target genes of Transcription Factors and miRNAs (C3), and the downstream targets of receptor signaling (Omnipath) in the highly contributing PGX and NGX features of the severity factor. The filled circles represent pathways with significant enrichment and the open circles without. Joint = aggregated p-value across omics. F) The left panel displays the correlation between metabolites in the phenylalanine pathway and genes identified to play a role in actively converting phenylpyruvate from phenylalanine. The right panel features a simplified diagram of the tryptophan, phenylalanine, and tyrosine pathways. IDO1 can also degrade phenylalanine to phenylpyruvate, which can lead to a decrease in phenylalanine hydroxylase activity and an accumulation of phenylalanine in the body. The tyrosine metabolites (HVA, VMA, and VLA) are major terminal urinary metabolites that result from the conversion of L-Dopa, dopamine, and norepinephrine during catecholamine biosynthesis and degradation.

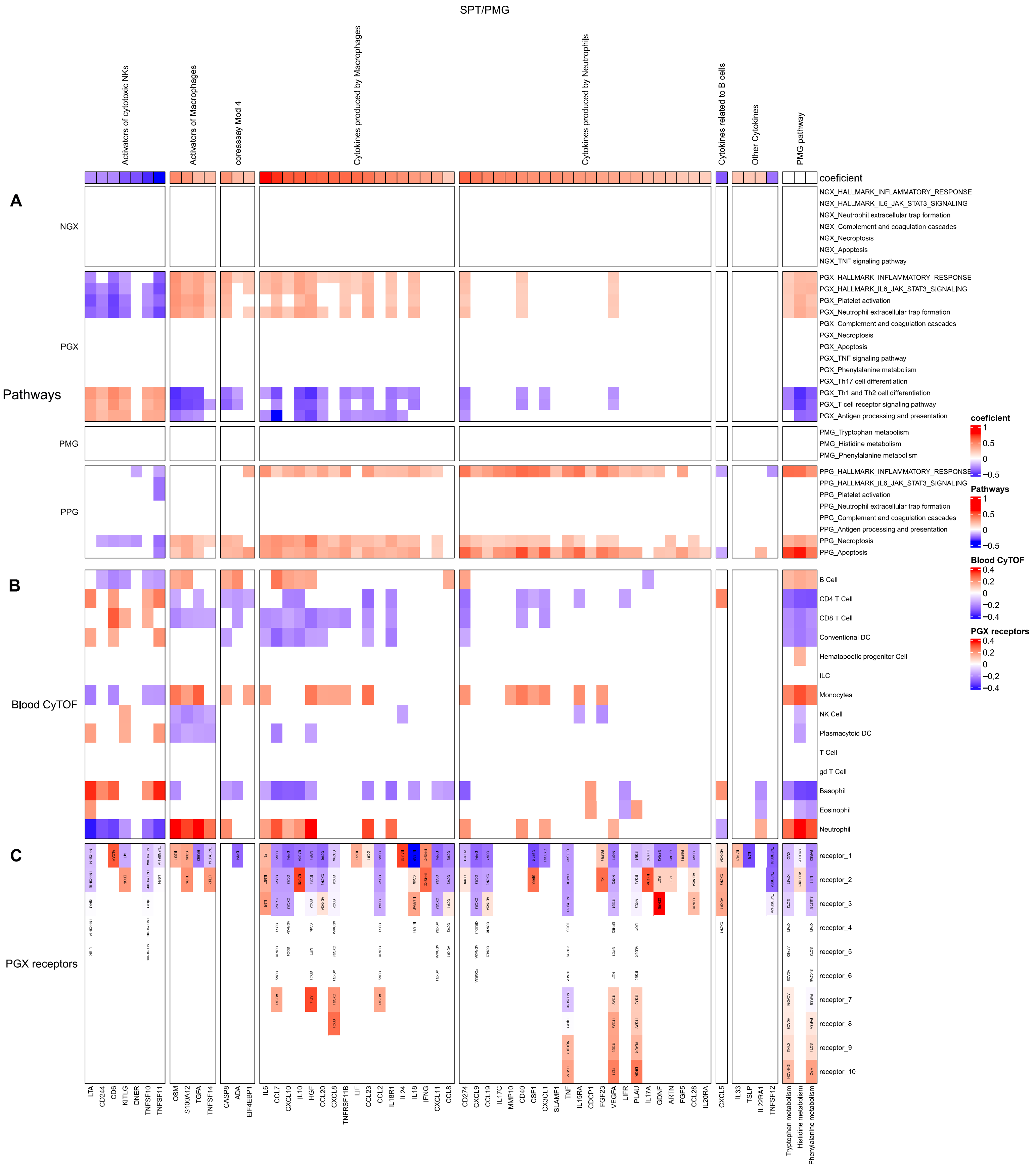

SPT/PMG
Activators of cytotoxic NKs
Activators of Macrophages
coreassay Mod 4
Cytokines produced by Macrophages
Cytokines produced by Neutrophils
Cytokines related to B cells
Other Cytokines
PMG pathway
coeficient
A
NGX
PGX
Pathways
PMG
PPG
B
Blood CyTOF
C
PGX receptors
NGX_HALLMARK_INFLAMMATORY_RESPONSE
NGX_HALLMARK_IL6_JAK_STAT3_SIGNALING
NGX_Neutrophil extracellular trap formation
NGX_Complement and coagulation cascades
NGX_Necroptosis
NGX_Apoptosis
NGX_TNF signaling pathway
PGX_HALLMARK_INFLAMMATORY_RESPONSE
PGX_HALLMARK_IL6_JAK_STAT3_SIGNALING
PGX_Platelet activation
PGX_Neutrophil extracellular trap formation
PGX_Complement and coagulation cascades
PGX_Necroptosis
PGX_Apoptosis
PGX_TNF signaling pathway
PGX_Phenylalanine metabolism
PGX_Th17 cell differentiation
PGX_Th1 and Th2 cell differentiation
PGX_T cell receptor signaling pathway
PGX_Antigen processing and presentation
PMG_Tryptophan metabolism
PMG_Histidine metabolism
PMG_Phenylalanine metabolism
PPG_HALLMARK_INFLAMMATORY_RESPONSE
PPG_HALLMARK_IL6_JAK_STAT3_SIGNALING
PPG_Platelet activation
PPG_Neutrophil extracellular trap formation
PPG_Complement and coagulation cascades
PPG_Antigen processing and presentation
PPG_Necroptosis
PPG_Apoptosis
B Cell
CD4 T Cell
CD8 T Cell
Conventional DC
Hematopoetic progenitor Cell
ILC
Monocytes
NK Cell
Plasmacytoid DC
T Cell
gd T Cell
Basophil
Eosinophil
Neutrophil
receptor_1
receptor_2
receptor_3
receptor_4
receptor_5
receptor_6
receptor_7
receptor_8
receptor_9
receptor_10
coeficient
1
0.5
0
−0.5
Pathways
1
0.5
0
−0.5
Blood CyTOF
0.4
0.2
0
−0.2
−0.4
PGX receptors
0.4
0.2
0
−0.2
−0.4
LTA
CD244
CD6
KITLG
DNER
TNFSF10
TNFSF11
OSM
S100A12
TGFA
TNFSF14
CASP8
ADA
EIF4EBP1
IL6
CCL7
CXCL10
IL10
HGF
CCL20
CXCL8
TNFRSF11B
LIF
CCL23
CCL2
IL18R1
IL24
IL18
IFNG
CXCL11
CCL8
CD274
CXCL9
CCL19
IL17C
MMP10
CD40
CSF1
CX3CL1
SLAMF1
TNF
IL15RA
CDCP1
FGF23
VEGFA
LIFR
PLAU
IL17A
GDNF
ARTN
FGF5
CCL28
IL20RA
CXCL5
IL33
TSLP
IL22RA1
TNFSF12
Tryptophan metabolism
Histidine metabolism
Phenylalanine metabolism

**FIGURE S4: Inter-omics analysis on top-contribution cytokines and significant pathways for the severity factor, related to Figures 3 and 4.** Serum soluble proteins were grouped into six WGCNA clusters previously defined in *Diray-Arce, et al.*[23], alongside the top three pathways identified in PMG. All correlation matrices were generated by controlling covariates such as sex, admission date, and enrollment site. A) Correlation of Serum soluble proteins and Tryptophan, Histidine, and Phenylalanine metabolism with the top pathways from PMG, PPG, NGX, and PGX in the severity Factor. NGX data did not show significant trans-omics correlations. Cytokines produced by neutrophils positively correlate with PMG and PPG pathways. Similarly, cytokines produced by macrophages and activators of Macrophages positively correlate with all other omics pathways. Additionally, we found a negative association between activators of cytotoxic NKs, which are negatively associated with severity factors, and other omics pathways. B) Correlation of parent cell type population, serum soluble proteins and Tryptophan, Histidine, and Phenylalanine metabolism. Monocytes and neutrophils showed positive correlation with PMG pathways. C) The reported receptors in CellTalkDb for the serum soluble proteins and their strength in the severity Factor. We showed the top 10 receptors per serum soluble proteins and metabolite pathway.

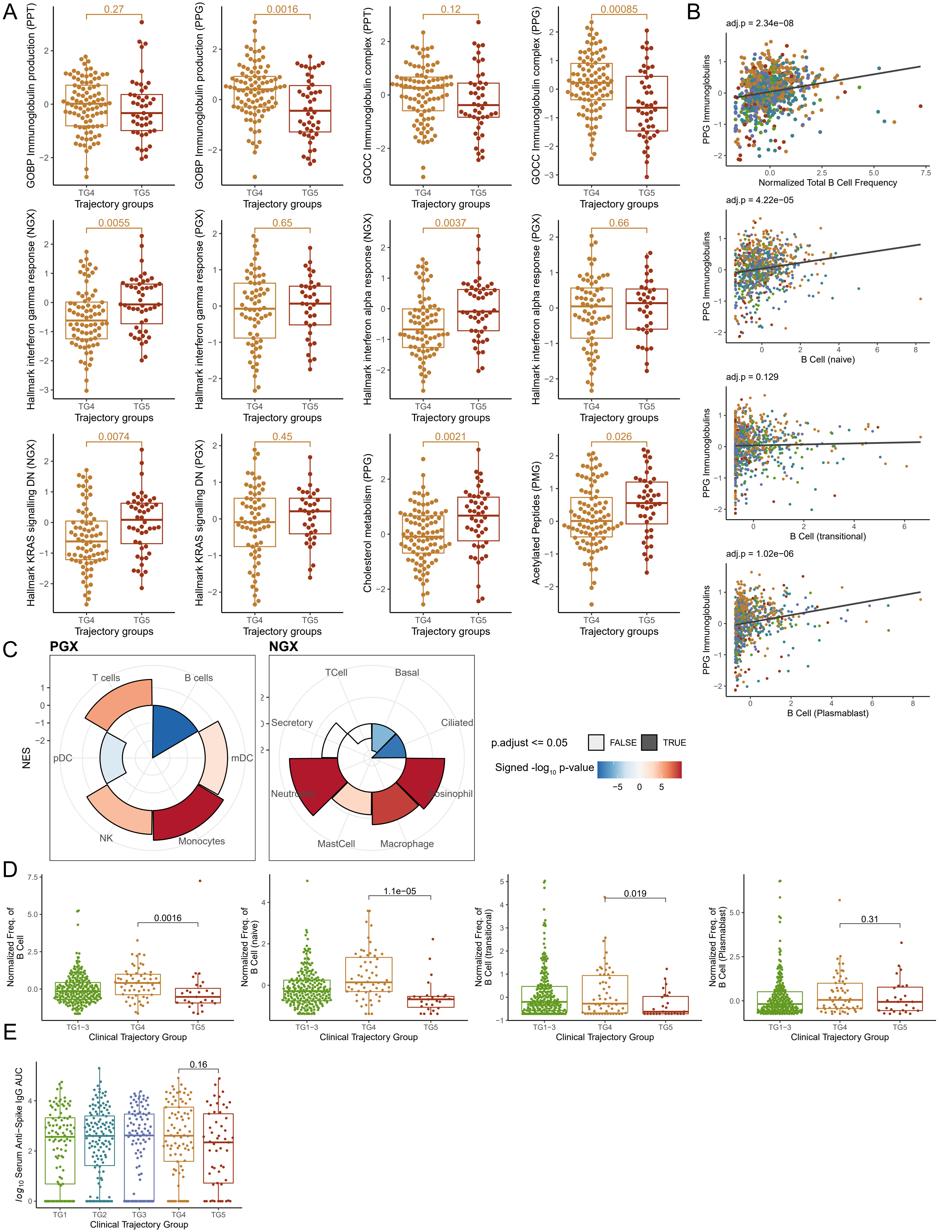
A
B
C
D
E
0.27
0.0016
0.12
0.00085
0.0055
0.65
0.0037
0.66
0.0074
0.45
0.0021
0.026
GOBP Immunoglobulin production (PPT)
GOBP Immunoglobulin production (PPG)
GOCC Immunoglobulin complex (PPT)
GOCC Immunoglobulin complex (PPG)
Hallmark interferon gamma response (NGX)
Hallmark interferon gamma response (PGX)
Hallmark interferon alpha response (NGX)
Hallmark interferon alpha response (PGX)
Hallmark KRAS signalling DN (NGX)
Hallmark KRAS signalling DN (PGX)
Cholesterol metabolism (PPG)
Acetylated Peptides (PMG)
TG4
TG5
Trajectory groups
adj.p = 2.34e−08
PPG Immunoglobulins
Normalized Total B Cell Frequency
adj.p = 4.22e−05
B Cell (naive)
adj.p = 0.129
B Cell (transitional)
adj.p = 1.02e−06
B Cell (Plasmablast)
PGX
NGX
NES
T cells
B cells
pDC
mDC
NK
Monocytes
TCell
Basal
Secretory
Ciliated
Neutrophil
Eosinophil
MastCell
Macrophage
p.adjust <= 0.05
FALSE
TRUE
Signed -log10 p-value
0.0016
1.1e−05
0.019
0.31
Normalized Freq. of B Cell
Normalized Freq. of B Cell (naive)
Normalized Freq. of B Cell (transitional)
Normalized Freq. of B Cell (Plasmablast)
TG1−3
Clinical Trajectory Group
0.16
log10 Serum Anti−Spike IgG AUC
TG1
TG2
TG3

**FIGURE S5: Additional characterization of immune pathway associated with COVID-19 mortality, related to Figure 5.** For the **mortality factor,** A) Boxplots comparing TG4 and TG5 at baseline across assays for enriched mortality factor pathway (adj.pval < 0.1) that separated by at least one measured omics (p.val < 0.05). B) Scatter plot between PPG (plasma proteomics global) immunoglobulin levels and total B cells/B cell subpopulation normalized frequencies. Significance was calculated using a linear mixed effect model (see Supplemental Methods, Inter-omics association analysis). C) Cell type enrichment of the PBMC and nasal transcriptomics based on their contribution to the mortality factor. D) Boxplots of total B cell normalized frequency and top associated B-cell subpopulations across TG groups at baseline. E) Boxplot of log10 Serum Anti-Spike IgG titers across TG groups at baseline.

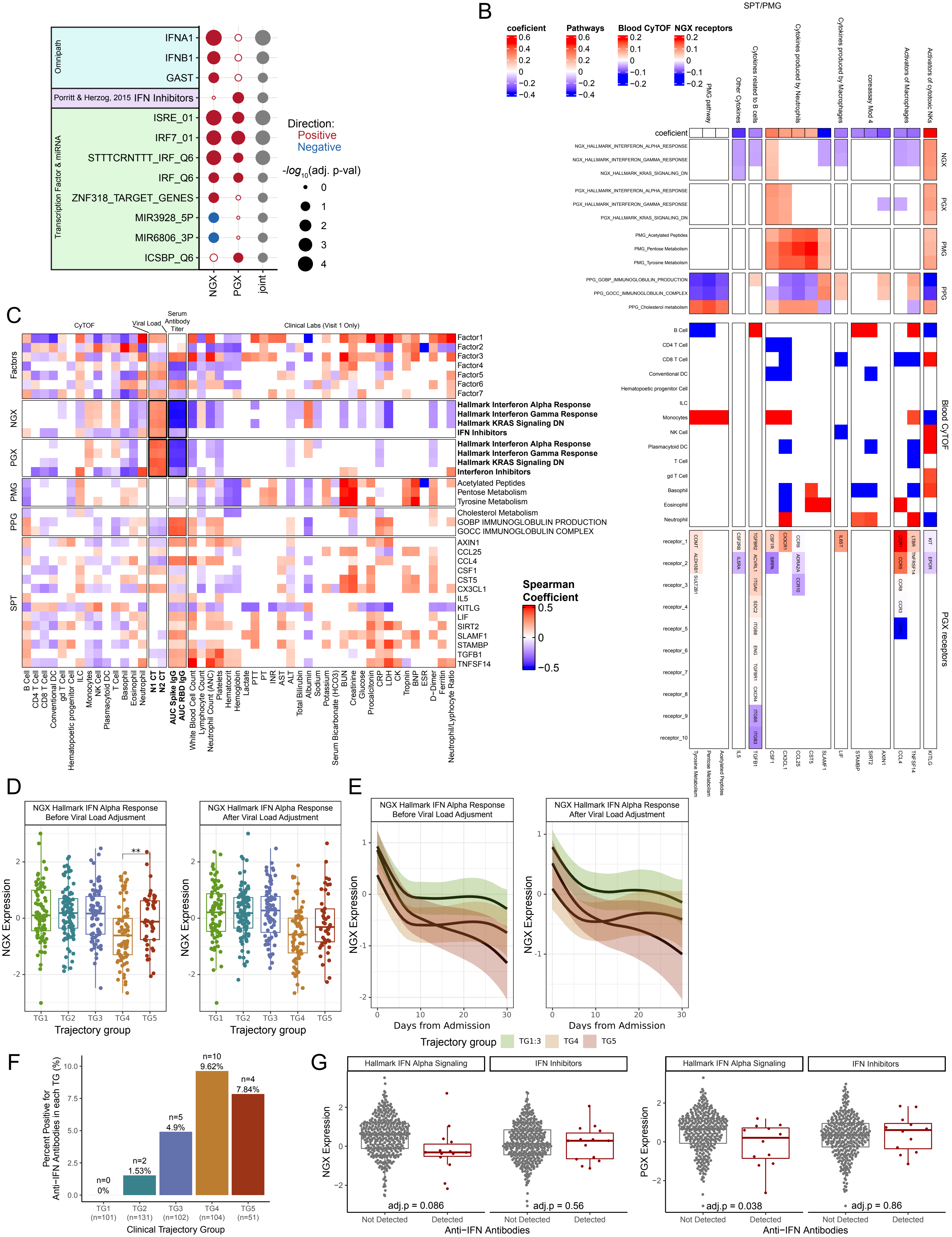

B
SPT/PMG
coeficient
Pathways
Blood CyTOF
NGX receptors
PMG pathway
Other Cytokines
Cytokines related to B cells
Cytokines produced by Neutrophils
Cytokines produced by Macrophages
coreassay Mod 4
Activators of Macrophages
Activators of cytotoxic NKs
NGX
PGX
PMG
PPG
Blood CyTOF
PGX receptors
Omnipath
IFNA1
IFNB1
GAST
Porritt & Herzog, 2015 IFN Inhibitors
Transcription Factor & miRNA
ISRE_01
IRF7_01
STTTCRNTTT_IRF_Q6
IRF_Q6
ZNF318_TARGET_GENES
MIR3928_5P
MIR6806_3P
ICSBP_Q6
NGX
PGX
joint
Direction:
Positive
Negative
-log10(adj. p-val)
C
CyTOF
Viral Load
Serum Antibody Titer
Clinical Labs (Visit 1 Only)
Factors
NGX
PGX
PMG
PPG
SPT
Factor1
Factor2
Factor3
Factor4
Factor5
Factor6
Factor7
Hallmark Interferon Alpha Response
Hallmark Interferon Gamma Response
Hallmark KRAS Signaling DN
IFN Inhibitors
Hallmark Interferon Alpha Response
Hallmark Interferon Gamma Response
Hallmark KRAS Signaling DN
Interferon Inhibitors
Acetylated Peptides
Pentose Metabolism
Tyrosine Metabolism
Cholesterol Metabolism
GOBP IMMUNOGLOBULIN PRODUCTION
GOCC IMMUNOGLOBULIN COMPLEX
AXIN1
CCL25
CCL4
CSF1
CST5
CX3CL1
IL5
KITLG
LIF
SIRT2
SLAMF1
STAMBP
TGFB1
TNFSF14
Spearman Coefficient
B Cell
CD4 T Cell
CD8 T Cell
Conventional DC
gd T Cell
Hematopoetic progenitor Cell
ILC
Monocytes
NK Cell
Plasmacytoid DC
T Cell
Basophil
Eosinophil
Neutrophil
N1 CT
N2 CT
AUC Spike IgG
AUC RBD IgG
White Blood Cell Count
Lymphocyte Count
Neutrophil Count (ANC)
Platelets
Hematocrit
Hemoglobin
Lactate
PTT
INR
AST
ALT
Total Bilirubin
Albumin
Sodium
Potassium
Serum Bicarbonate (HCO3)
BUN
Creatinine
Glucose
Procalcitonin
CRP
LDH
CK
Tropnin
BNP
ESR
D−Dimer
Ferritin
Neutrophil/Lymphocyte Ratio
D
NGX Hallmark IFN Alpha Response Before Viral Load Adjusment
NGX Hallmark IFN Alpha Response After Viral Load Adjusment
NGX Expression
Trajectory group
TG1
TG2
TG3
TG4
TG5
E
NGX Hallmark IFN Alpha Response Before Viral Load Adjusment
NGX Hallmark IFN Alpha Response After Viral Load Adjusment
NGX Expression
Days from Admission
Trajectory group
TG1:3
TG4
TG5
F
Percent Positive for Anti-IFN Antibodies in each TG (%)
n=0
0%
n=2
1.53%
n=5
4.9%
n=10
9.62%
n=4
7.84%
TG1 (n=101)
TG2 (n=131)
TG3 (n=102)
TG4 (n=104)
TG5 (n=51)
Clinical Trajectory Group
G
Hallmark IFN Alpha Signaling
IFN Inhibitors
NGX Expression
PGX Expression
adj.p = 0.086
adj.p = 0.56
adj.p = 0.038
adj.p = 0.86
Not Detected
Detected
Anti−IFN Antibodies

**FIGURE S6: Additional characterization of interferon signaling, anti-IFN auto-antibodies, and inter-omics analysis of top-contribution cytokines and significant pathways for the severity factor, related to Figures 5 and 6.** A) mHG Enrichment of Transcription Factors and miRNA (C3), IFN inhibitors, and downstream receptor signaling (Omnipath) for PBMC and Nasal Transcriptomics associated with the Mortality Factor. Joint = aggregated p-value across omics. B) Top: Spearman Correlation of top metabolic pathways and SPT soluble proteins against significant mortality associated pathways in the PMG, PPG, NGX, and PGX. Middle: Spearman Correlation of top metabolic pathways and SPT soluble proteins against whole blood CyTOF parent population frequencies. Bottom: Spearman Correlation of top metabolic pathways and SPT soluble proteins against their receptors' expression in the PGX (CelltalkDB for SPT and RaMP for PMG). C) Spearman Correlation of Trans-omic Severity and Mortality associated biological pathways/features with additional omics including whole blood CyTOF cell type frequency, nasal viral load (inverted RT-qPCR CT), Anti-Spike and Anti-RPB Antibody Titers, and Baseline Clinical laboratory tests. Non-significant correlations are white. D) Nasal and PBMC Hallmark Interferon alpha response at visit 1 between clinical trajectory groups unadjusted and adjusted for nasal viral load. E) Nasal Hallmark Interferon alpha response over 30 days unadjusted and adjusted for nasal viral load. F) Presence of Anti-IFN antibodies per clinical trajectory group. G) Hallmark IFN alpha response and expression of IFN inhibitors in Nasal and PBMC transcriptomics at visit 1 for participants with and without detectable anti-IFN antibodies.

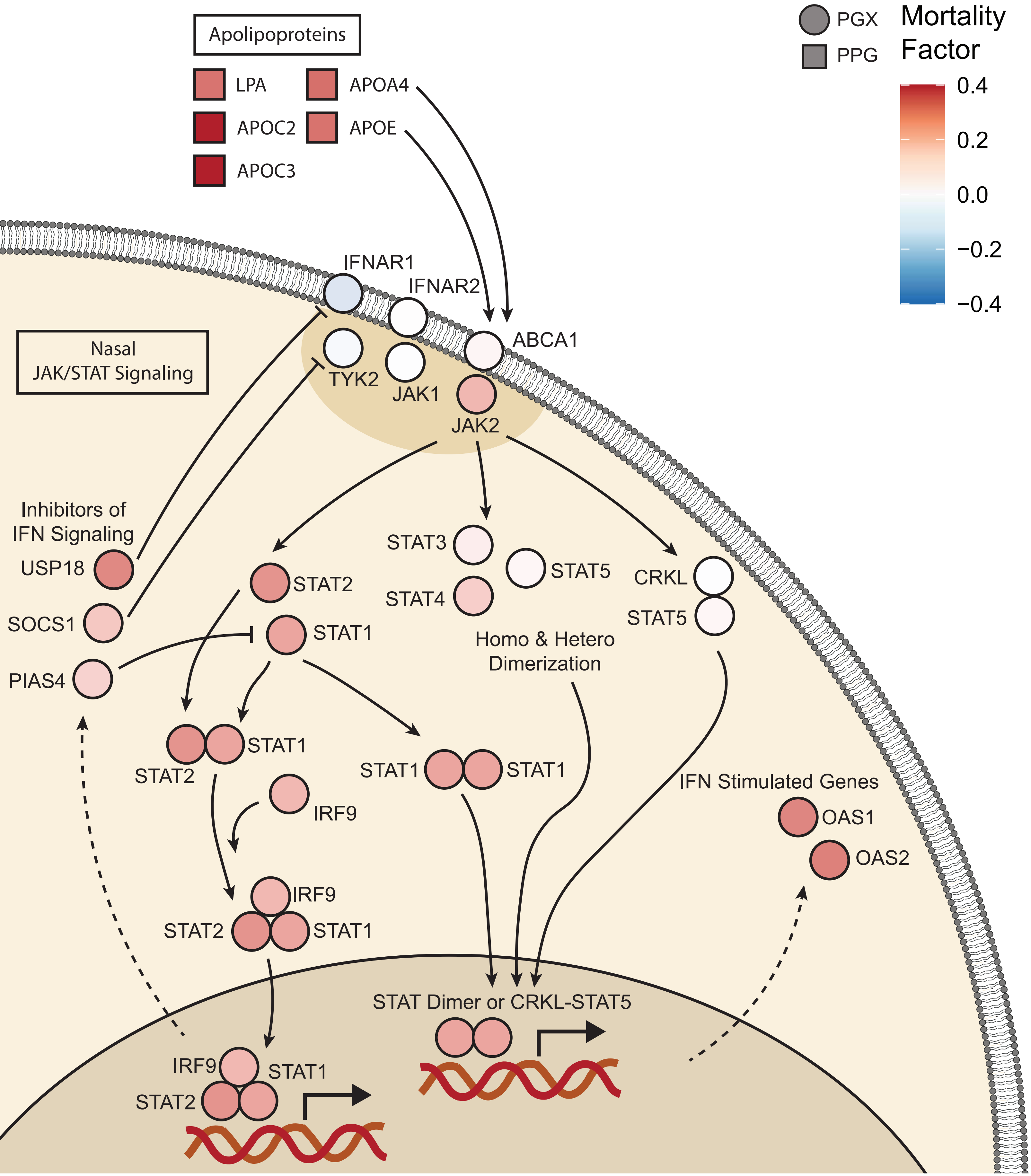

PGX
PPG
Mortality Factor
0.4
0.2
0.0
−0.2
−0.4
Apolipoproteins
LPA
APOA4
APOC2
APOE
APOC3
IFNAR1
IFNAR2
ABCA1
Nasal JAK/STAT Signaling
TYK2
JAK1
JAK2
Inhibitors of IFN Signaling
USP18
SOCS1
PIAS4
STAT3
STAT5
STAT4
STAT2
STAT1
CRKL
STAT5
Homo & Hetero Dimerization
STAT1
STAT2
STAT1
STAT1
IRF9
IFN Stimulated Genes
OAS1
OAS2
IRF9
STAT2
STAT1
STAT Dimer or CRKL-STAT5
IRF9
STAT1
STAT2

**FIGURE S7: Virus-centered integrative multi-omics network of the mortality(-associated) factor (Factor 4) in PGX, related to Figure 6.** IFN signaling is enriched in both the NGX (Fig 6) and PGX (Fig S7). Key IFN signaling is associated with the mortality factor in the PGX including STAT gene expression, ISGs, and downstream upregulation of IFN inhibitors which can generate negative feedback and inhibit upstream signaling.

**Key resources table**

| REAGENT or RESOURCE | SOURCE | IDENTIFIER |
|---|---|---|
| Antibodies | | |
| Maxpar® Direct™ Immune Profiling Assay (MDIPA) Kit | Fluidigm | Cat#201325 |
| CD8a-146Nd | Fluidigm | Cat#3146001B; RRID:AB_2687641 |
| Granzyme B Antibody, anti-human/mouse/rat, REAfinity | Miltenyi | Cat#130-116-486 |
| Goat Anti-Human IgA-UNLB | Southern Biotech | Cat#2050-01 |
| Purified anti-human IgM Antibody | Biolegend | Cat#314502 |
| Mouse Anti-Human IgG1 Fc-UNLB | Souther Biotech | Cat#9054-01 |
| Purified anti-mouse/human CD11b Antibody | Biolegend | Cat#101202 |
| Purified anti-human/mouse/rat CD278 (ICOS) Antibody | Biolegend | Cat#313502 |
| Purified anti-human CD39 Antibody | Biolegend | Cat#328202 |
| Purified anti-human CD169 (Sialoadhesin, Siglec-1) Antibody | Biolegend | Cat#346002 |
| Purified anti-human CD64 (Maxpar® Ready) Antibody | Biolegend | Cat#305029 |
| Purified anti-human CD71 Antibody | Biolegend | Cat#334102 |
| Anti-Human CD279/PD-1 (EH12.2H7)-175Lu | Fluidigm | Cat#3175008B |
| Anti-Human CD61 (VI-PL2)-209Bi | Fluidigm | Cat#3209001B |
| Anti-Human CD3 (UCHT1)-141Pr antibody | Fluidigm | Cat#3141019B |
| Anti-Human HLA-DR (L243)-143Nd antibody | Fluidigm | Cat#3143013B |
| Anti-Human CD69 (FN50)-144Nd antibody | Fluidigm | Cat#3144018B |
| Anti-Human CD4 (RPA-T4)-145Nd antibody | Fluidigm | Cat#3145001B |
| Anti-Human CD8a (RPA-T8)-146Nd antibody | Fluidigm | Cat#3146001B |
| Anti-Human CD20 (2H7)-147Sm antibody | Fluidigm | Cat#3147001B |
| Anti-Human CD127 (A019D5)-149Sm antibody | Fluidigm | Cat#3149011B |
| Anti-Human MIP-1β (D21-1351)-150Nd antibody | Fluidigm | Cat#3150004B |
| Anti-Human CD123 (6H6)-151Eu antibody | Fluidigm | Cat#3151001B |
| Anti-Human TNFα (Mab11)-152Sm antibody | Fluidigm | Cat#3152002B |
| Anti-Human CD62L (DREG-56)-153Eu antibody | Fluidigm | Cat#3153004B |
| Anti-Human CD45 (HI30)-154Sm antibody | Fluidigm | Cat#3154001B |
| Anti-Human IL-6 (MQ2-13A5)-156Gd antibody | Fluidigm | Cat#3156011B |
| Anti-Human IFN-γ (B27)-158Gd antibody | Fluidigm | Cat#3158017B |
| Anti-Human CD11c (Bu15)-159Tb antibody | Fluidigm | Cat#3159001B |
| Anti-Human CD14 (M5E2)-160Gd antibody | Fluidigm | Cat#3160001B |
| Anti-Human CD80/B7.1 (2D10.4)-161Dy antibody | Fluidigm | Cat#3161023B |
| Anti-Human CD66b (80H3)-162Dy antibody | Fluidigm | Cat#3162023B |
| Anti-Human CD56 (NCAM16.2)-163Dy antibody | Fluidigm | Cat#3163007B |
| Anti-Human CD15 (W6D3)-164Dy antibody | Fluidigm | Cat#3164001B |
| Anti-Human CD61 (VI-PL2)-165Ho antibody | Fluidigm | Cat#3165010B |
| Anti-Human CD11b (ICRF44)-167Er antibody | Fluidigm | Cat#3167011B |
| Anti-Human CD206 (15-2)-168Er antibody | Fluidigm | Cat#3168008B |
| Anti-Human CD54 (HA58)-170Er antibody | Fluidigm | Cat#3170014B |
| Anti-Human CD68 (Y1/82A)-171Yb antibody | Fluidigm | Cat#3171011B |
| Anti-Human CD16 (3G8)-209Bi antibody | Fluidigm | Cat#3209002B |
| Anti- CoV Nucleocapsid protein (6H3) antibody | Abcam | Cat#ab273434 |
| Anti-Human Eotaxin (43915) antibody | R&D | Cat#MAB3201 |
| Anti-Human ACE-2 (535919) antibody | NOVUS | Cat#MAB9332-100 |

| | | |
|---|---|---|
| Anti-Human Cytokeratin (C-11) antibody | Biolegend | Cat#628602 |
| Anti- CoV Spike protein (1A9) antibody | GeneTex | Cat#GTX632604 |
| Anti-Human EPX (MM82.2.1) antibody | MAYO CLINIC | https://www.mayoclinic.org |
| Anti-Human IL-8 (E8N1) antibody | Biolegend | Cat#511402 |
| Anti-Human IL-1β (H1b-27) antibody | Biolegend | Cat#511602 |
| Anti-Human IFN-β (IFNb/A1 ) antibody | Biolegend | Cat#514002 |
| Anti-Human Siglec-8 (837535) antibody | R&D | Cat#MAB7975 |
| Anti-human IgG (Fc specific)-Peroxidase antibody produced in goat | Sigma-Aldrich | Cat#A0170; RRID: AB_257868 |
| Goat anti-human IgM-HRP | SouthernBiotech | Cat#2020-05; RRID: AB_2795603 |
| Anti-human IgA (α-chain specific)-Peroxidase antibody produced in goat | Sigma-Aldrich | Cat#A0295; RRID: AB_257876 |
| Anti-Glial Fibrillary Associated Protein | Agilent | Cat#Z033429-2 |
| Anti-human IgG (PE) | ThermoScientific | Cat#12-4998-8 |
| Anti-human pSTAT1 (AF647) | BD | Cat#612597 |
| Anti-human CD14 (FITC) | BD | Cat#555397 |
| Bacterial and virus strains | | |
| BLT5403, T7 Select Kit | Novagen | Cat#70550-3 |
| T7 Bacteriophage, T7 Select Kit | Novagen | Cat#70550-3 |
| Biological samples | | |
| Plasma samples from IMPACC cohort | Multiple clinical sites | N/A |
| Whole blood from hospitalized COVID19 patients- collected in EDTA tubes | Multiple clinical sites | N/A |
| Veri-Cells™ Heavy Metal (Ta) PBMC | Biolegend | Cat#427203 |
| Serum samples from IMPACC cohort | Multiple clinical sites | N/A |
| Stimulated Plasma from Healthy Controls | Stanford University | N/A |
| Plasma from Healthy Controls | Stanford University | N/A |
| Serum from Healthy Controls | Stanford University | N/A |
| Chemicals, peptides, and recombinant proteins | | |
| DNA/RNA Shield Collection Tube w/ Swab - DX | Zymo Research | Cat#R1107-E |
| Quick-DNA/RNA MagBead | Zymo Research | Cat#R2131 |
| Stranded Total RNA Prep, Ligation with Ribo-Zero Plus | Illumina | Cat#20040529 |
| HS NGS Fragment Kit | Agilent | Cat#DNF-474-0500 |
| K-562 Total RNA | Thermo Fisher | Cat#AM7832 |
| qScript XLT 1-Step RT-qPCR ToughMix | Quantabio | Cat#95133-02K |
| 2-propanolol (LC-MS) | MilliporeSigma | Cat#1027814000 |
| Acetonitrile (LC-MS) | MilliporeSigma | Cat# 1000294000 |
| Water, Baker Analyzed LC/MS Reagent Grade | J.T. Baker | Cat#9831-02 |
| Ammonium Formate (LC-MS) | J.T. Baker | Cat#M530-08 |
| Perfluoropentanoic acid | Sigma | Cat#396575 |
| Ammonium Bicarbonate | Fisher | Cat#A643 |
| Ammonium Hydroxide | Sigma | Cat#338818 |
| Cell-ID™ 20-Plex Pd Barcoding Kit | Fluidigm | Cat#201060 |
| Saponin | Sigma | Cat#47036 |
| Human TruStain FcX™ (Fc Receptor Blocking Solution) | Biolegend | Cat#422302; RRID:AB_2818986 |
| Heparin sodium salt | Sigma | Cat#H3393 |

| | | |
|---|---|---|
| SmartTube PROT1 stabilizer PROT1-250ML | SmartTube | Fisher Cat# 501351692 |
| SmartTube ThawLyse - THAWLYSE1 | SmartTube | Fisher Cat# 501351696 |
| Paraformaldehyde (PFA), 16% w/v aqueous, methanol-free | Alfa Aesar | Fisher Cat# AA433689L |
| Fetal bovine serum, characterized, heat-inactivated | HyClone | Fisher Cat#SH30396.03 |
| Dimethyl sulfoxide | Fisher | Cat#BP231-100 |
| Maxpar MCP9 Antibody Labeling Kit, 111Cd | Fluidigm | Cat#201111A |
| Maxpar MCP9 Antibody Labeling Kit, 112Cd | Fluidigm | Cat#201112A |
| Maxpar MCP9 Antibody Labeling Kit, 114Cd | Fluidigm | Cat#201114A |
| Maxpar MCP9 Antibody Labeling Kit, 116Cd | Fluidigm | Cat#201116A |
| Maxpar® X8 Antibody Labeling Kit, 142Nd | Fluidigm | Cat#201142B |
| Maxpar® X8 Antibody Labeling Kit, 159Tb | Fluidigm | Cat#201159B |
| Maxpar® X8 Antibody Labeling Kit, 162Dy | Fluidigm | Cat#201162B |
| Maxpar® X8 Antibody Labeling Kit, 165Ho | Fluidigm | Cat#201165B |
| Maxpar® X8 Antibody Labeling Kit, 169Tm | Fluidigm | Cat#201169B |
| Maxpar® X8 Antibody Labeling Kit, 142Nd—4 Rxn | Fluidigm | Cat#201142A |
| Maxpar® X8 Antibody Labeling Kit, 148Nd—4 Rxn | Fluidigm | Cat#201148A |
| Maxpar® X8 Antibody Labeling Kit, 155Gd—4 Rxn | Fluidigm | Cat#201155A |
| Maxpar® X8 Antibody Labeling Kit, 166Er—4 Rxn | Fluidigm | Cat#201166A |
| Maxpar® X8 Antibody Labeling Kit, 169Tm—4 Rxn | Fluidigm | Cat#201169A |
| Maxpar® X8 Antibody Labeling Kit, 172Er—4 Rxn | Fluidigm | Cat#201172A |
| Maxpar® X8 Antibody Labeling Kit, 173Yb—4 Rxn | Fluidigm | Cat#201173A |
| Maxpar® X8 Antibody Labeling Kit, 174Yb—4 Rxn | Fluidigm | Cat#201174A |
| Maxpar® X8 Antibody Labeling Kit, 175Lu—4 Rxn | Fluidigm | Cat#201175A |
| Maxpar® X8 Antibody Labeling Kit, 176Yb—4 Rxn | Fluidigm | Cat#201176A |
| Cell-ID™ Cisplatin | Fluidigm | Cat#201064 |
| Cell-ID™ Intercalator | Fluidigm | Cat#201192A |
| Cell-ID™ 20-Plex Pd Barcoding Kit | Fluidigm | Cat#201060 |
| Maxpar® Water—500 mL | Fluidigm | Cat#201069 |
| Maxpar® Cell Staining Buffer | Fluidigm | Cat#201068 |
| Maxpar® PBS | Fluidigm | Cat#201058 |
| EQ Four Element Calibration Beads | Fluidigm | Cat#201078 |
| Bond-Breaker TCEP Solution, Neutral pH | Thermo Fisher | Cat#77720 |
| PFA | EMC | 50-980-487 |
| Osmium tetroxide | ACROS ORGANICS | 319010050 |
| Recombinant SARS-CoV-2 receptor binding domain (RBD) | Krammer Laboratory at the Icahn School of Medicine at Mount Sinai | https://labs.icahn.mssm.edu/krammerlab/reagents/ |
| Recombinant SARS-CoV-2 spike protein (S) | Krammer Laboratory at the Icahn School of Medicine at Mount Sinai | https://labs.icahn.mssm.edu/krammerlab/reagents/ |
| SIGMAFAST™ OPD (o-Phenylenediamine dihydrochloride) | Sigma-Aldrich | Cat#P9187 |
| 3-molar hydrochloric acid | Thermo Fisher Scientific | Cat#S25856 |
| Tween-20 | Fisher Bioreagents | Cat#BP337-100 |

| | | |
|---|---|---|
| Non-fat dry milk Omniblok | AmericanBio | Cat#AB10109-01000 |
| Bovine Serum Albumin Fraction V | Roche | Cat#10735078001 |
| Protein A conjugated magnetic beads | Invitrogen | Cat#10008D |
| Protein G conjugated magnetic beads | Invitrogen | Cat#10009D |
| T4 ligase | New England Biolabs | Cat#M0202S |
| Phusion DNA Polymerase | New England Biolabs | Cat# M0530L |
| Urea | Sigma-Aldrich | |
| Ammonium Bicarbonate | Sigma-Aldrich | 09830-1KG |
| Iodoacetamide | Sigma-Aldrich | I1149-25G |
| Dithiothreitol | Sigma-Aldrich | D9779-10G |
| LC/MS grade Formic Acid | Thermo Scientific | A117-50 |
| Perchloric Acid | Sigma-Aldrich | 311421-50ML |
| 1-Propanol | Sigma-Aldrich | 34871-1L |
| Sera-Mag Speed Beads 65 | Sigma-Aldrich | 65152105050250 |
| Sera-Mag Speed Beads 45 | Sigma-Aldrich | 45152105050250 |
| HPLC grade Water | Fisher chemical | W5-4 |
| LC/MS grade Water | Fisher chemical | W6-1 |
| LC/MS grade Acetonitrile | Fisher chemical | A955-1 |
| HPLC grade Methanol | Fisher chemical | A452-4 |
| LC/MS grade Methanol | Fisher chemical | A456-4 |
| LC/MS grade Isopropanol | Fisher chemical | A461-1 |
| Sequence grade Porcine Trypsin | Promega | V5117 |
| K562 Cell Line Tryptic Peptide Mixture Standard 100 µg | Promega | V6951 |
| Trifluoroacetic acid | Sigma-Aldrich | T6508-100ML |
| Ambion Nuclease-Free Water | Invitrogen | Cat#AM9937 |
| Recombinant human IFNa | R&D | Cat#11101-2 |
| Recombinant human IFNb | Peprotech | Cat#300-02BC |
| Recombinant human IFNw | Peprotech | Cat#300-02J |
| Sulfo-NHS | ThermoScientific | Cat#A39269 |
| EDC | ThermoScientific | Cat#77149 |
| Critical commercial assays | | |
| Quick-DNA/RNA Pathogen MagBead | Zymo Research | R2146 |
| RNase-Free DNase Set | Qiagen | 79254 |
| NEBNext Ultra II Directional RNA Library Prep Kit for Illumina | New England Biolabs | E7760 |
| AMPure XP Beads | Beckman-Coulter | A63882 |
| Quick-RNA MagBead Kit | Zymo Research | R2133 |
| SMART-Seq v4 Ultra Low Input RNA Kit for Sequencing | Takara Bio | 634894 |
| Nextera XT DNA Library Preparation Kit | Illumina | FC-131-1096 |
| DNA Prep, Tagmentation | Illumina | 20018705 |
| Chemagic Blood 400 (96) kit | Perkin Elmer | CMG-1091 |
| Global Diversity Array (GDA) | Illumina | 20031810 |
| Covaris E210 | Covaris, LLC. | 10521 |
| T7 Select 10-3b Cloning kit | EMD Millipore | EMD Millipore |
| AMPure XP Beads | Beckman Coulter | Cat#A63881 |
| Olink Target 96 Inflammation Reagent Kit | Olink Proteomics | Cat#95302, Lot#B02101 |
| Deposited data | | |
| Experimental models: Cell lines | | |

| | | |
|---|---|---|
| Expi293F cells | Thermo Fisher | Cat#A14528 |
| Experimental models: Organisms/strains | | |
| Oligonucleotides | | |
| 2019-nCOV_N1-F GAC CCC AAA ATC AGC GAA AT | Integrated DNA technologies | Cat#10006713 |
| 2019-nCOV_N1-R TCT GGT TAC TGC CAG TTG AAT CTG | Integrated DNA technologies | Cat#10006713 |
| 2019-nCOV_N1-P ACC CCG CAT TAC GTT TGG TGG ACC | Integrated DNA technologies | Cat#10006713 |
| 2019-nCOV_N2-F TTA CAA ACA TTG GCC GCA AA | Integrated DNA technologies | Cat#10006713 |
| 2019-nCOV_N2-R GCG CGA CAT TCC GAA GAA | Integrated DNA technologies | Cat#10006713 |
| 2019-nCOV_N2-P ACA ATT TGC CCC CAG CGC TTC AG | Integrated DNA technologies | Cat#10006713 |
| RP-F AGA TTT GGA CCT GCG AGC G | Integrated DNA technologies | Cat#10006713 |
| RP-R GAG CGG CTG TCT CCA CAA GT | Integrated DNA technologies | Cat#10006713 |
| RP-P TTC TGA CCT GAA GGC TCT GCG CG | Integrated DNA technologies | Cat#10006713 |
| SARS-CoV-2 tilling oligonucleotides for whole genome amplification | Gonzalez-Reiche, et al. 2020 | https://doi.org/10.1126/science.abc1917 |
| Recombinant DNA | | |
| Vector pCAGGS Containing the SARS-Related Coronavirus 2, Wuhan-Hu-1 Spike Glycoprotein Gene (soluble, stabilized) | BEI Resources | Cat#NR-52394 |
| Vector pCAGGS Containing the SARS-Related Coronavirus 2, Wuhan-Hu-1 Spike Glycoprotein Receptor Binding Domain (RBD) | BEI Resources | Cat#NR-52309 |
| Human Coronavirus Synthetic DNA | Twist Bioscience | https://www.twistbioscience.com |
| Software and algorithms | | |
| CZID Pipeline | Chan Zuckerberg Initiative | www.czid.org |
| bcl2fastq v2.20.0.422 | Illumina | https://support.illumina.com/sequencing/sequencing_software/bcl2fastq-conversion-software.html |
| FastQC_v0.11.5 | Andrew S | N/A |
| STARv2.4.3a | Dobin et al, 2013 | https://github.com/alexdobin/STAR |
| Qualimap | Okonechnikov et al, 2015 | http://qualimap.conesalab.org |
| Cutadapt_v3.7 | DOI:10.14806/ej.17.1.200 | https://cutadapt.readthedocs.io/en/stable/ |
| Preseq_v3.1.1 | Timothy D and Andrew Smith et al, 2013 | https://github.com/smithlabcode/preseq |
| Samtools_v1.12 | Heng Li et al, 2009 | http://samtools.sourceforge.net |

| | | |
|---|---|---|
| MultiQC | Philip Ewels | https://multiqc.info |
| WGCNA R package (version 1.69-81 ) | Langfelder, Peter, and Steve Horvath. "WGCNA: an R package for weighted correlation network analysis." BMC bioinformatics 9, no. 1 (2008): 1-13. | https://cran.r-project.org/web/packages/WGCNA/index.html |
| lme4 R package (version 1.1-27.1) | Bates, Douglas, Deepayan Sarkar, Maintainer Douglas Bates, and L. Matrix. "The lme4 package." R package version 2, no. 1 (2007): 74 | https://cran.r-project.org/web/packages/lme4/index.html |
| ordinal R package (version 2019.12-10) | Christensen, Rune Haubo B. "Cumulative link models for ordinal regression with the R package ordinal." Submitted in J. Stat. Software 35 (2018). | https://cran.r-project.org/web/packages/ordinal/index.html |
| gamm4 R package (version 0.2-6) | Wood, Simon, Fabian Scheipl, and Maintainer Simon Wood. "Package ‘gamm4’." Am Stat 45, no. 339 (2017): 0-2. | https://cran.r-project.org/web/packages/gamm4/index.html |
| ComplexHeatmap R package (version 2.6.2) | Gu Z, Eils R, Schlesner M (2016). “Complex heatmaps reveal patterns and correlations in multidimensional genomic data.” Bioinformatics. | https://www.bioconductor.org/packages/release/bioc/html/ComplexHeatmap.html |
| circlize R package (version 0.4.16) | Gu, Z. circlize implements and enhances circular visualization in R. Bioinformatics 2014. | https://cran.r-project.org/web/packages/circlize/index.html |
| pvca R package (version 1.30.0) | Bushel P (2021). pvca: Principal Variance Component Analysis (PVCA). R package version 1.34.0. | https://www.bioconductor.org/packages/release/bioc/html/pvca.html |

| | | |
|---|---|---|
| clusterProfiler R package (version 3.18.0) | Guangchuang Yu, Li-Gen Wang, Yanyan Han and Qing-Yu He. clusterProfiler: an R package for comparing biological themes among gene clusters. OMICS: A Journal of Integrative Biology 2012, 16(5):284-287 | https://bioconductor.org/packages/release/bioc/html/clusterProfiler.html |
| Msigdbr R package (version 7.5.1) | Igor Dolgalev (2022). msigdbr: MSigDB Gene Sets for Multiple Organisms in a Tidy Data Format. R package version 7.5.1. | https://igordot.github.io/msigdbr/ |
| ggbeeswarm R package (version 0.6.0) | Erik Clarke and Scott Sherrill-Mix (2017). ggbeeswarm: Categorical Scatter (Violin Point) Plots. R package version 0.6.0. | https://github.com/eclarke/ggbeeswarm |
| ggpubr R package (version 0.4.0) | Alboukadel Kassambara (2020). ggpubr: 'ggplot2' Based Publication Ready Plots. R package version 0.4.0. | https://rpkgs.datanovia.com/ggpubr/ |
| ggeffects R package (version 1.1.1) | Lüdecke D (2018). “ggeffects: Tidy Data Frames of Marginal Effects from Regression Models.” _Journal of Open Source Software_,*3*(26), 772. doi: 10.21105/joss.00772 (URL: https://doi.org/10.21105/joss.00772). | https://cran.r-project.org/web/packages/ggeffects/index.html |

| | | |
|---|---|---|
| Tidyverse R package (version 1.3.2) | Wickham H, Averick M, Bryan J, Chang W, McGowan LD, François R, Grolemund G, Hayes A, Henry L, Hester J, Kuhn M, Pedersen TL, Miller, E, Bache SM, Müller K, Ooms J, Robinson D, Seidel DP, Spinu V, Takahashi K, Vaughan D, Wilke C, Woo K, Yutani H (2019). “Welcome to the tidyverse.” _Journal of Open Source Software_, *4*(43), 1686. doi: 10.21105/joss.01686 (URL: https://doi.org/10.21105/joss.01686). | https://doi.org/10.21105/joss.01686 |
| mHG R package (version 1.1) | Eden, E. (2007). Discovering Motifs in Ranked Lists of DNA Sequences. Haifa. Retrieved from http://bioinfo.cs.technion.ac.il/people/zohar/thesis/eran.pdf | https://cran.r-project.org/web/packages/mHG/index.html |
| R6 R package (version 2.5.0) | https://github.com/r-lib/R6/ | https://cran.r-project.org/web/packages/R6/index.html |
| impute R package (version 1.64.0) | Hastie T, Tibshirani R, Narasimhan B, Chu G (2023). impute: impute: Imputation for microarray data. | https://bioconductor.org/packages/release/bioc/html/impute.html |
| limma R package (version 3.46.0) | Ritchie ME, Phipson B, Wu D, Hu Y, Law CW, Shi W, Smyth GK (2015). “limma powers differential expression analyses for RNA-sequencing and microarray studies.” Nucleic Acids Research, 43(7), e47 | http://bioconductor.org/packages/release/bioc/html/limma.html |
| boot R package (version1.3.28.1) | Canty A, Ripley BD (2022). boot: Bootstrap R (S-Plus) Functions. R package version 1.3-28.1. | https://cran.r-project.org/web/packages/boot/index.html |

| | | |
|---|---|---|
| ordinal R package (version 2022.11.16) | Christensen, R. H. B. (2022). ordinal - Regression Models for Ordinal Data. R package version 2022.11-16. https://CRAN.R-project.org/package=ordinal. | https://cran.r-project.org/web/packages/ordinal/index.html |
| ggalluvial R package (version 0.12.5) | Jason Cory Brunson and Quentin D. Read (2023). ggalluvial: Alluvial Plots in 'ggplot2'. R package version 0.12.5. http://corybrunson.github.io/ggalluvial/ | https://cran.r-project.org/web/packages/ggalluvial/index.html |
| Cytoscape (version 3.8.2) | Shannon P (2003) Cytoscape: a software environment for integrated models of biomolecular interaction networks *Genome Research* 13(11):2498-504 | cytoscape.org |
| BioRender | Biorender | biorender.com |
| SamTools bam2fq (v1.4, v1.2) | Danecek et al, 2021 | RRID:SCR_002105 |
| Trimmomatic-toolkit (v0.36.5) | Bolger, A. M., Lohse, M., & Usadel, B. (2014). Trimmomatic: A flexible trimmer for Illumina Sequence Data. *Bioinformatics*, btu170. | RRID:SCR_011848 |
| STAR aligner (v2.4.2a) | Dobin et al, Bioinformatics 2012 | RRID:SCR_004463 |
| HTSeq-count (v0.4.1) | Putri et al, 2021 | RRID:SCR_011867 |
| Picard (v1.134) | Broad Institute | RRID:SCR_006525 |
| FASTQC (v0.11.3) | Babraham Institute | RRID:SCR_014583 |
| Data.table R package 1.14.2 | Dowle, M, et al Data.table R package version 1.14.2 | https://cran.r-project.org/web/packages/data.table/index.html |
| DT R package 0.21 | Xue, Yihui, et al. DT: A Wrapper of the JavaScript Library DataTables R package version 0.21 | https://cran.r-project.org/web/packages/DT/index.html |

| | | |
|---|---|---|
| E1071 R package | Meyer, D, et al. e1071: Misc Functions of the Dept of Statistics, Probability Theory Group. R package version 1.7-9. | https://cran.r-project.org/web/packages/e1071/index.html |
| Metabolon Laboratory Information Management System (LIMS) | Metabolon | Metabolon |
| MassFragment Application Manager | Waters | Waters MassLynx v.4.1 Waters Corp Milford, USA |
| MetaboAnalyst 5.0 | MetaboAnalyst | https://www.metaboanalyst.ca/ |
| Cytutils R package v0.1.0 | Amir et al, 2017 | https://github.com/ismms-himc/cytutils |
| Fluidigm software-acquisition, normalization, concatenation v7.0.8493 | Fluidigm | https://www.fluidigm.com/products-services/software |
| Cytobank | Beckman Coulter | https://premium.cytobank.org |
| Prism 9 | GraphPad | https://www.graphpad.com/ |
| R v4.0.2 | The Comprehensive R Archive Network | https://cran.r-project.org/ |
| FLASH v1.2.11 | Magoc and Salzberg, 2011 | https://ccb.jhu.edu/software/FLASH/ |
| Bowtie2 v2.2.7 | Langmead and Salzberg, 2012 | http://bowtie-bio.sourceforge.net/bowtie2/index.shtml |
| Samtools v1.11 | Li et al., 2009 | http://samtools.sourceforge.net/ |
| NCBI BLAST v2.11.0 | Altschul et al., 1990 | https://blast.ncbi.nlm.nih.gov/Blast.cgi |
| CD-HIT | Li and Godzik, 2006 Fu et al., 2012 | http://weizhong-lab.ucsd.edu/cd-hit/download.php |
| COVID_pipe (https://github.com/mjsull/COVID_pipe) | mjsull, Gonzalez-Reiche, et al. 2020 | https://doi.org/10.5281/zenodo.3775031 |
| Minimap2 v2.17-r941 | Li, 2018 | https://doi.org/10.1093/bioinformatics/bty191 |
| Shovill v1.1.0 | Kwong, Gladman and Goncalves da Silva | https://github.com/tseemann/shovill |
| Pilon v1.24 | Walker et al. 2014 | http://doi.org/10.1371/journal.pone.0112963 |
| Canu v2.2 | Koren, et al. 2017 | http://doi.org/10.1101/gr.215087.116 |
| Prokka v1.14.6 | Seeman, 2014 | http://doi.org/10.1093/bioinformatics/btu153 |
| Seqkit v2.1.0 | Shen, et al. 2016 | http://doi.org/10.1371/journal.pone.0163962 |

| | | |
|---|---|---|
| Kraken2 v2.1.2 | Wood, et al. 2019 | https://doi.org/10.1186/s13059-019-1891-0 |
| Skyline v.21.2.1.377 | MacCossLab | http://skyline.ms |
| LabSolutions v.5.97 | Shimadzu Scientific Instruments | https://www.ssi.shimadzu.com/products/informatics/labsolutions.html |
| Perseus | Tyanova, et al. 2016 | https://maxquant.org/perseus// |
| Fluidigm Real-Time PCR Analysis v4.7.1 | Fluidigm | https://www.fluidigm.com/products-services/software |
| Olink NPX Manager v3.3.2.434 | Olink Proteomics | https://www.olink.com/products-services/data-analysis-products/npx-manager/ |
| Nextstrain v. 3.2.0 | Hadfield, et al. 2018 | https://github.com/nextstrain/ncov |
| Nextclade v. 1.11.0 | Aksamentov, et al. 2021 | https://doi.org/10.21105/joss.03773 |
| Pangolin v. 1.11.0 | O'Toole, et al. 2021 | https://doi.org/10.1093/ve/veab064 |
| Baltic v.0.1.6 | Dudas, 2016 | https://github.com/evogytis/baltic |
| IQ-TREE2 v.1.6.12 | Minh et al, 2020, Hoang et al 2018 | https://doi.org/10.1093/molbev/msaa015, https://doi.org/10.1093/molbev/msx281 |
| Other | | |
| Turbovap Evaporator | Biotage | Zymark TurboVap Cat#Z-TLVE |
| Waters Acquity UPLC | Waters | Waters Acquity |
| BEH C18 columns | Waters | Waters Acquity 2.1 x100 mm, 1.7 um columns |
| Q-Exactive with Orbitrap mass analyzer | Thermo Scientific | Cat#IQLAAEGAAPFALGMBDK |
| HILIC columns | Waters UPLC | Waters UPLC BEH Amide 2.1 x 150 mm, 1.7 um |
| Hamilton MicroLab Star Liquid Handling Robotic System | Hamilton Company | https://www.hamiltoncompany.com/automated-liquid-handling/platforms/microlab-star |
| Geno/Grinder 2000 | SPEX Sample Prep | Geno/Grinder 2000 |
| NovaSeq 6000 | Illumina | N/A |
| 0.45μm filter plates | Arctic White | AWFP-F20022 |
| 1000 ul Pipette Tips | Opentrons | 991-00005 |
| 300 ul Pipette Tips | Opentrons | 991-00008 |
| 20 ul Pipette Tips | Opentrons | 999-00014 |

| | | |
|---|---|---|
| 10 ul Pipette Tips | Opentrons | 999-00014 |
| 20 ul Pipette Tips | Axygen | T-20-R-S |
| 200 ul Pipette Tips | Axygen | T-200-C-L-R-S |
| Sealing tape 96-well Plates | 4titude | 4ti-0581 |
| 25ml Reservoir | Argos | B3125-100 |
| 4-well Reservoir | Axygen | RES-MW4-HP |
| 12-well Reservoir | Axygen | RES16MC-12-N |
| 0.5 ml 96-well Plates | VWR | 76210-520 |
| 0.8 ml 96-well Plates | VWR | 76210-524 |
| MACROSpin C18 plates | The Nest Group Inc. | SNS SS18VL |
| EvoTip | Evosep | EV2008 |
| PepSep LC 8cm column | Pepsep | PSC-8-150-15-UHP-nC - 8 cm nanoConnect column |
| Shimadzu LC column | Shimadzu | 227-32100-02 |
| Captive Spray Emitter (ZDV) 20 μm | Bruker | 1865710 |
| Combitips® advanced, Eppendorf Quality™, 0.5 mL | Eppendorf | 0030089421 |
| Combitips® advanced, Eppendorf Quality™, 2.5 mL | Eppendorf | 0030089448 |
| Combitips® advanced, Eppendorf Quality™, 5 mL | Eppendorf | 0030089448 |
| Combitips® advanced, Eppendorf Quality™, 10 mL | Eppendorf | 0030089464 |
| EvoSep One | Evosep | EV-1000 |
| Thermomixer | Eppendorf | N/A |
| timsTOF Pro | Bruker Daltonik GmBH | N/A |
| Column Oven Sonation PRSO-V2 | Sonication lab solutions | PRSO-V2 |
| Nexera Mikros | Shimadzu Scientific Instruments | N/A |
| LCMS 8060 | Shimadzu Scientific Instruments | N/A |
| Fluidigm Dynamic Array 96.96 GE IFC | Fluidigm | Cat#BMK-M-96.96 |
| Fluidigm Ctril Line Fluid,150ul | Fluidigm | Cat#89000021 |
| Magnetic COOH Beads Region 34 | BioRad | Cat#MC10034-01 |
| Magnetic COOH Beads Region 43 | BioRad | Cat#MC10043-01 |
| Magnetic COOH Beads Region 63 | BioRad | Cat#MC10063-01 |
| Amine coupling kit | BioRad | Cat#171406001 |
